# CRISPRi screens in human astrocytes elucidate regulators of distinct inflammatory reactive states

**DOI:** 10.1101/2021.08.23.457400

**Authors:** Kun Leng, Indigo V.L. Rose, Hyosung Kim, Wenlong Xia, Wilber Romero-Fernandez, Brendan Rooney, Mark Koontz, Emmy Li, Yan Ao, Shinong Wang, Mitchell Krawczyk, Julia TCW, Alison Goate, Ye Zhang, Erik M. Ullian, Michael V. Sofroniew, Stephen P.J. Fancy, Matthew S. Schrag, Ethan S. Lippmann, Martin Kampmann

## Abstract

In response to central nervous system injury or disease, astrocytes become reactive, adopting context-dependent states and functional outputs. Certain inflammatory insults induce reactive astrocytes that lose homeostatic functions and gain harmful outputs through cellular pathways that are not fully understood. Here, we combined single-cell transcriptomics with CRISPRi screening in human iPSC-derived astrocytes to systematically interrogate inflammatory astrocyte reactivity. We found that autocrine-paracrine IL-6 and interferon signaling downstream of canonical NF-κB activation drove two distinct inflammatory reactive signatures – one promoted by and the other inhibited by STAT3. These signatures overlapped with those observed in other experimental contexts, including mouse models, and their markers were upregulated in the human brain in Alzheimer’s disease and hypoxic ischemic encephalopathy. Furthermore, we validated that these signatures were regulated by Stat3 *in vivo.* These results and the platform we established have the potential to guide the development of therapeutics to selectively modulate different aspects of inflammatory astrocyte reactivity.

## INTRODUCTION

Astrocytes perform critical homeostatic functions in the central nervous system (CNS), such as providing trophic support for neurons, regulating the formation and function of synapses, and maintaining the blood–brain barrier^1^. In CNS injury or disease, astrocytes respond to pathophysiological perturbations by becoming “reactive”, which is defined as the adoption of context-specific states with associated transcriptomic signatures and alterations in morphology and function^1, 2^.

Frequently, inflammatory processes play an important, if not central, role in the pathophysiology of CNS injuries, as seen in stroke^3^ and trauma^4^, and diseases such as multiple sclerosis^5^ and Alzheimer’s disease^6^. As an immunocompetent CNS cell type, astrocytes actively participate in inflammatory signaling cascades and interact with both microglia, the CNS resident immune cells, as well as infiltrating immune cells from the periphery^6, 7^. Proinflammatory cytokines, such as IL-1α, TNF, and C1q, induce reactive astrocytes that lose homeostatic functions while releasing factors that are potentially harmful in specific contexts^8–10^. From here on, we will refer to this form of astrocyte reactivity induced by IL-1α+TNF+C1q as belonging to the category of “inflammatory reactivity”, which we will use as an umbrella term that captures a potential multitude of context-specific inflammatory reactive astrocyte signatures.

Given that inflammatory astrocyte reactivity has been implicated in numerous neurodegenerative and neuroinflammatory diseases^8, 11^, in addition to being associated with normal aging^12^, it is an attractive target for therapeutic development. However, the cellular pathways that control inflammatory astrocyte reactivity are still not fully understood, partly due to limitations in experimental scalability. On the one hand, animal models have provided key insights into genes required for astrocyte reactivity^9, 13, 14^, but the throughput to test genetic perturbations *in vivo* is extremely limited. On the other hand, *in vitro* astrocyte culture is a powerful alternative, but the scalability of experiments is still limited by the necessity to isolate primary astrocytes or the long duration of the differentiation process (up to 6 months) required to generate mature human iPSC (hiPSC)-derived astrocytes^15^. Lastly, even though molecular profiling approaches such as RNA-seq have been used extensively to identify cellular pathways altered in inflammatory reactive astrocytes^9, 16^, these assays by themselves can only provide correlative information and cannot uncover causal pathways controlling inflammatory reactivity.

Here, we developed a scalable method to generate hiPSC-derived astrocytes that allowed us to harness the power of pooled CRISPRi screening^17^ to systematically identify cellular pathways controlling inflammatory astrocyte reactivity induced by IL-1α+TNF+C1q. Following up on the top hits from CRISPRi screens with single-cell transcriptomics, we found that autocrine-paracrine IL-6 and interferon signaling drove two distinct inflammatory reactive signatures – one promoted by and the other inhibited by STAT3. We found that the inflammatory reactive signatures we identified overlapped with those observed in other experimental contexts, both *in vitro* and *in vivo*. Furthermore, we validated the induction of these distinct inflammatory reactive signatures by lipopolysaccharide (LPS) *in vivo* and also their regulation by *Stat3*. Lastly, we performed immunostaining supporting the existence of these inflammatory reactive signatures in human brains in the context of Alzheimer’s disease and hypoxic-ischemic encephalopathy.

## RESULTS

### iPSC-derived astrocytes (iAstrocytes) perform canonical astrocyte functions and sufficiently model inflammatory reactivity

To generate hiPSC-derived astrocytes in a scalable manner, we modified the protocol from TCW *et al.*^18^ by inducing the expression of the gliogenic transcription factors NFIA and SOX9 during the differentiation process as previously published by Li *et al.*^19^ (see Methods). Our protocol resulted in astrocytes, which we will here on refer to as “iAstrocytes”, with increased expression of astrocyte markers (Fig. 1a-b) as well as astrocyte-specific genes in general (Extended Data Fig. 1a-b) compared to astrocytes generated using the original protocol from TCW *et al*.^18^ (“TCW astrocytes”). Compared to Li *et al.*^19^, our iAstrocyte protocol differs in the following aspects (see Methods): 1) using a splice isoform of *NFIA* that is more highly expressed in astrocytes (ENST00000403491 in this study vs. ENST00000371187 in Li *et al.*^19^; Extended Data Fig. 1c, 2) ensuring the purity of neural progenitor cells through sorting for CD133+/CD271-cells^20^, 3) culturing astrocyte precursors during the differentiation process in monolayer, which is more straightforward in terms of media changes and cell dissociation compared to spheroid culture in Li *et al.*^19^, and 4) using a commercially available astrocyte media (ScienCell Astrocyte Media) adapted from TCW*et al*.

**Fig. 1.**
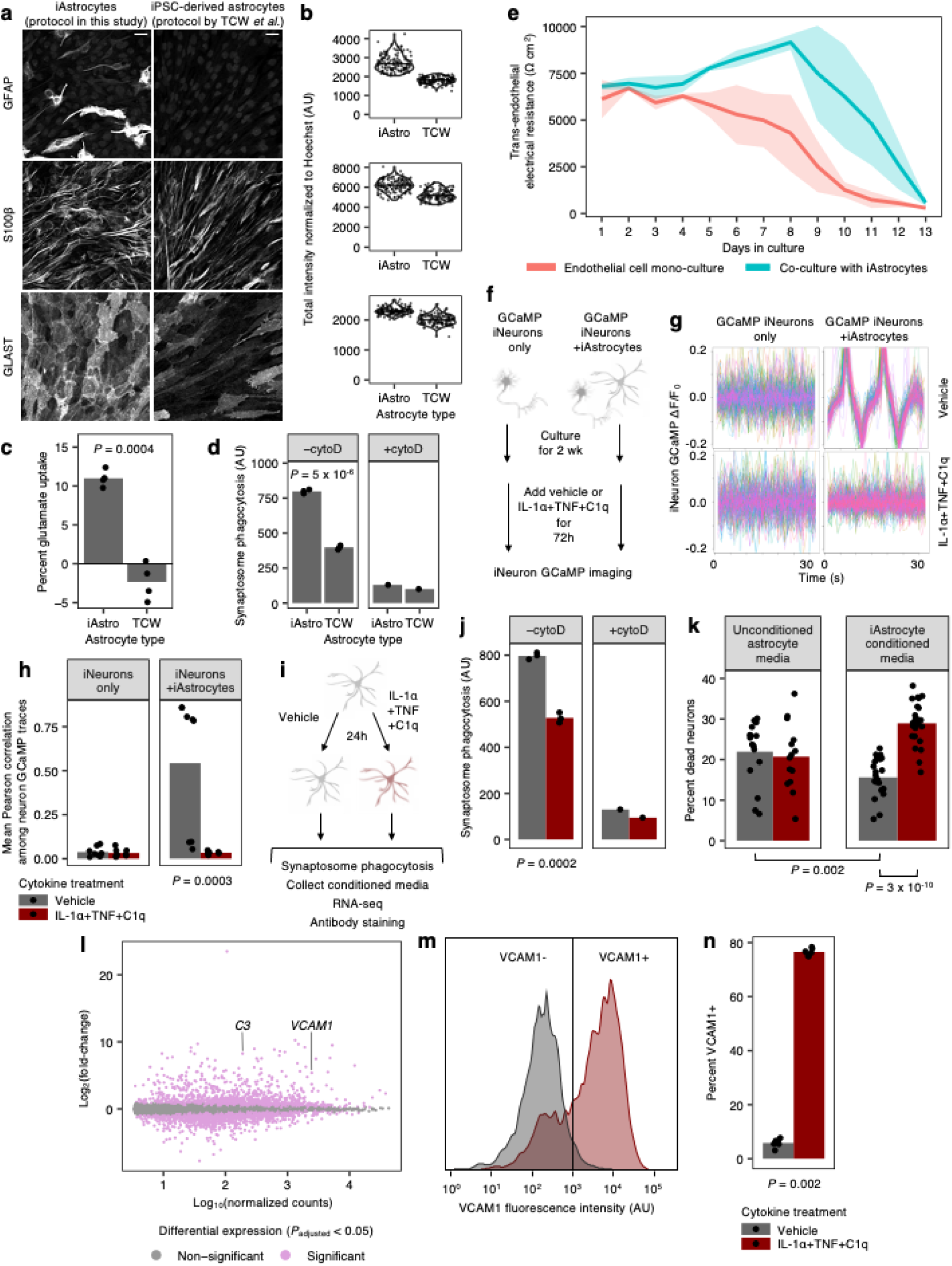
iPSC-derived astrocytes (iAstrocytes) perform canonical astrocyte functions and recapitulate key aspects of inflammatory reactivity upon treatment with IL-1α+TNF+C1q. **a**, Representative immunofluorescence micrographs of astrocyte marker expression in iAstrocytes (“iAstro”) vs. iPSC-derived astrocytes generated using the protocol from *TCW et al.*^18^ (“TCW”). Scale bar: 60 μm. **b**, Quantification of astrocyte marker expression measured by immunofluorescence (see Methods) in iAstrocytes vs. TCW astrocytes; data points represent fields of view collected over two experimental replicates. **c**, Glutamate uptake (see Methods) by iAstrocytes vs. TCW astrocytes (*n* = 4 wells). **d**, Phagocytosis of pHrodo-labeled rat synaptosomes (median pHrodo fluorescence intensity measured by flow cytometry) by iAstrocytes vs. TCW astrocytes in the absence (*n* = 3 wells) or presence (*n* = 1 wells) of cytochalasin D (cytoD). **e**, Barrier integrity of brain endothelial-like cells cultured alone or with iAstrocytes over time; lines represent group means (*n* = 5-6 wells), and shaded bands around lines represent 95% confidence intervals. **f**, Workflow schematic of calcium imaging experiments using GCaMP-expressing iPSC-derived neurons (iNeurons) cultured alone or with iAstrocytes. **g**, Neuronal calcium activity traces of GCaMP iNeurons co-cultured with iAstrocytes treated with vehicle control or IL-1α+TNF+C1q; traces from individual neurons are colored differently and overlaid. **h**, Quantification of synchrony between neuronal calcium activity traces in iNeuron mono-cultures or iNeuron + iAstrocyte co-cultures treated with vehicle control or IL-1α+TNF+C1q (*n* = 8 wells). **i**, Workflow schematic of experiments assessing inflammatory reactivity induced by IL-1α+TNF+C1q in iAstrocytes. **j**, Phagocytosis of pHrodo-labeled rat synaptosomes (median pHrodo fluorescence intensity measured by flow cytometry) by iAstrocytes treated with vehicle control or IL-1α+TNF+C1q in the absence (*n* = 3 wells) or presence (*n* = 1 wells) of cytoD. **k**, Percentage of dead cells (measured by TO-PRO-3 permeability) for iNeurons incubated with unconditioned astrocyte media +/- IL-1α+TNF+C1q or astrocyte media conditioned by iAstrocytes treated with vehicle control or IL-1α+TNF+C1q (*n* = 23-24 wells). **l**, Log-scaled fold change vs. average expression of differentially expressed genes (measured by RNA-seq) induced by IL-1α+TNF+C1q in iAstrocytes (*n* = 3 wells). **m**, Representative histogram of cell-surface VCAM1 levels (measured by flow cytometry) in iAstrocytes treated with vehicle control or IL-1α+TNF+C1q. **n**, Percent of VCAM1+ iAstrocytes (measured by flow cytometry) after treatment with vehicle control or IL-1α+TNF+C1q (*n* = 3 wells). In panels **c**, **h**, **k,** and n, *P* values were calculated using the two-sided Mann-Whitney U-test. In panels **d** and **j**, *P* values were calculated using the two-sided Student’s t-test.

We validated our iAstrocyte protocol in a total of three hiPSC lines derived from independent donors of both male and female sex (Extended Data Fig. 2a-b). To confirm that iAstrocytes were capable of performing typical astrocyte functions such as recycling glutamate^21^ and phagocytosing synapses^22^, we measured the ability of iAstrocytes to uptake glutamate or phagocytose pHrodo-labeled synaptosomes. We found that iAstrocytes had higher glutamate uptake activity (Fig. 1c) and synaptosome phagocytic activity (Fig. 1d) compared to TCW astrocytes. Furthermore, to see if iAstrocytes were capable of performing other canonical astrocyte functions such as maintaining the blood-brain barrier^23^ or promoting neuronal synapse maturation^24^, we co-cultured iAstrocytes with hiPSC-derived brain endothelial-like cells or neurons (see Methods). We found that brain endothelial-like cells co-cultured with iAstrocytes displayed improved barrier formation and integrity compared to mono-culture (Fig. 1e), demonstrating that iAstrocytes can promote the expected functional maturation of these cells. In addition, hiPSC-derived neurons (iNeurons) co-cultured with iAstrocytes developed synchronized calcium oscillations^25, 26^ compared to iNeurons in mono-culture (Fig. 1f-h), demonstrating the ability of iAstrocytes to promote neuronal synapse maturation and consistent with the high expression of genes encoding synapse-promoting proteins such as GPC4 ^27^ and CADM1 ^28^ in iAstrocytes (Supplementary Table 1). Furthermore, we found that iAstrocytes co-cultured with iNeurons acquired more stellate morphology compared to iAstrocytes in mono-culture (Extended Data Fig. 1d).

Having validated that iAstrocytes performed typical astrocyte functions, we next tested whether iAstrocytes could be used to model inflammatory astrocyte reactivity. It has been shown that the cytokines IL-1α+TNF+C1q induce a form of inflammatory reactivity associated with neurotoxicity and loss of homeostatic functions in both primary mouse astrocytes^8^ and human cerebral organoid-derived astrocytes^29^. We found that treatment of iAstrocytes generated from multiple hiPSC lines with IL-1α+TNF+C1q recapitulated previously reported *in vitro* phenotypes such as decreased phagocytosis of CNS substrates, decreased support of neuronal synapse maturation, and neurotoxicity^8, 29^. Specifically, iAstrocytes displayed decreased phagocytosis of pHrodo-labeled synaptosomes after treatment with IL-1α+TNF+C1q (Fig. 1i-j, Extended Data Fig. 2c). In addition, IL-1α+TNF+C1q treatment of iAstrocytes co-cultured with iNeurons abolished the development of synchronized neuronal calcium oscillations (Fig. 1f-h), demonstrating decreased support of neuronal synapse maturation by iAstrocytes. Finally, we found that the conditioned media from IL-1α+TNF+C1q-treated iAstrocytes was toxic to iNeurons, whereas the conditioned media from vehicle control-treated iAstrocytes was modestly protective compared to unconditioned media (Fig. 1k, Extended Data Fig. 2e).

In addition to recapitulating key functional phenotypes of IL-1α+TNF+C1q-induced inflammatory reactivity, iAstrocytes also responded to IL-1α+TNF+C1q at the transcriptomic level in a similar manner as primary mouse astrocytes and human iPSC-derived astrocytes generated using alternative protocols (Extended Data Fig. 3a-e, Supplementary Table 1). We noted, however, that the differentially expressed genes (DEGs) induced by IL-1α+TNF+C1q in hiPSC-derived astrocytes were not restricted to the previously defined “pan-reactive” and “A1” reactive categories (Extended Data Fig. 3c), which were derived from transcriptomic characterization of astrocytes from LPS-treated mice^8, 16^. This finding mirrored similar observations from Barbar *et al.*^29^. The discrepancy in behavior between hiPSC-derived astrocytes and mouse astrocytes could be due to differences in the experimental conditions or phylogenetic differences between human and mouse astrocytes^30^.

To find a cell-surface marker for inflammatory reactivity that is also functionally involved in neuroinflammation, we examined the DEGs induced by IL-1α+TNF+C1q in iAstrocytes. We found a dramatic increase in the transcript levels of *VCAM1* (Fig. 1l), which encodes a cell-adhesion sialoglycoprotein known to facilitate the infiltration of peripheral immune cells into the central nervous system (CNS) during neuroinflammation^31, 32^. We subsequently validated by flow cytometry that iAstrocytes generated from multiple hiPSC lines indeed upregulated cell-surface VCAM1 after treatment with IL-1α+TNF+C1q (Fig. 1m-n, Extended Data Fig. 2d), corroborating previous reports in the literature demonstrating induction of VCAM1 in astrocytes under pro-inflammatory conditions^33–35^.

### iAstrocytes differentiated from CRISPRi hiPSCs maintain robust knockdown activity

In order to investigate the effect of genetic perturbations on inflammatory reactivity in iAstrocytes, we generated iAstrocytes from hiPSCs stably expressing CRISPRi machinery (Fig. 2a), which enables both specific and non-toxic knockdown of genes targeted by single guide RNAs (sgRNAs) delivered to the cell and large-scale genetic screens^36^. We found that these iAstrocytes maintained robust CRISPRi activity, demonstrating ∼100% knockdown of TFRC protein levels 6 days after lentiviral transduction with a single guide RNA (sgRNA) targeting *TFRC* compared to iAstrocytes transduced with a non-targeting control (NTC) sgRNA, regardless of whether the iAstrocytes were treated with IL-1α+TNF+C1q (Fig. 2b-c).

**Fig. 2.**
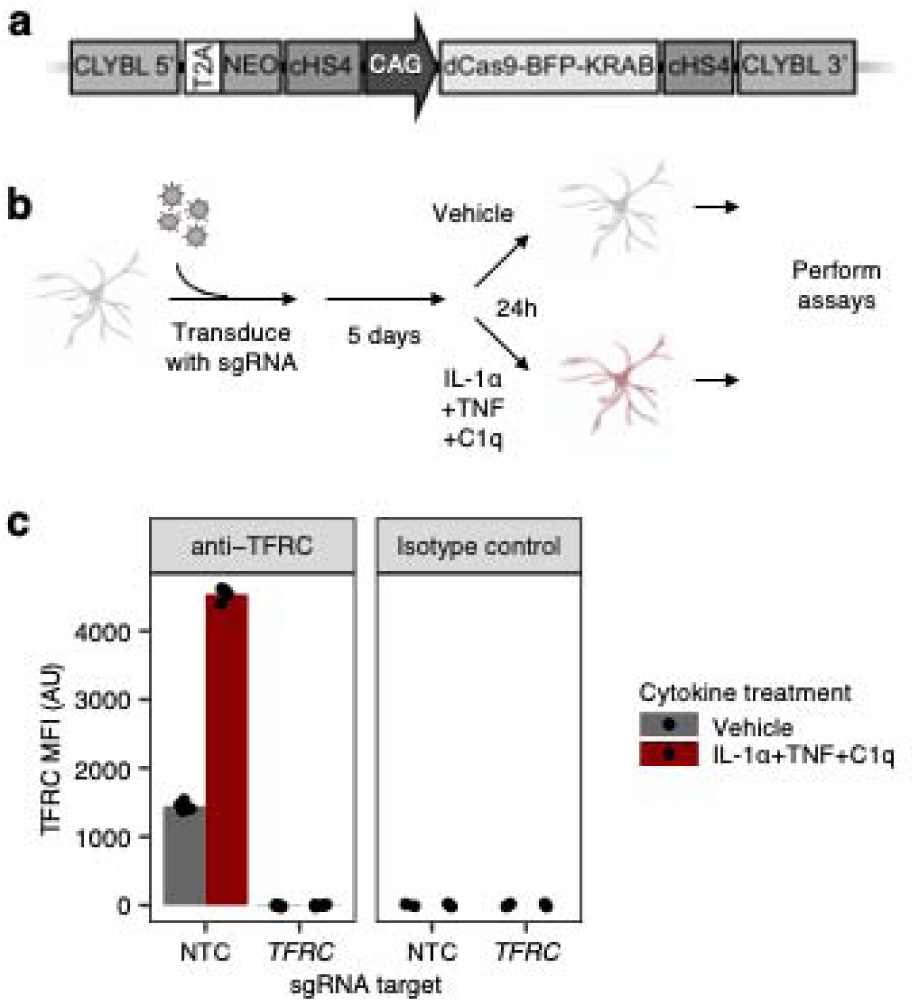
CRISPR interference (CRISPRi) platform in iAstrocytes. **a**, Schematic of CRISPRi machinery cassette integrated into the CLYBL safe-harbor locus. **b**, Workflow schematic of experiments involving lentiviral sgRNA transduction of iAstrocytes for CRISPRi knockdown. **c**, Cell-surface TFRC levels (measured by flow cytometry using an anti-TFRC antibody or isotype control; MFI: median fluorescence intensity) in iAstrocytes transduced with a non-targeting control (NTC) sgRNA or a sgRNA targeting *TFRC*, treated with vehicle control or IL-1α+TNF+C1q (*n* = 4 wells for anti-TFRC, *n* = 2 wells for isotype control).

### CRISPRi screening and computational master regulator analysis uncover cellular pathways controlling inflammatory reactivity

Having established CRISPRi in iAstrocytes, we aimed to systematically interrogate IL-1α+TNF+C1q-induced inflammatory reactivity in iAstrocytes with two orthogonal approaches – pooled CRISPRi screening^17, 37^ and computational master regulator analysis (MRA)^38–40^.

For pooled CRISPRi screening, we used the same experimental timeline as described in Fig. 2b. We conducted screens based on two cell-autonomous phenotypes induced by IL-1α+TNF+C1q that can be analyzed by fluorescence activated cell sorting (FACS): decreased phagocytosis of pHrodo-labeled synaptosomes (Fig. 1i-j) and upregulation of cell-surface of VCAM1 (Fig. 1m-n). For synaptosome phagocytosis, we performed the screens in both vehicle control-treated and IL-1α+TNF+C1q-treated iAstrocytes to find genes whose knockdown could rescue or exacerbate the decrease in synaptosome phagocytosis induced by IL-1α+TNF+C1q (Fig. 3a). For upregulation of cell-surface VCAM1, we performed the screens in only IL-1α+TNF+C1q-treated iAstrocytes (Fig. 3a) given that in the absence of cytokine treatment, very few iAstrocytes expressed cell-surface VCAM1 (Fig. 1m). To ensure that the results from the screens are generalizable, we validated that IL-1α+TNF+C1q also caused decreased synaptosome phagocytosis and upregulation of cell-surface VCAM1 in hiPSC-derived astrocytes generated using alternative protocols (Extended Data Fig. 3f-h), as well as upregulation of cell-surface VCAM1 in primary mouse astrocytes (Extended Data Fig. h-i).

**Fig. 3.**
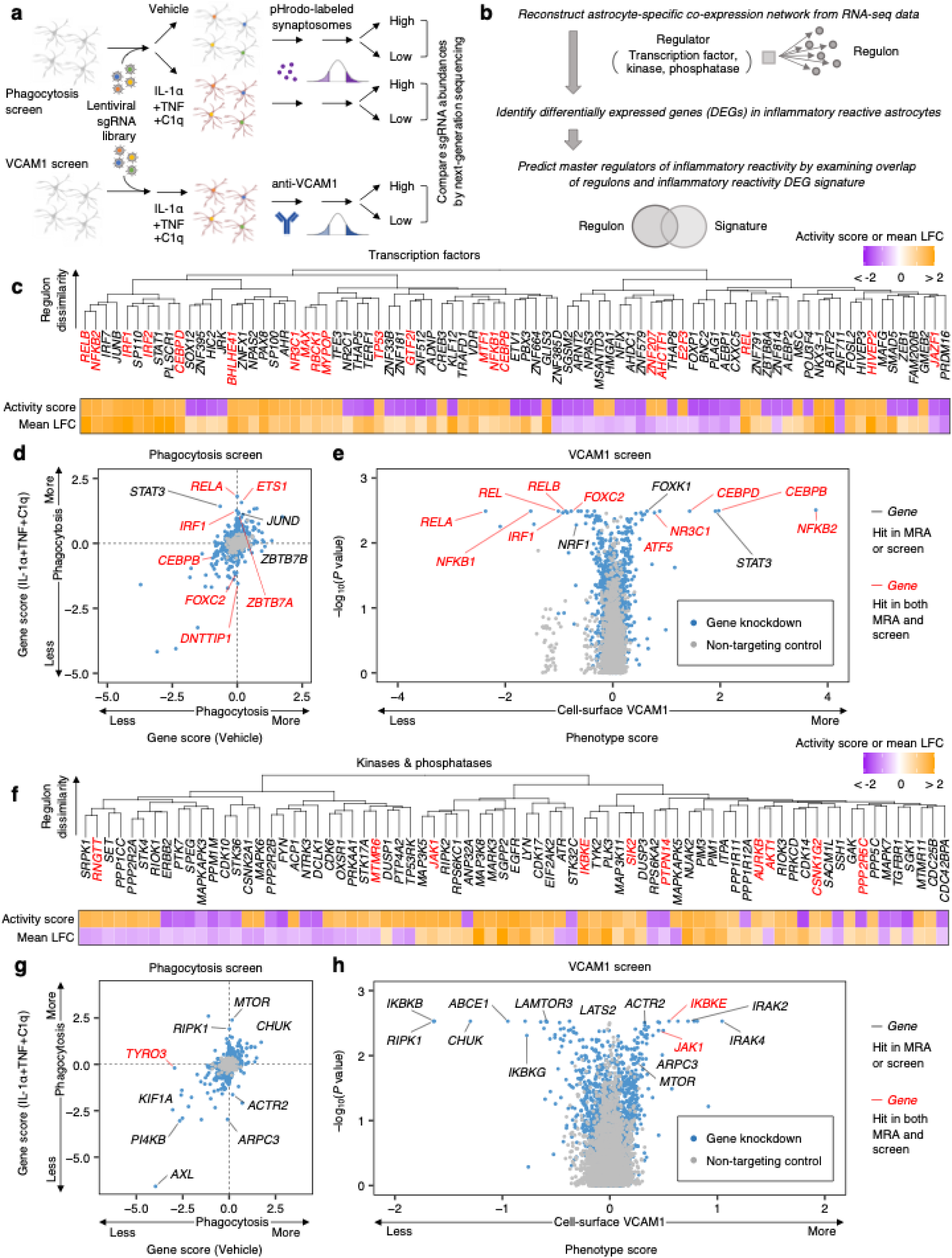
CRISPRi screening and master regulator analysis uncover regulators of inflammatory reactivity. **a**, Workflow schematic of synaptosome phagocytosis and cell-surface VCAM1 CRISPRi screens (*n* = 2 independent screens). **b**, Workflow schematic of bioinformatic master regulator analysis (MRA). **c**, **f**, Clustering of transcription factors (**c**) or kinases and phosphatases (**f**) predicted to regulate inflammatory reactivity based on regulon overlap (see Methods); the activity score and regulon mean log-fold-change (LFC) associated with each predicted regulator (see Methods) are shown below the dendrogram. **d**, **g**, Scatterplot of gene scores (see Methods) of transcription factors (**d**) or the druggable genome (**g**) from synaptosome phagocytosis screens on iAstrocytes treated with vehicle control vs. IL-1α+TNF+C1q. **e**, **h**, Volcano plot of phenotype scores and associated log-scaled *P* values (see Methods) of transcription factors (**e**) or the druggable genome (**h**) from cell-surface VCAM1 screens on iAstrocytes treated with IL-1α+TNF+C1q.

For MRA, the workflow consisted of first reconstructing a gene regulatory network between upstream regulators (e.g. transcription factors, or kinases and phosphatases) and target genes in human astrocytes based on coexpression, which required integrating a large number of human astrocyte expression profiles collected under diverse experimental conditions^41^ (see Methods); upstream regulators that control inflammatory reactivity were then predicted by examining the overlap between the regulons of upstream regulators with the DEGs induced by IL-1α+TNF+C1q in iAstrocytes (Fig. 3b).

We performed MRA using the human transcription factors^42^ (Fig. 3c) or kinases and phosphatases^43, 44^ (Fig. 3f) as upstream regulators, and similarly conducted pooled CRISPRi screening using a new custom sgRNA library targeting the human transcription factors (“TF library; Fig. 3d-e) and a library targeting all kinases, phosphatases, and other genes representing the “druggable genome”^45^ (“H1 library”; Fig. 3g-h). Together, these two approaches uncovered both expected regulators of inflammatory reactivity as well as potentially novel regulators.

For example, the top hits from both MRA and CRISPRi screens (Supplementary Tables 2-3) included the entire set of genes encoding all potential subunits of the NF-κB transcription factor (*REL*, *RELA*, *RELB*, *NFKB1*, *NFKB2*; Fig. 3c-e) as well as genes encoding the upstream kinases of the canonical NF-κB pathway (*IKBKB*, *CHUK*, *IKBKG*; Fig. 3f-h)^46^. For all the above genes (with the exception of *NFKB2*), the directionality of their screening phenotypes or master regulator analysis activity scores (see Methods) was consistent with the activation of the canonical NF-kB pathway being required for the induction of inflammatory reactivity. As an example, knockdown of *RELA*, which encodes the p65 subunit of the NF-κB transcription factor, blocked the induction of cell-surface VCAM1 (Fig. 3e) and rescued the decrease in phagocytosis (Fig. 3d) induced by IL-1α+TNF+C1q (also see Extended Data Fig. 4b). In addition, the activity score of *RELA* from MRA was positive, consistent with the positive average log-fold change of DEGs under the control of *RELA* (Supplementary Tables 2-3). For the case of *NFKB2*, knockdown of which actually increased VCAM1 induction by IL-1α+TNF+C1q (Fig. 3e), it is possible that the encoded protein, p100, may act to inhibit *RELA*-dependent transcription through the cytoplasmic sequestration of p65 ^47^.

Given that the canonical NF-κB pathway is a well-studied master regulator of cellular responses to inflammatory stimuli^48^, its strong enrichment in the top hits from the screens and MRA serves as a positive control for the technical quality of the screens (see Extended Data Fig. 4a for quality control data) and the validity of the MRA pipeline. As a further control for the technical quality of the screens, we validated the phenotypes of selected top hits in independent experiments (Extended Data Fig. 4c-d).

In addition to the canonical NF-κB pathway, we also recovered genes involved in numerous other cellular pathways known to mediate inflammatory processes. One group of genes (*STAT3*, *CEBPB*, *CEBPD*) consisted of transcription factors classically associated with the acute phase response in the context of systemic inflammation^49, 50^. In the central nervous system, these transcription factors control similar responses in reactive astrocytes, such as the production of acute phase proteins C3 and α1-antitrypsin (*SERPINA1*) by astrocytes challenged with IL-1 and TNF^49^ (also see Fig. 1l and Supplementary Table 1). Furthermore, *STAT3* is required for the full induction of astrocyte reactivity caused by spinal cord injury^13, 51, 52^, and *CEBPB* has been implicated in the pathogenesis of Alzheimer’s disease^53, 54^. We found that knockdown of *STAT3*, *CEBPB*, and *CEBPD* in the context of IL-1α+TNF+C1q treatment surprisingly promoted the induction VCAM1 while having mixed effects on phagocytosis (Fig. 3d,e; Extended Data Fig. 4b).

Another group of genes consisted of those involved in the response to interferons (*IRF1*, *STAT1*, *IKBKE*)^55–57^. For example, we found that knockdown of *IRF1* rescued decreased phagocytosis and reduced VCAM1 upregulation (Fig. 3d,e,h; Extended Data Fig. 4b). Given that interferons have been shown to potentiate the induction of VCAM1 by inflammatory cytokines^33, 58^, the reduction of VCAM1 upregulation with *IRF1* knockdown suggests that interferons are released by iAstrocytes after IL-1α+TNF+C1q treatment.

The release of interferons by iAstrocytes after IL-1α+TNF+C1q treatment would also explain the effect we observed with knockdown of genes encoding kinases responsible for signal transduction from the IL-1 receptor (*IRAK2, IRAK4*)^59^. Knockdown of *IRAK2* or *IRAK4*, which should block signaling from the IL-1 receptor, strongly increased VCAM1 induction (Fig. 3h). This can be explained by the fact that IL-1 signaling is known to antagonize the action of interferons^59^.

Lastly, we also found genes involved in cellular pathways that have been relatively less studied in the context of inflammatory astrocyte reactivity. For example, we uncovered genes involved in or related to the mTOR pathway (*MTOR*, *LAMTOR3*, *LATS2*, *FOXK1*)^60, 61^, the glucocorticoid receptor pathway (*NR3C1*), the actin cytoskeleton (*ARPC3*, *ACTR2*), and also relatively uncharacterized genes such as *FOXC2* and *ZBTB7A*.

Overall, the results from CRISPRi screening and MRA demonstrate that numerous cellular pathways regulate distinct aspects of inflammatory reactivity induced by IL-1α+TNF+C1q. Of particular interest to us was the fact that knockdown of the acute phase response-related genes *STAT3*, *CEBPB*, or *CEBPD* increased the induction of VCAM1 by IL-1α+TNF+C1q, whereas knockdown of the interferon-related gene *IRF1* decreased VCAM1 induction. Since all of the above genes are known to be involved in inflammatory responses, their opposing phenotypes suggested to us that VCAM1 upregulation may only be capturing one particular aspect of IL-1α+TNF+C1q-induced inflammatory reactivity, i.e. that IL-1α+TNF+C1q may induce distinct inflammatory reactive signatures, one of which is marked by VCAM1 upregulation.

### CROP-seq reveals two distinct inflammatory reactive signatures

To gain deeper insight into how regulators uncovered by CRISPRi screening and MRA may control distinct inflammatory reactive signatures, we turned to CROP-seq^62^. By coupling CRISPRi perturbations to single-cell transcriptomics, CROP-seq enables the recovery of perturbation-associated changes in gene expression from a pooled experiment. We selected 30 regulators (Extended Data Fig. 5d; Supplementary Table 6) that were strong hits from CRISPRi screening and MRA for pooled knockdown in iAstrocytes treated with vehicle control or IL-1α+TNF+C1q, and then performed CROP-seq to characterize the effect of knockdown on IL-1α+TNF+C1q-induced gene expression (see Methods).

Before examining the effect of knocking down different regulators on inflammatory reactivity, we first focused on iAstrocytes transduced with non-targeting control (NTC) sgRNAs. In the absence of IL-1α+TNF+C1q, NTC iAstrocytes partitioned largely by cell cycle in uniform manifold approximation projection (UMAP), with cluster 3 corresponding to dividing cells. A small fraction (<10%) of endothelial-like and stromal-like cells were also present, corresponding to clusters 5 and 6, respectively (Fig. 4a, Extended Data Fig. 5a-c, Supplementary Table 4). Upon treatment with IL-1α+TNF+C1q, NTC iAstrocytes partitioned into two distinct clusters (clusters 1 and 2), with an additional small cluster (cluster 4) corresponding to cycling cells (Fig. 4a, Extended Data Fig. 5a-c, Supplementary Table 4).

**Fig. 4.**
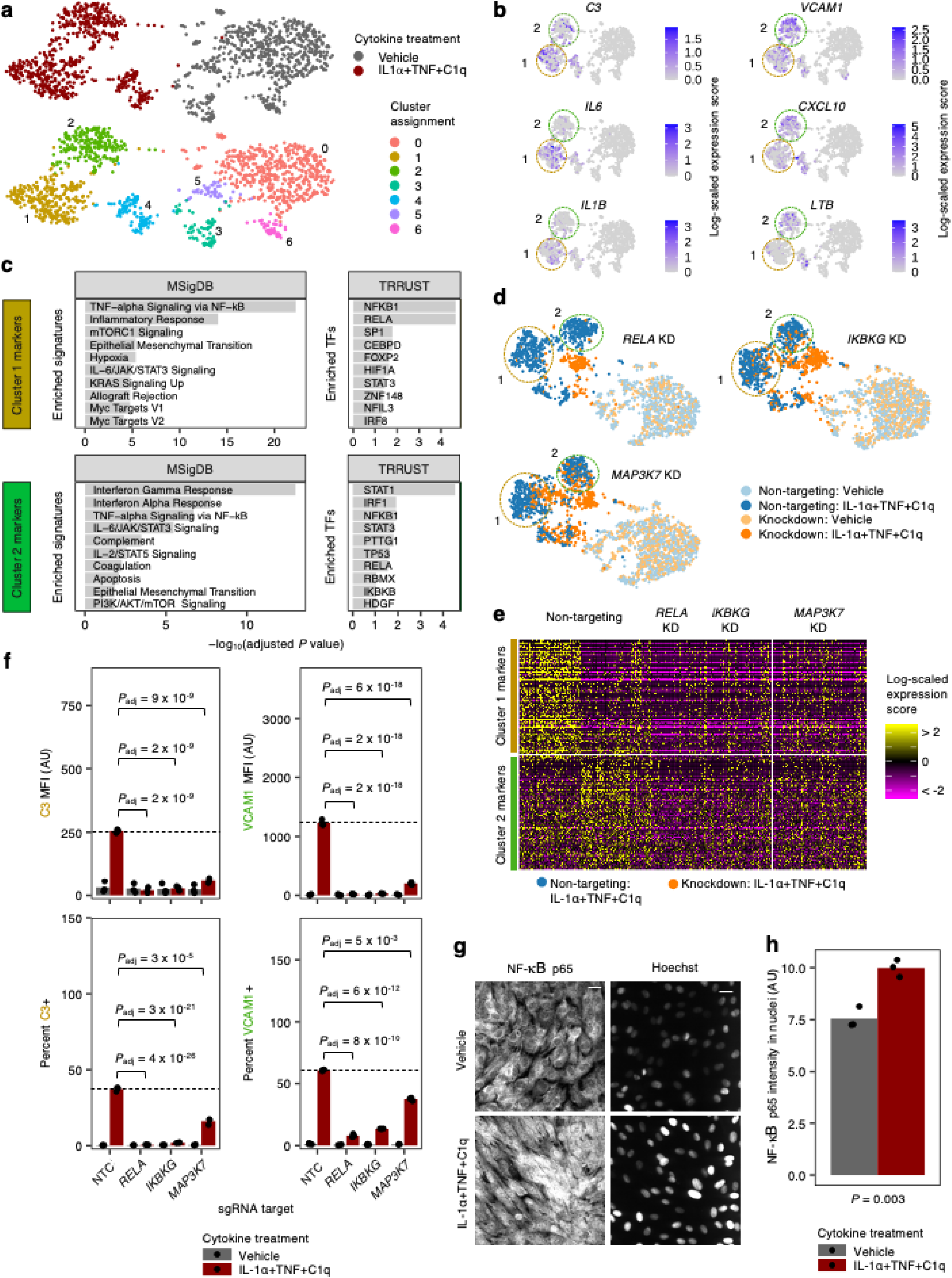
CROP-seq of iAstrocytes reveals two distinct inflammatory reactive signatures dependent on the canonical NF-kB pathway. **a**, Uniform manifold approximation projection (UMAP) of single-cell transcriptomes of iAstrocytes transduced with a non-targeting control (NTC) sgRNA treated with vehicle control or IL-1α+TNF+C1q, colored by cytokine treatment or cluster assignment. **b**, Visualization of transcript levels of selected Cluster 1 and Cluster 2 markers in the iAstrocytes shown in panel **a**, overlaid onto the same UMAP embedding. **c**, Cellular pathway (MSigDB^131^) and upstream transcription factor (TRRUST^132^) enrichment analysis of Cluster 1 and Cluster 2 markers; TF – transcription factor. **d**, Aligned UMAP embedding (see Methods) of NTC sgRNA transduced iAstrocytes with iAstrocytes transduced with sgRNAs knocking down *RELA*, *IKBKG*, or *MAP3K7*. **e**, Heatmap of Cluster 1 and Cluster 2 marker transcript levels in IL-1α+TNF+C1q-treated NTC sgRNA iAstrocytes compared to IL-1α+TNF+C1q-treated *RELA*, *IKBKG*, or *MAP3K7* sgRNA iAstrocytes. **f**, VCAM1/C3 levels (MFI: median fluorescence intensity) or percent positive cells measured by flow cytometry in NTC, *RELA*, *IKBKG*, or *MAP3K7* sgRNA iAstrocytes treated with vehicle control or IL-1α+TNF+C1q (*n* = 3 wells). **g**, Representative images of NF-κB p65 immunostaining or Hoechst counterstain in iAstrocytes treated with vehicle control or IL-1α+TNF+C1q. Scale bar: 60 μm. **h**, Quantification of NF-κB p65 nuclear localization (integrated fluorescence intensity masked to Hoechst; *n =* 3 wells). In panel **c**, the adjusted *P* values were derived from Enrichr^123^. In panel **f**, *P* values were calculated using linear regression for MFI values or beta regression for percentages (see Methods) and adjusted for multiple testing (*P*_adj_; Holm’s method) per family of tests (all comparisons made within a plot). In panel **h**, the *P* value was calculated using the two-sided student’s t-test. In panels **b** and **d**, NTC sgRNA astrocytes in Cluster 1 or Cluster 2 are circled by colored dotted lines.

Upon further examination of the two major clusters induced by IL-1α+TNF+C1q, we found that iAstrocytes in cluster 1 expressed markers related to the acute phase response such as *C3*, *IL6*, and *IL1B*, whereas iAstrocytes in cluster 2 expressed markers related to interferon (IFN) signaling such as *VCAM1*, *CXCL10*, and *LTB*^63^ (Fig. 4b). Enrichment analysis of cluster 2 markers confirmed a strong, specific enrichment (i.e. unique to cluster 2 markers) for interferon-related pathways (e.g. “Interferon Gamma Response”, “Interferon Alpha Response”) and transcription factor regulons (e.g. STAT1, IRF1) (Fig. 4c, Supplementary Table 5). On the other hand, cluster 1 markers showed a specific enrichment for pathways and transcription factor regulons related to the mTORC1-HIF1α/MYC axis of metabolic control^64^ (e.g. “mTORC1 Signaling”, “Hypoxia”, “Myc Targets V1”; HIF1A, CEBPD^65^). For brevity, we will hereon provisionally refer to the inflammatory reactive astrocyte signature (IRAS) corresponding to cluster 1 as IRAS1 and the one corresponding to cluster 2 as IRAS2.

Given that the results from CRISPRi screening and MRA pointed towards the canonical NF-κB pathway being required for the induction of inflammatory reactivity, we examined the effect of knocking down *RELA* (NF-κB p65 subunit), *IKBKG* (NEMO), or *MAP3K7* (TAK1) on the induction of IRAS1 and IRAS2 by IL-1α+TNF+C1q in our CROP-seq data. Aligned UMAP embedding (see Methods) of *RELA*, *IKBKG*, or *MAP3K7* knockdown iAstrocytes with NTC iAstrocytes showed that *RELA* knockdown prevented the full induction of both IRAS1 and IRAS2, with *IKBKG* and *MAP3K7* knockdown showing similar but less complete effects (Fig. 4d). Examining the expression of IRAS1 and IRAS2 markers in IL-1α+TNF+C1q-treated NTC iAstrocytes vs. IL-1α+TNF+C1q-treated *RELA*, *IKBKG*, or *MAP3K7* knockdown iAstrocytes showed the same pattern (Fig. 4e). To validate the above findings in an independent experiment, we measured the induction of C3, a marker of IRAS1, and VCAM1, a marker of IRAS2, by flow cytometry in NTC iAstrocytes and *RELA*, *IKBKG*, or *MAP3K7* knockdown iAstrocytes treated with IL-1α+TNF+C1q; we found that *RELA*, *IKBKG*, or *MAP3K7* knockdown strongly reduced the induction of VCAM1 and C3 (Fig. 4f). Lastly, we performed immunostaining against NF-κB p65 and confirmed that IL-1α+TNF+C1q treatment induced the nuclear localization of p65 in iAstrocytes (Fig. 4g-h).

To gain a more global view of how knocking down different regulators affected the induction of inflammatory reactivity, we performed differential gene expression analysis for each regulator to find how regulator knockdown altered the differential expression induced by IL-1α+TNF+C1q (Extended Data Fig. 5d, Supplementary Table 6; see Methods). We then performed hierarchical clustering on the log-fold-changes (weighted by statistical significance) of the union of all DEGs (see Methods). We found that regulators segregated into three major groups, with one group (G1) consisting largely of regulators involved in the canonical NF-κB pathway (*RELA*, *IKBKG*, *NFKB1*, *REL*, *CHUK*) and upstream signal transduction (*IRAK4*, *RIPK1*, *MAP3K7*), another group (G2) that contained acute phase response-related transcription factors (*CEBPB*, *CEBPD*, *STAT3*), and the last group (G3) consisting of regulators with weak knockdown phenotypes (Extended Data Fig. 5e). In terms of coordinately regulated genes, we recovered 10 gene modules that displayed distinct patterns across knockdown of different regulators (Extended Data Fig. 5e, Extended Data Fig. 6, Supplementary Table 7).

Of particular interest were modules M3, M4, and M9, which all had reduced induction upon knockdown of G1 regulators and thus appeared to be downstream of the canonical NF-κB pathway. Interestingly, M4 genes contained and had similar enrichments as IRAS1 markers (e.g. *IL6*; see Supplementary Table 7 and Extended Data Fig. 6) and displayed reduced induction upon knockdown of G2 regulators (Extended Data Fig. 5e), whereas M3 and M9 genes contained and had similar enrichments as IRAS2 markers (e.g. *VCAM1*; see Supplementary Table 7 and Extended Data Fig. 6) and displayed increased induction upon knockdown of G2 regulators (Extended Data Fig. 5e). This led us to hypothesize that the acute phase response-related transcription factors *CEBPB*, *CEBPD*, and *STAT3* promote IRAS1 while inhibiting IRAS2.

### IL-6 and interferons act in an autocrine-paracrine manner to drive IRAS1 and IRAS2

Given that IRAS1 iAstrocytes expressed markers related to the acute phase response, which is driven by IL-6 ^66^, and that IRAS2 iAstrocytes expressed markers related to the response to interferons, we hypothesized that IL-1α+TNF+C1q induced iAstrocytes to secrete IL-6 and interferons, which then acted on the iAstrocytes in an autocrine-paracrine manner. We found that iAstrocytes derived from multiple hiPSC lines indeed secreted appreciable amounts of IL-6 (∼40,000 pg/mL) and IFN-β (∼1,000 pg/mL) in response to IL-1α+TNF+C1q (Fig. 5a, Extended Data Fig. 7d). iAstrocytes also secreted appreciable amounts of GM-CSF (∼3,500 pg/mL), which is known to act synergistically with IL-6 ^67^, as well as CXCL10 (∼30,000 pg/mL), which is produced in response to interferons^68^.

**Fig. 5.**
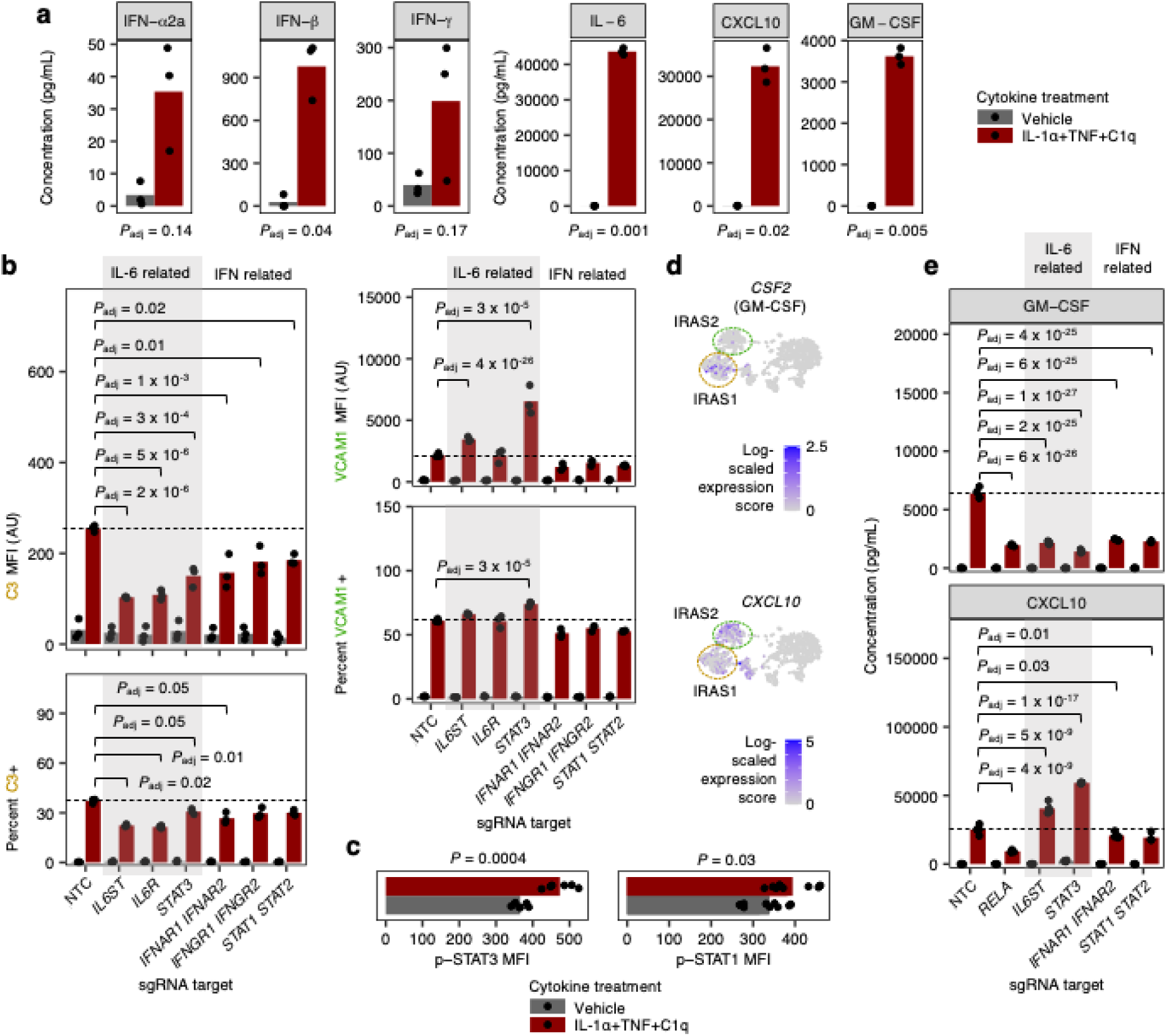
IL-6 and interferons act in an autocrine-paracrine manner downstream of IL-1α+TNF+C1q. **a**, Cytokine concentrations in conditioned media (measured by multi-spot electrochemiluminescence) from iAstrocytes treated with vehicle control or IL-1α+TNF+C1q (*n* = 3 wells). **b**, VCAM1/C3 levels (MFI: median fluorescence intensity) or percent positive cells measured by flow cytometry from NTC sgRNA iAstrocytes compared to iAstrocytes transduced with sgRNAs knocking down genes or gene pairs involved in IL-6 or interferon (IFN) signaling as indicated (*n* = 3 wells). **c**, Levels of phosphorylated STAT3 (Y705; p-STAT3) or phosphorylated STAT1 (Y701; p-STAT1) measured by flow cytometry in iAstrocytes treated with vehicle control or IL-1α+TNF+C1q (*n* = 6 wells for p-STAT3 and *n* = 9 wells for p-STAT1). **d**, Transcript levels of *CSF2* and *GM-CSF* overlaid onto the UMAP embedding from Fig. 3a. **e**, Concentration of GM-CSF or CXCL10 in conditioned media from iAstrocytes transduced with sgRNAs knocking down genes or gene pairs involved in IL-6 or IFN signaling as indicated (*n* = 4 wells). In panels **a** and **c**, *P* values were calculated using the two-sided Student’s t-test. In panels **b** and **e**, *P* values were calculated using linear regression for MFI values or beta regression for percentages (see Methods), and only comparisons with statistically significant differences are marked. Where appropriate, *P* values were adjusted for multiple testing (*P*_adj_; Holm’s method) per family of tests (all comparisons made within a plot).

To test whether the secreted IL-6 and interferons acted on the iAstrocytes in an autocrine-paracrine manner, we knocked down their receptors and downstream transcription factors and measured the induction of C3 and VCAM1, markers of IRAS1 and IRAS2 respectively, by flow cytometry. We found that knockdown of *IL6R* or *IL6ST*, which respectively encode the IL-6 receptor and its signal transducing partner gp130, decreased the induction of C3+ iAstrocytes; the induction of VCAM1 in iAstrocytes increased slightly for the case of *IL6ST* knockdown but did not change for *IL6R* knockdown (Fig. 5b). Knockdown of *STAT3*, which is activated downstream of IL-6 ^69^, similarly decreased the induction of C3+ iAstrocytes while increasing the induction of VCAM1+ iAstrocytes (Fig. 5b). We confirmed that IL-1α+TNF+C1q induced STAT3 activation as measured by increased STAT3 phosphorylation at Y705 (Fig. 5c).

On the other hand, concurrent knockdown of *IFNAR1* and *IFNAR2* (see Methods), which encode subunits of the type I interferon (IFN) receptor, or *IFNGR1* and *IFNGR2*, which encode subunits of the type II interferon receptor, decreased the induction of both C3+ iAstrocytes and VCAM1+ iAstrocytes (Fig. 5b). Concurrent knockdown of *STAT1* and *STAT2*, which are activated downstream of interferons^70^, also resulted in a decrease in the induction of both C3+ iAstrocytes and VCAM1+ iAstrocytes (Fig. 5b). We confirmed that IL-1α+TNF+C1q induced STAT1 activation as measured by increased STAT1 phosphorylation at Y701 (Fig. 5c).

As an alternative approach to assay the induction of IRAS1 and IRAS2, we assayed the production of GM-CSF and CXCL10, which should be produced by IRAS1 and IRAS2 iAstrocytes respectively based on mRNA expression (Fig. 5d). Knockdown of *IL6ST* or *STAT3* decreased the production of GM-CSF but increased the production of CXCL10, whereas knockdown of *IFNAR1/2* or *STAT1/2* decreased the production of both GM-CSF and CXCL10 (Fig. 5e).

A limitation of the above experiments is that only a single marker of IRAS1 (C3, GM-CSF) or IRAS2 (VCAM1, CXCL10) was examined at a time, even though these markers individually do not perfectly distinguish between IRAS1 and IRAS2 (Fig. 4b, Fig. 5c). To better resolve IRAS1 and IRAS2, we performed flow cytometry on iAstrocytes stained for VCAM1 and C3 simultaneously (see Methods). With IL-1α+TNF+C1q treatment, we were able to resolve VCAM1–/C3+, VCAM1+/C3–, and VCAM1+/C3+ iAstrocytes (Fig. 6a). Given the expression pattern of C3 and VCAM1 in our single-cell data (Fig. 4b), we mapped VCAM1–/C3+ iAstrocytes to IRAS1 and VCAM1+/C3– iAstrocytes to IRAS2. However, we were uncertain whether VCAM1+/C3+ iAstrocytes mapped to IRAS1 or IRAS2.

**Fig. 6.**
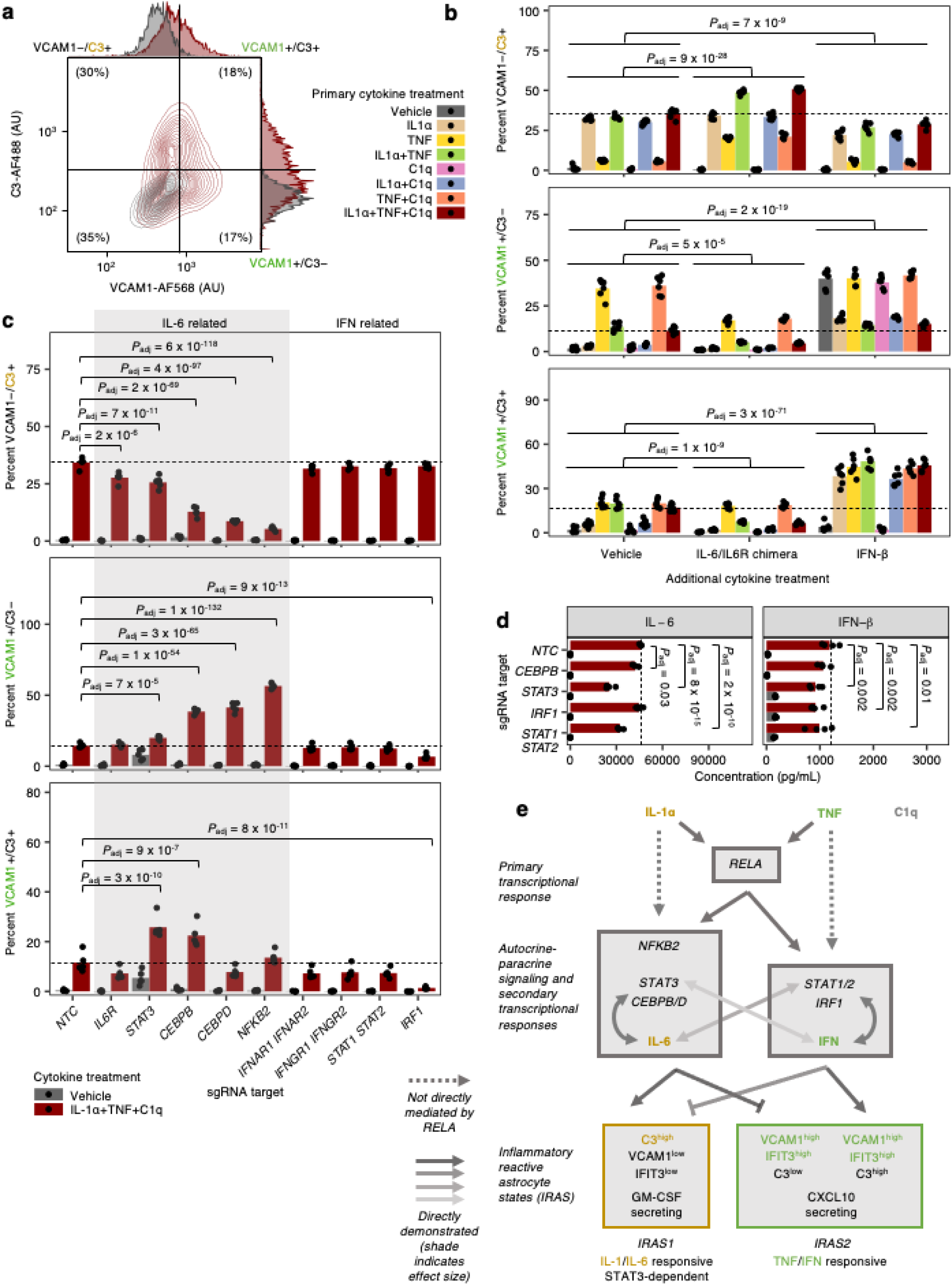
Differential regulation of distinct inflammatory reactive signatures by cytokines and cellular factors. **a**, Representative contour plot of VCAM1 and C3 levels measured by flow cytometry in iAstrocytes treated with vehicle control or IL-1α+TNF+C1q. **b**, Percent VCAM1−/C3+, VCAM1+/C3−, or VCAM1+/C3+ cells measured by flow cytometry in iAstrocytes treated with vehicle control vs. all possible combinations of IL-1α, TNF, and C1q, in the absence or presence of additional IL-6/IL6R chimera (25 ng/mL) or IFN-β (5 ng/mL) added concurrently (*n* = 6 wells). **c**, Percent VCAM1−/C3+, VCAM1+/C3−, or VCAM1+/C3+ cells measured by flow cytometry in NTC sgRNA iAstrocytes compared to iAstrocytes transduced with sgRNAs knocking down genes involved in IL-6 or IFN signaling (*n* = 5 wells). **d**, IL-6 or IFN-β concentration in conditioned media (measured by multi-spot electrochemiluminescence) from NTC iAstrocytes vs. iAstrocytes with knockdown of *CEBPB*, *STAT3*, *IRF1*, or *STAT1/2* treated with vehicle control or IL-1α+TNF+C1q (*n* = 4 wells). **e**, Model of transcription factors and signaling pathways controlling inflammatory reactivity induced by IL-1α+TNF+C1q. In panels **b**, **c**, and **d**, *P* values were calculated using linear regression for MFI values or beta regression for percentages (see Methods), and only comparisons with statistically significant differences are marked. Where appropriate, *P* values were adjusted for multiple testing (*P*_adj_; Holm’s method) per family of tests (all comparisons made within a plot).

To better understand the behavior of VCAM1+/C3+ iAstrocytes, we treated iAstrocytes with all possible combinations of IL-1α, TNF, and C1q, with or without concurrent addition of exogenous IFN-β or IL-6 (in the form of an IL-6/IL-6R chimera; see Methods). We found that additional IL-6 boosted the induction of VCAM1–/C3+ iAstrocytes while inhibiting the induction of VCAM1+/C3– and VCAM1+/C3+ iAstrocytes, whereas additional IFN-β inhibited the induction of VCAM1–/C3+ iAstrocytes while boosting the induction of VCAM1+/C3– and VCAM1+/C3+ iAstrocytes (Fig. 6b). Given that IL-6 and IFN-β had similar effects on VCAM1+/C3– and VCAM1+/C3+ iAstrocytes, we mapped VCAM1+/C3+ iAstrocytes to IRAS2. This is also consistent with the fact that in our single-cell data, a subset of IRAS2 iAstrocytes had high transcript levels of both *VCAM1* and *C3* (Fig. 4b).

Upon examining the activity of IL-1α, TNF, and C1q alone or in combination, we found that IL-1α by itself induced VCAM1–/C3+ iAstrocytes, whereas TNF by itself induced VCAM1+/C3– and VCAM1+/C3+ iAstrocytes (Fig. 6b). Interestingly, IL-1α+TNF decreased the induction of VCAM1+/C3– astrocytes compared to TNF by itself (Fig. 6b). To validate these findings, we performed immunostaining against C3 and IFIT3, an alternative IRAS2 marker (Extended Data Fig. 7a-b), in iAstrocytes derived from three different hiPSC lines and observed similar results (Extended Data Fig. 7c). In terms of cytokine production (Extended Data Fig. 7d), IL-1α by itself, but not TNF by itself induced IL-6 and GM-CSF, whereas TNF by itself induced higher levels of CXCL10 compared to IL-1α by itself. Also, IL-1α+TNF decreased CXCL10 production compared to TNF by itself. These findings were similar between iAstrocytes derived from two different hiPSC lines (Extended Data Fig. 7d).

Taking stock of both VCAM1/C3 induction and cytokine production, the inhibitory effect of IL-1α on the induction of CXCL10 and VCAM1+/C3– iAstrocytes by TNF likely reflects the antagonistic effect of IL-1 on the activity of autocrine-paracrine interferon signaling^59^, which can be induced by both IL-1α and TNF (Extended Data Fig. 7d). As for the activity of C1q, C1q by itself did not induce any cytokine production or any inflammatory reactive astrocytes marked by VCAM1/C3; C1q also had little additional effect in combination with IL-1α, TNF, or IL-1α+TNF (Fig. 6b; Extended Data Fig. 7c-d). However, C1q may exert effects not detectable by the assays used above, e.g. at the transcriptional level.

With dual staining of VCAM1 and C3, we revisited the effect of knocking down receptors and transcription factors involved in IL-6 or interferon signaling (Fig. 6c) and found that the results were consistent with those obtained with single staining of VCAM1 or C3 (Fig. 5b). In addition, we also tested the effect of knocking down *CEBPB, CEBPD*, *NFKB2*, or *IRF1*, which were strong hits from the VCAM1 CRISPRi screen. Similarly to knockdown of *STAT3*, knockdown of *CEBPB, CEBPD*, or *NFKB2* reduced the induction of VCAM1–/C3+ iAstrocytes and increased the induction of VCAM1+/C3– or VCAM1+/C3+ iAstrocytes (Fig. 6c). Knockdown of *IRF1* reduced the induction of VCAM1+/C3– and VCAM1+/C3+ iAstrocytes and also marginally reduced the induction of VCAM1– /C3+ iAstrocytes (Fig. 6c), similarly to *STAT1/2* knockdown. We also tested the effect of *STAT3*, *CEBPB*, *NFKB2*, or *IRF1* knockdown in iAstrocytes derived from two additional hiPSC lines and obtained consistent results (Extended Data Fig. 8a-b).

In addition, we sought to better understand the regulation of autocrine-paracrine IL-6 and interferon production downstream of IL-1α+TNF+C1q treatment. We observed that production of IL-6 was decreased by *CEBPB*, *STAT3*, or *STAT1/2* knockdown, whereas production of IFN-β was decreased by on *STAT3*, *IRF1*, or *STAT1/2* knockdown (Fig. 6d).

Lastly, to complement our experiments employing CRIPSRi knockdown, we also tested the effect of small molecule known to modulate STAT3 or STAT1/2 activity. We found that napabucasin, which is known to inhibit STAT3-dependent transcription^71^, reduced STAT3 Y705 phosphorylation in a dose-dependent manner (Extended Data Fig. 9b) and at doses above 1.25 μM abrogated the induction of VCAM1–/C3+, VCAM1+/C3–, and VCAM1+/C3+ in IL-1α+TNF+C1q-treated iAstrocytes. The different effect of napabucasin vs. *STAT3* knockdown is not surprising given that the mechanism of action of napabucasin is not well understood and likely involves multiple cellular pathways^72^. For modulation of STAT1/2 activity, we found that RGFP966, a HDAC3 inhibitor known to reduce STAT1 activity^73^, boosted the induction of VCAM1–/C3+ iAstrocytes while inhibiting the induction of VCAM1+/C3– and VCAM1+/C3+ iAstrocytes (Extended Data Fig. 9d). The different effect of RGFP966 vs. *STAT1/2* knockdown could be attributed to effects of HDAC3 inhibition beyond reducing STAT1 activity.

Based on our findings so far and also existing literature, we created a model (Fig. 6e) of how IL-1α+TNF+C1q treatment induces IRAS1 and IRAS2 through a primary NF-κB-dependent transcriptional response (Fig. 4d-h) followed by autocrine-paracrine IL-6 and interferon signaling feedback loops involving *CEBPB/D*, *NFKB2*, *STAT3*, *STAT1/2*, and *IRF1* (Fig. 5, Fig. 6). Downstream of NF-κB, *CEBPB/D*, *NFKB2*, and *STAT3* drive the induction of IRAS1 and inhibit the induction of IRAS2 through IL-6 (also see refs. ^74, 75^), whereas *STAT1/2* and *IRF1* promote the induction of IRAS2 through interferons (also see ref. ^76^), which inhibit the induction of IRAS1 (Fig. 6b-d). IL-1α polarizes astrocytes towards IRAS1 by driving IL-6 production (Extended Data Fig. 7d) and antagonizing interferon signaling (also see ref. ^59^), whereas TNF polarizes astrocytes towards IRAS2 by driving interferon production (Extended Data Fig. 7d), which in turn antagonizes IL-1 signaling (also see ref. ^59^). The discrepancy between the inhibitory effect of IFN-β on IRAS1 induction (Fig. 6b) and the weak requirement of *STAT1/2* and *IRF1* for IRAS1 induction (Fig. 6c) could be explained by the promiscuity of *STAT1/2* in responding to and promoting the production of both IL-6 and IFN-β (Fig. 6d; also see refs. ^76, 77–79^). On the other hand, STAT3 activation has also been reported in response to interferons^80, 81^, and our data supports some role of STAT3 in the production of IFN-β (Fig. 6d); however, in our system STAT3 is much more strongly associated with IL-6 signaling given the consistent effect of IL-6 addition and *STAT3* knockdown.

We do not intend for the terms IRAS1 and IRAS2 to encompass reactive astrocyte signatures outside those generated within our experimental system. Thus, in light of the cellular pathways driving IRAS1 and IRAS2, we propose that IRAS1 be referred to as “IL-1/IL-6-responsive”, and IRAS2 as “TNF/IFN-responsive” reactivity.

### Signatures of IL-1/IL-6-responsive and TNF/IFN-responsive reactive astrocytes overlap with reactive astrocyte signatures found across different species and disease contexts

To see if signatures of IL-1/IL-6-responsive and TNF/IFN-responsive reactive astrocytes are found across different experimental paradigms of neuroinflammation, we integrated the NTC sgRNA-transduced iAstrocytes from our single-cell RNA-seq data with previously published astrocyte single-cell RNA-seq datasets using the anchor-based data integration functionality of Seurat^82^ (see Methods).

In Barbar *et al.*^29^, mixed cultures of neurons and glia derived from dissociation of human cerebral organoids were treated with vehicle control or IL-1α+TNF+C1q. After isolating astrocytes from the Barbar *et al.* dataset and integrating them with NTC iAstrocytes from our study (see Methods), we found that a subset of IL-1α+TNF+C1q-treated astrocytes from Barbar *et al.* co-clustered with IRAS1 (IL-1/IL-6 responsive) and IRAS2 (TNF/IFN-responsive) iAstrocytes (Fig. 7a-c). While Barbar *et al.* astrocytes co-clustering with IRAS1 iAstrocytes expressed IL-1/IL-6-responsive markers such as *IL6*, *CXCL2*, and *CXCL5*, those co-clustering with IRAS2 iAstrocytes expressed TNF/IFN-responsive markers such as *CXCL10*, *IFIT3*, and *ISG15* (Fig. 7d; Extended Data Fig. 10e; Supplementary Table 8). Furthermore, the DEGs between IRAS2 vs. IRAS1 iAstrocytes overlapped strongly with the DEGs between the corresponding co-clustering Barbar *et al.* astrocytes (Extended Data Fig. 10a).

**Fig. 7.**
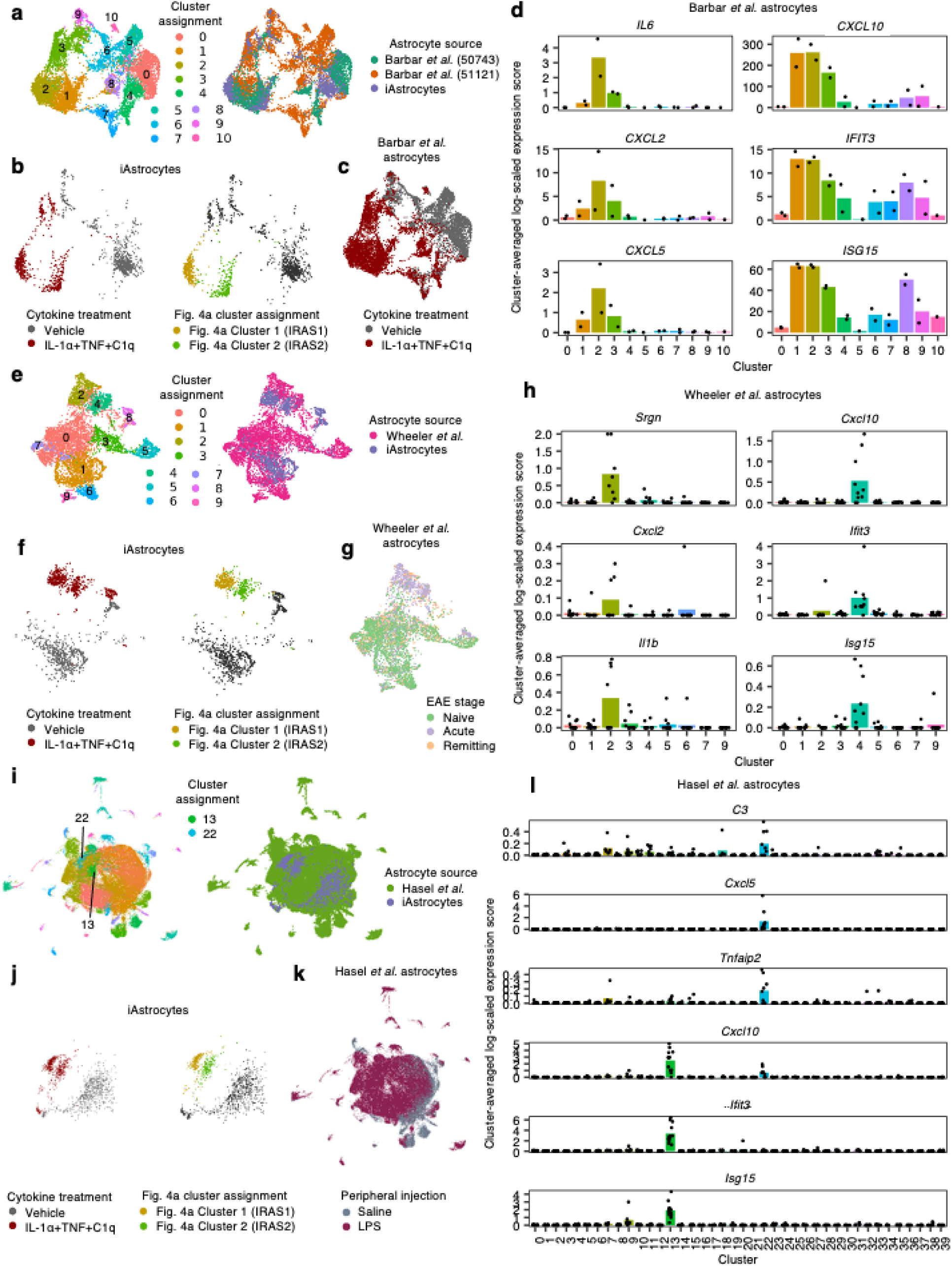
Integration of iAstrocyte single-cell data with published single-cell datasets shows overlap of inflammatory reactive signatures across species in diverse disease contexts. **a**, UMAP of integrated analysis (see Methods) of NTC sgRNA iAstrocytes with astrocytes from Barbar *et al.*, colored by cluster assignment or astrocyte source. **b**-**c**, The same UMAP embedding as in **a**, showing only iAstrocytes colored by cytokine treatment or Fig. 4a cluster assignment (**b**) or Barbar *et al.* astrocytes colored by cytokine treatment (**c**). **d**, Cluster-averaged expression levels of selected cluster markers in Barbar *et al.* astrocytes (*n* = 2 cell lines). **e**, UMAP of integrated analysis of NTC sgRNA iAstrocytes with astrocytes from Wheeler *et al.*, colored by cluster assignment or astrocyte source. **f**-**g**, The same UMAP embedding as in e, showing only iAstrocytes colored by cytokine treatment or Fig. 4a cluster assignment (**f**) or astrocytes from Wheeler *et al.* colored by EAE stage (**g**). **h**, Cluster-averaged expression levels of selected cluster markers in Wheeler *et al.* astrocytes (*n* = 8 mice). **i**, UMAP of integrated analysis of NTC sgRNA iAstrocytes with astrocytes from Hasel *et al.*, colored by cluster assignment or astrocyte source. **j**-**k**, The same UMAP embedding as in **i**, showing only iAstrocytes colored by cytokine treatment or Fig. 4a cluster assignment (**j**) or astrocytes from Hasel *et al.* colored by LPS treatment (**k**). **l**, Cluster-averaged expression levels of selected cluster markers in Hasel *et al.* astrocytes (*n* = 12 mice).

In Wheeler *et al.*^83^, astrocytes were isolated by FACS from mice induced with experimental autoimmune encephalomyelitis (EAE), a widely used model of multiple sclerosis (MS), at different stages of EAE progression (naïve, acute, and remitting). After removing low-quality astrocytes from the Wheeler *et al.* dataset and integrating the remaining astrocytes with NTC iAstrocytes from our study (see Methods), we found that the vast majority of Wheeler *et al.* astrocytes from the acute stage of EAE co-clustered with IRAS1 and IRAS2 iAstrocytes (Fig. 7e-g) and also correspondingly expressed IL-1/IL-6-responsive markers such as *Srgn*, *Cxcl2*, and *Il1b* or TNF/IFN-responsive markers such as *Cxcl10*, *Ifit3*, and *Isg15*, respectively (Fig. 7h; Extended Data Fig. 10f; Supplementary Table 8). Overlap analysis of the DEGs between IRAS2 vs. IRAS1 iAstrocytes and the DEGs between the corresponding co-clustering astrocytes from Wheeler *et al.* also showed good agreement (Extended Data Fig. 10b), although the degree of overlap (Jaccard index) was lower compared to the integration with Barbar *et al* likely due to the difference in species.

Lastly, in Hasel *et al.*^58^, astrocytes were isolated by FACS from mice injected peripherally with saline or LPS, which induces acute neuroinflammation similar to that caused by sepsis. After removing non-astrocyte cells from the Hasel *et al.* dataset, integration of the Hasel *et al.* astrocytes with NTC iAstrocytes from our study showed that a subset of Hasel *et al.* astrocytes from LPS-injected mice co-clustered with IRAS1and IRAS2 iAstrocytes (Fig. 7i-k) and also correspondingly expressed IL-1/IL-6 responsive markers such as *C3*, *Cxcl5*, and *Tnfaip2* or TNF/IFN-responsive markers such as *Cxcl10*, *Ifit3*, and *Isg15* (Fig. 7l; Extended Data Fig. 10g; Supplementary Table 8). Overlap analysis of the DEGs between IRAS2 vs. IRAS1 iAstrocytes and the DEGs between the corresponding co-clustering astrocytes from Hasel *et al.* further supported this correspondence (Extended Data Fig. 10c).

In addition to single-cell RNA-seq datasets, we also reanalyzed bulk RNA-seq data from Anderson *et al*. ^51^, where astrocyte-specific RNA was purified from wild-type or astrocyte-specific *Stat3* conditional knockout (cKO) mice subject to spinal cord injury (SCI). We found that Stat3-dependent genes (i.e. genes with lower expression in *Stat3* cKO SCI vs. WT SCI; e.g. *C3*) tended to be highly expressed by IRAS1 (IL-1/IL-6-responsive) iAstrocytes (Extended Data Fig. 10h), whereas Stat3-repressed genes (i.e. genes with higher expression in *Stat3* cKO SCI vs. WT SCI; e.g. *Iigp1*) were enriched for genes involved in the response to interferons (Extended Data Fig. 10i), overlapping with IRAS2 (TNF/IFN-responsive) markers (see Supplementary Table 9 for *Stat3* cKO-related DEGs from Anderson *et al.*). Thus, the results from Anderson *et al.* further corroborate the regulatory role of STAT3 in promoting IL-1/IL-6-responsive reactivity and inhibiting TNF/IFN-responsive reactivity that we have proposed.

### Markers of IL-1/IL-6-responsive and TNF/IFN-responsive reactive astrocytes are upregulated in human disease

Given that we found signatures of IL-1/IL-6 responsive and TNF/IFN-responsive reactive astrocytes in various experimental models of neuroinflammation, we sought to test whether they can be found in human disease. In Alzheimer’s disease (AD), neuroinflammation mediated by microglia and astrocytes contributes to disease progression^84^ potentially through IL-6^85^ and interferons^86^. We performed immunostaining of the IL-1/IL-6 responsive marker C3 and the TNF/IFN-responsive marker VCAM1 in post-mortem brain tissue derived from AD cases (61-92 years old, Braak stage IV-VI) vs. age-matched controls (62-102 years old, Braak stage 0-III; see Supplementary Table 10 for clinical metadata). We found a significant increase in the abundance of C3+ astrocytes in AD (Fig. 8a-b), consistent with previous studies^8^. Notably, C3+ astrocytes tended to be localized near C3+ plaques (Fig. 8a), which we deduced to be amyloid plaques given the fact that they are known to accumulate complement proteins such as C3 ^87^. Amyloid plaques are also known to accumulate SERPINA3 ^88^, which is a IL-1/IL-6 responsive marker and also a marker of disease-associated reactive astrocytes found in mouse models of AD^89^. As for VCAM1, we did not observe a statistically significant difference in the abundance of VCAM1+ astrocytes between AD cases vs. controls (Fig. 8c-d). Lastly, C3+ astrocytes tended to be VCAM1– (416 VCAM1– vs. 31 VCAM1+ astrocytes among C3+ astrocytes aggregated across all individuals; Fig. 8a), and VCAM1+ astrocytes tended to be C3– (135 C3– vs. 31 C3+ astrocytes among VCAM1+ astrocytes aggregated across all individuals; Fig. 8c); there were 31 VCAM1+/C3+ astrocytes among 12,885 total astrocytes.

**Fig. 8.**
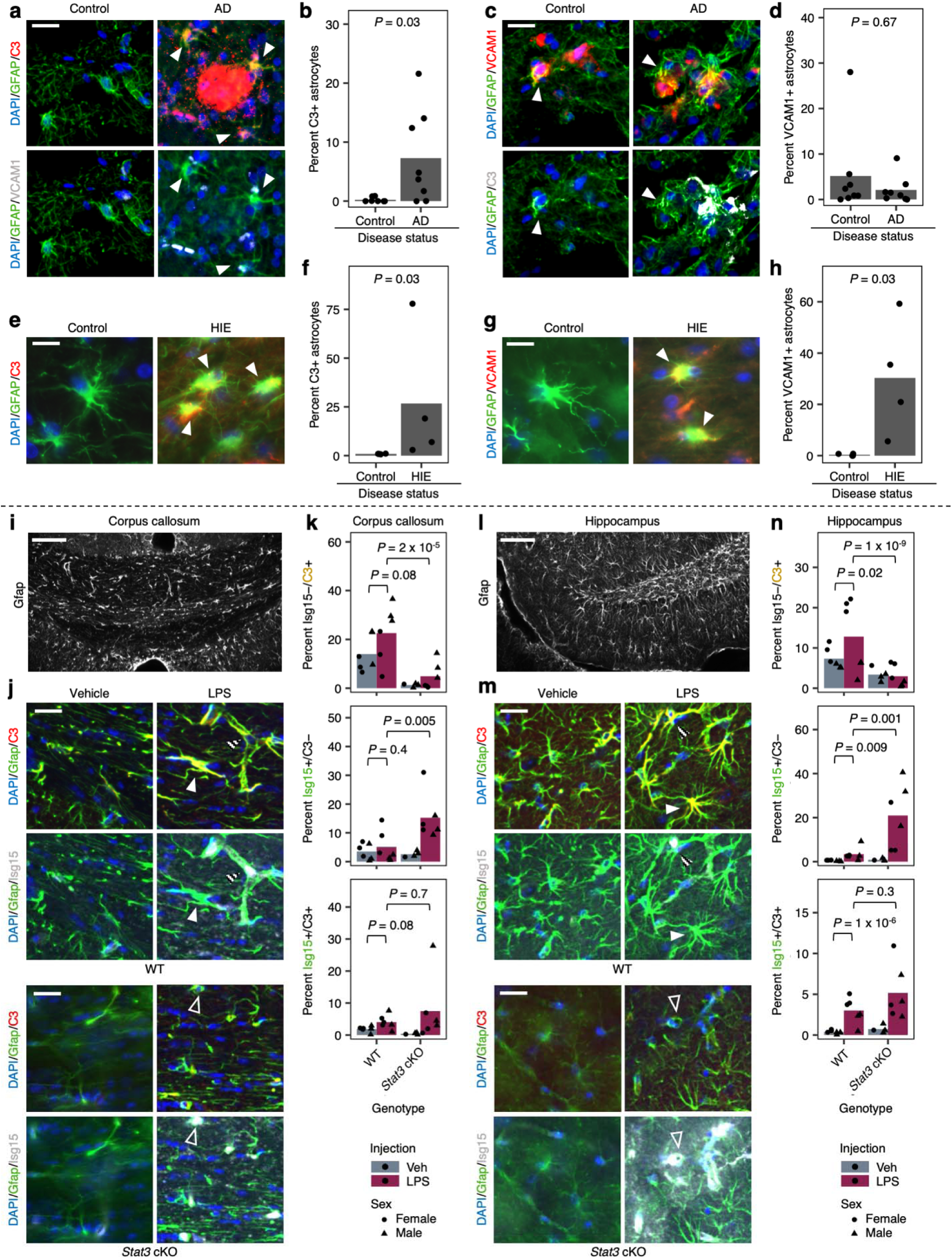
Markers of distinct inflammatory reactive signatures are upregulated in astrocytes in human Alzheimer’s disease (AD) and hypoxic-ischemic encephalopathy (HIE) and are regulated by Stat3 in a mouse model of neuroinflammation. **a-h,** Post-mortem human brain sections. **a,c**, Representative immunofluorescence images of C3+ (**a**) or VCAM1+ (**c**) astrocytes (marked by arrowheads) in an AD case compared to an age-matched control. Staining for VCAM1 and C3 was performed simultaneously (the same image is shown twice with different marker combinations). Scale bars correspond to 20 μm. **b,d**, Quantification of the percentage of C3+ (**b**) or VCAM1+ (**d**) astrocytes in AD cases (*n* = 4 individuals) vs. age-matched controls (*n* = 4 individuals). **e,g**, Representative immunofluorescence images of C3+ (**e**) or VCAM1+ (**g**) astrocytes (marked by arrowheads) in a HIE case compared to an age-matched control. Staining for VCAM1 or C3 was performed separately. Scale bars correspond to 10 μm. **f,h**, Quantification of the percentage of C3+ (**f**) or VCAM1+ (**h**) astrocytes in HIE cases (*n* = 4 individuals) vs. age-matched controls (*n* = 4 individuals). **i-n,** Brain sections from mice injected with vehicle or LPS. **i,l,** Representative immunofluorescence images of GFAP in the corpus callosum (**i**) or hippocampus (**l**). Scale bars correspond to 100 μm. **j,m,** Representative images of C3, Isg15, and Gfap staining in the corpus callosum (**j**) or hippocampus (**m**) in wild-type (WT) or Stat3 astrocyte-specific conditional knockout (Stat3-cKO) mice (n = 4-6 mice per group). Staining for C3 and Isg15 was performed simultaneously (the same image is shown twice with different marker combinations). Representative Isg15−/C3+ (solid arrowheads), Isg15+/C3− (empty arrowheads), and Isg15+/C3+ astrocytes (striped arrowheads) are labeled. **k,n,** Quantification of the percentage of Isg15−/C3+, Isg15+/C3−, and Isg15+/C3+ astrocytes in the corpus callosum (**k**) or hippocampus (**n**). *P* values were calculated using the two-sided Mann-Whitney U-test in **b**, **d**, **f**, and **h** and beta regression in **k** and **n**. For beta regression in **k** and **n**, sex was included as a covariate.

In addition to AD, we also examined brain tissue derived from age-matched controls (38-41 gestational weeks) vs. cases of neonatal hypoxic-ischemic encephalopathy (HIE; 37-41 gestational weeks), in which neuroinflammation involving IL-1, TNF, IL-6, and interferons^90–92^ contributes significantly to neuronal injury and negative neurologic sequela^93^. Compared to AD, HIE may be better modeled by our experimental paradigm (hiPSC-derived astrocytes treated with cytokines) given its acute course and the immature status of astrocytes in neonates^94, 95^. We observed increased abundance of both C3+ and VCAM1+ astrocytes in HIE (Fig. 8e-h). However, experimental limitations prevented us from staining for C3 and VCAM1 simultaneously.

### *Stat3* drives reactive astrocytes expressing IL-1/IL-6-responsive markers and inhibits those expressing TNF/IFN-responsive markers *in vivo*

Lastly, to validate *in vivo* the regulatory role of STAT3 in promoting IL-1/IL-6-responsive reactivity and inhibiting TNF/IFN-responsive reactivity, we performed immunostaining against the IL-1/6-responsive marker C3, the TNF/IFN-responsive marker Isg15, and the astrocyte marker Gfap in mice peripherally injected with vehicle control or LPS to induce acute neuroinflammation (see Methods). We focused on the corpus callosum and the hippocampus, where we observed easily segmentable Gfap immunoreactivity for astrocyte labeling (Fig. 8i,l) so we could verify astrocyte specificity. We found that in wild-type mice, LPS induced Isg15−/C3+ astrocytes (IL-1/IL-6-responsive) as well as Isg15+/C3− and Isg15+/C3+ astrocytes (TNF/IFN-responsive) in the hippocampus (Fig. 8m,n) but not the corpus callosum (Fig. 8j,k). In both brain regions, astrocyte-specific conditional knockout of *Stat3* in LPS-treated mice reduced the prevalence of Isg15−/C3+ astrocytes and increased the prevalence of Isg15+/C3− astrocytes (Fig. 8j,k,m,n) compared to wild-type LPS-treated mice, validating our *in vitro* finding of *Stat3* as a key regulator with opposing effects on IL-1/IL-6-responsive and TNF/IFN-responsive reactivity.

## DISCUSSION

Inflammatory reactive astrocytes have been implicated in numerous neurodegenerative and neuroinflammatory diseases such as Alzheimer’s disease, Parkinson’s disease, Huntington’s disease, amyotrophic lateral sclerosis, and multiple sclerosis^8, 11^. However, we do not yet have a full understanding of the cellular pathways controlling inflammatory reactivity in these contexts. Here, we employed pooled CRISPRi screening to systematically identify genes that control the response of hiPSC-derived astrocytes to the cytokines IL-1α+TNF+C1q, which induce a form of inflammatory reactivity associated with neurotoxicity under certain conditions and loss of homeostatic functions^8–10^.

The scalability and homogeneity (see Methods) of our hiPSC-derived astrocytes (“iAstrocytes”) was critical to our ability to perform pooled screens. However, it should be noted that iAstrocytes represent one of many available hiPSC-derived astrocyte models, and that there is a tradeoff between the scalability and maturation status of hiPSC-derived astrocytes. For our study, we found that iAstrocytes derived from multiple hiPSC lines sufficiently recapitulated published phenotypes associated with IL-1α+TNF+C1q-induced inflammatory reactivity, and we also ensured that the phenotypes we investigated were conserved across different hiPSC-derived astrocyte models and different hiPSC lines.

In our analysis of the results from CRISPRi screens and computational master regulator analysis (MRA), we gave more weight to the results from the CRISPRi screens, which demonstrate causality, than those from MRA, which are correlational and likely to contain more false positive results. However, given that the CRISPRi screens are more likely to be limited by false negative results (e.g. due to sgRNA dropout), further exploration of the hits from MRA is warranted in future studies. We believe that the gene regulatory network we generated from MRA will have broad utility for identifying potential master regulators of diverse astrocyte phenotypes for which transcriptomic data is available.

Following up on the top hits from our CRISPRi screens with single-cell transcriptomics, we found that IL-1α+TNF+C1q induced two distinct inflammatory reactive astrocyte signatures – provisionally named “IRAS1” and “IRAS2” – dependent on canonical NF-κB activation and driven by autocrine-paracrine IL-6 and interferon signaling. We observed that *CEBPB/D*, *NFKB2*, and *STAT3* promoted IRAS1 while inhibiting IRAS2 through IL-6, whereas *STAT1/2* and *IRF1* promoted both IRAS1 and IRAS2 partially through IFN-β; IL-1α polarized astrocytes towards IRAS1 by driving IL-6 production and antagonizing interferon signaling, while TNF polarized astrocytes towards IRAS2 by driving interferon production and antagonizing IL-1 signaling. Given the findings above, we proposed that IRAS1 be referred to as “IL-1/IL-6-responsive” and that IRAS2 be referred to as “TNF/IFN-responsive” reactivity.

Although our data supporting the above conclusions were largely *in vitro*, which is limited by reductionistic conditions compared to the complex *in vivo* environment, we validated the regulation of IL-1/IL-6-responsive and TNF/IFN-responsive reactivity by Stat3 *in vivo*. Furthermore, we found signatures of IL-1/IL-6-responsive or TNF/IFN-responsive astrocyte reactivity in other experimental contexts, both *in vitro* and in mouse models *in vivo,* as well as in human Alzheimer’s disease (AD) and hypoxic-ischemic encephalopathy (HIE).

It is likely that IL-1/IL-6-responsive and TNF/IFN-responsive reactive astrocytes have distinct functional outputs. Several studies already point towards potential functional outputs of IL-1/IL-6-responsive reactivity, which is dependent upon STAT3. For example, in Anderson *et al.* ^51^, astrocyte-specific deletion of *Stat3* was shown to prevent proper axon regeneration after spinal cord injury, suggesting that IL-1/IL-6-responsive reactivity may promote axon regeneration. In addition, in Kim *et al.*^96^, inhibition of STAT3 in hiPSC-derived astrocytes co-cultured with hiPSC-derived brain endothelial-like cells rescued the decrease in endothelial barrier integrity caused by treatment with TNF, suggesting that IL-1/IL-6-responsive reactivity may perturb blood-brain barrier and cerebrovascular function under inflammatory conditions^97^.

On the other hand, interferon-responsive reactivity may play an important role in EAE and MS. For example, Rothhammer *et al.*^98^ showed that inhibiting type I interferon signaling in astrocytes exacerbated the severity of EAE. Similarly, Hindinger *et al.*^99^ showed that inhibiting type II interferon in in astrocytes exacerbated the severity of clinical symptoms during peak disease. Furthermore, given that interferon-responsive reactive astrocytes upregulate VCAM1 and were shown to be adjacent to vasculature in Hasel *et al.*^58^, they may be important for controlling the trafficking of peripheral immune cells into the CNS parenchyma.

With regard to AD, literature on mouse models of AD supports a role for both IL-1/IL-6-responsive reactivity and TNF/IFN-responsive reactivity. Reichenbach *et al.* demonstrated that astrocyte-specific deletion of Stat3, which would be expected to reduce IL-1/IL-6-responsive reactivity, decreased the burden of amyloid plaques and dystrophic neurites, lowered inflammatory cytokine levels, and ameliorated cognitive deficits in the APP/PS1 mouse model of AD^100^. Roy *et al.* detected interferon-responsive astrocytes that were expanded in a amyloid pathology-dependent in the 5XFAD mouse model of AD^86^. Although we did not observe an increase in the percentage of VCAM1+ astrocytes in our examination of human AD brain tissue, this does not rule out the presence of interferon-responsive reactive astrocytes in AD, since VCAM1 may not be an optimal marker of interferon-responsive reactive astrocytes in all circumstances. Interestingly, Sadick *et al.* detected a strong interferon-responsive signature in astrocytes profiled by single-nucleus RNA-seq from a donor with vascular dementia^101^.

We believe that our iAstrocyte platform will assist in future investigations using patient-derived hiPSCs to dissect the effect of disease-associated mutations on inflammatory astrocyte reactivity. In addition, we believe our work here will serve as a valuable resource for future work to further characterize the functional outputs of IL-1/IL-6-responsive vs. TNF/IFN-responsive astrocyte reactivity in animal models of neuroinflammation and neurodegeneration. More generally, the approach pioneered here could be applied to uncover regulators of different reactive astrocytes signatures induced by other perturbations, which will pave the way for characterizing their functions and targeting them therapeutically.

## ACKNOWLEDGEMENTS

We thank Brandon Desousa, Vukasin Jovanovic, Zuzana Krejciova, and Nawei Sun for contributions to preliminary studies and discussions. We thank Professors Anna Molofsky, Aimee Kao, and Michael Oldham for serving on KL’s thesis committee. We thank members of the Kampmann lab (Greg Mohl, Sydney Sattler, Olivia Teter) for discussions and feedback on the manuscript. We thank Brian Woo for cloning the transcription factors sgRNA library, and Biswa Ramani for help with obtaining primary mouse astrocytes. We thank the Conklin lab for the gift of the WTC11 hiPSC line. This research was supported by NIH grant F30 AG066418 to KL, CIRM grant EDUC4-12812 & NIH grant T32 NS115706 to IVLR, Chan Zuckerberg Initiative Ben Barres Early Career Acceleration Awards to ESL and M. Kampmann, NIH/NINDs grants (R01NS097551, P01NS083513, R21NS119954) to SF, and NIH grants P30 EY02162-39 and R03AG063157 to EMU. SF is a Harry Weaver Neuroscience Scholar of the National Multiple Sclerosis Society. The TCW-1E44 iPSC line was generated with the support of NIH NIA K01AG062683 (J. TCW) and the Druckenmiller Fellowship from the New York Stem Cell Foundation (J. TCW).

## AUTHOR CONTRIBUTIONS

KL and M. Kampmann conceptualized and led the overall project, and wrote the manuscript with input from all co-authors. KL performed the majority of experiments with support from BR and performed all data analysis. IVLR performed immunostaining of mouse tissue provided by YA and SW with guidance from MVS. HK performed immunostaining of AD tissues provided by MSS and co-culture experiments with guidance from ESL, and WX performed immunostaining of HIE tissue with guidance from SF. WRF also performed immunostaining of AD tissues provided by MSS. EL generated the CRISPRi TCW-1E44 hiPSC line. J. TCW supplied the TCW-1E44 hiPSC line with guidance from AG. M. Koontz generated hiPSC-derived astrocytes using the methods of Krencik *et al.* and Li *et al.* with guidance from EMU, who also provided the 162D iPSC line. M. Krawcyzk and YZ supplied unpublished human astrocyte RNA-seq data for master regulator analysis.

## DATA AVAILABILITY STATEMENT

Bulk RNA-seq data of hiPSC-derived astrocytes generated in this study shown in Extended Data Fig. 3 are available on the Gene Expression Omnibus (GEO) under accession code GSE182307. The raw single-cell RNA-seq data and UMI count matrices from the CROP-seq experiment are available on GEO under accession code GSE182308. Processed data from the CRISPRi screens and CROP-seq experiment can also be interactively explored on CRISPRbrain (https://www.crisprbrain.org/).

## CODE AVAILABILITY STATEMENT

The full analysis pipeline (including code and processed data objects) used for master regulator analysis, analysis of CROP-seq data, and integration with previously published single-cell RNA-seq datasets is available at https://kampmannlab.ucsf.edu/inflammatory-reactive-astrocyte-analysis.

## METHODS

### Human iPSC (hiPSC) culture

hiPSCs lines (male WTC11^102^, female TCW-1E44^103^ or female 162D^104^ background) were obtained from the Conklin laboratory (UCSF), Goate laboratory (Icahn School of Medicine at Mt. Sinai), and the Ullian laboratory (UCSF), respectively. For the TCW-1E44 line, the consent for reprogramming human somatic cells to hiPSC was carried out under hSCRO protocol 19-04 (J. TCW).

hiPSCs were cultured in Essential 8 (E8) Medium (ThermoFisher Scientific cat. no. A1517001) on BioLite Cell Culture Treated Dishes (ThermoFisher Scientific) coated with Growth Factor Reduced, Phenol Red-Free, LDEV-Free Matrigel Basement Membrane Matrix (Corning cat. no. 356231) diluted 1:100 in DMEM/F12 (ThermoFisher Scientific cat. no. 11330032). Essential 8 Medium was replaced daily. When hiPSC colonies demonstrated mature morphology, the hiPSCs were either clump passaged with EDTA for routine maintenance or dissociated to a near single-cell suspension with Accutase Cell Dissociation Reagent (ThermoFisher Scientific cat. no. A11105-01) for applications requiring cell counting. For clump passaging with EDTA, hiPSCs were washed with Dulbecco’s phosphate buffered saline (DPBS; Milipore Sigma cat. no. D8537) and then incubated with Versene (ThermoFisher Scientific cat. no. 15040066) for 5-7 min at room temperature; the Versene solution was then aspirated and replaced with E8 + 10 nM Y-27632 dihydrochloride ROCK inhibitor (Tocris cat. no. 125410); hiPSC colonies were then gently detached mechanically using a cell scraper, resuspended gently, and passaged at 1:10-1:30 dilution in E8 + Y-27632, with Y-27632 removed the next day. For near single-cell dissociation, hiPSCs were washed with DPBS, incubated with Accutase for 5-10 min at 37 °C, and then gently triturated with a P1000 pipette tip; the cell suspension was then diluted with PBS, collected into conical tubes, and spun down at 300 g for 3 min; hiPSCs were then resuspended in E8 + Y-27632, counted, and plated onto Matrigel-coated plates at the desired density in E8 + Y-27632; Y-27632 would be maintained until the hiPSC colonies reached the appropriate size (>∼40 cells). Studies with hiPSCs at UCSF were approved by The Human Gamete, Embryo and Stem Cell Research (GESCR) Committee. Informed consent was obtained from the human subjects when the hiPSC lines originally derived.

### Cloning of *NFIA* and *SOX9* cDNA into dox-inducible cassette

To obtain *NFIA* cDNA, we designed PCR primers to amplify cDNA corresponding to transcript ENST00000403491(*NFIA* isoform 1) from astrocyte cDNA (Forward primer complementary sequence: ATGTATTCTCCGCTCTGTCTCAC; reverse primer complementary sequence: TCCCAGGTACCAGGACTGTG). We chose to amplify cDNA corresponding to *NFIA* isoform 1 because the cDNA clone (BC022264) used in Li *et al*.^19^ corresponds to transcript ENST00000371187 (*NFIA* isoform 2), which we found was not expressed highly in human astrocytes. To obtain *SOX9* cDNA, we ordered cDNA clone OHu19789 (which corresponds to MGC clone BC056420) from GeneScript and then amplified *SOX9* cDNA from the plasmid (Forward primer complementary sequence: ATGAATCTCCTGGACCCCTTCA; reverse primer complementary sequence: TCAAGGTCGAGTGAGCTGTGT). We then inserted *NFIA* and *SOX9* cDNA joined by a T2A sequence into an AAVS1 safe-harbor plasmid containing a dox-inducible cassette (Addgene plasmid no. 105840, gift from Michael Ward; digested with AflIIand ClaI) using Gibson assembly (New England Biolabs; cat. no. E2611L), resulting in pKL100.

### Generation of hiPSC lines with stable integration of NFIA-SOX9 cassette and CRISPRi cassette

WTC11, TCW-1E44, and 162D hiPSCs were transfected with pC13N-dCas9-BFP-KRAB^105^ (Addgene plasmid no. 127968) to stably integrate constitutive CRISPRi machinery into the CLYBL locus using TALEN-based editing as previously described^105^. CRISPRi WTC11, TCW-1E44, and 162D hiPSCs were then transfected with pKL100 to stably integrate the dox-inducible NFIA-SOX9 cassette into the AAVS1 locus using the same TALEN-based editing approach. Briefly, CRISPRi WTC11, TCW-1E44, and 162D hiPSCs were dissociated with Accutase to a single-cell suspension and plated at 500,000 cells per well in a Matrigel-coated 6-well plate in E8 + 10 µM Y-27632. The next day, a media change with E8 + Y-27632 was performed and then the hiPSCs were transfected with 2.5 ug of pKL100, 1.25 ug of left and right AAVS1 TALEN plasmids (Addgene plasmid no. 59025 and 59026, gift from Danwei Huangfu), and 0.5 ug of Bcl-XL plasmid (pEF1-BCL-XL-wpre-polyA P1102, gift from Xiaobing Zhang, described in ref ^106^) using Lipofectamine Stem Transfection Reagent (ThermoFisher Scientific cat. no. STEM00015) following the manufacture’s protocol. After hiPSCs reached confluence, they were dissociated with Accutase and passaged to a 10 cm dish in E8 + 10 uM Y-27632 + 0.1 ug/mL puromycin (ThermoFisher Scientific cat. no. A1113803) to select for clones with stable integration. Y-27632 was maintained until stable colonies formed, and puromycin was maintained for 5-7 days. The hiPSCs were then sorted for mCherry+ cells, which were then plated at 500,000 cells per well in a Matrigel-coated 6-well plate for transfection with 1.2 ug of Cre recombinase mRNA (TriLink Biotechnologies cat. no. L-7211) using Lipofectamine Stem to remove the puromycin resistance gene and mCherry. The Cre-transfected hiPSCs were then expanded and sorted for mCherry-cells, which were then plated for colony picking to generate monoclonal hiPSC lines. hiPSC clones were tested for integration of the NFIA-SOX9 cassette and removal of the puromycin resistance and mCherry by genomic PCR with the following pairs of primers:

AAVS1 FWD: CTGCCGTCTCTCTCCTGAGT

bGHpolyA REV: GCTGGCAACTAGAAGGCACAG

AAVS1 FWD: CTGCCGTCTCTCTCCTGAGT

Puro REV: GTGGGCTTGTACTCGGTCAT

TRE3G FWD: GTGTTGTGGAATTGCTCCAG

AAVS1 REV: AAGAGTGAGTTTGCCAAGCAGT

T2A-SOX9 FWD: AATCCCGGCCCTATGAATCTCCTG

T2A-SOX9 REV: TGGGCGATCGATTCAAGGTCGAGT

### Neural induction of hiPSCs

Embryoid body (EB)-based neural induction of WTC11, TCW-1E44, and 162D hiPSCs stably integrated with CRISPRi machinery and dox-inducible NFIA-SOX9 was performed as previously described^107^ with some modifications. Briefly, on day 0, hiPSCs were dissociated to a near single-cell suspension with Accutase, resuspended in neural induction media (NIM; see recipe below) + 10 nM Y-27632, and then transferred to an Aggrewell 800 plate (StemCell Technologies cat. no. 34815) pre-coated with Anti-adherence rinsing solution (StemCell Technologies cat. no. 07010) at 3 million cells per well for EB formation. The next day (day 1), a half media change with NIM was performed and LDN193189 (LDN; Tocris cat. no. 6053) and SB431542 (SB; Tocris cat. no. 1614) were added to final concentrations of 0.1 μM and 10 μM, respectively. A half media change of NIM + LDN + SB was performed every day until day 7, when EBs were transferred to a Matrigel-coated 6-well plate after performing a half media change (one well from the Aggrewell plate would be transferred to one well in the 6-well plate). The next day (day 8), a full media change with NIM + LDN + SB was performed, and then every other day afterwards until day 14, during which time neural rosettes would appear in the attached EBs. On day 14, neural rosettes were detached non-enzymatically with the following method: the attached EBs were incubated Neural Rosette Selection Reagent (NRSR; StemCell Technologies cat. no. 05832) for 1 hour at 37 °C to weaken the attachment of the neural rosettes, the NRSR was aspirated and replaced with DMEM/F12, and the rosettes were detached with targeted jetting of DMEM/F12 using a wide-orifice P1000 pipette tip and then collected into a conical tube; targeted jetting with DMEM/F12 and collection of released rosettes was repeated until the majority of rosettes had detached. Rosettes were then spun down at 100 g for 3 min, resuspended in NPC media (see recipe below), and transferred to a Matrigel-coated 6-well plate (rosettes collected from 1-2 wells would be replated into 1 well depending on the yield). From day 15 to 21, a full media change with NPC media was performed every other day, during which time neural progenitor cells (NPCs) would spread out from the attached rosettes and cover the well completely. Once NPCs reached confluency, they were dissociated with Accutase and replated at high density (at least 1 million cells per well of a 6-well plate) in Matrigel-coated plates for expansion.

Neural induction media (NIM) formulation:

DMEM/F12 (+HEPES, +Glutamine) basal media (ThermoFisher Scientific cat. no. 11330032)

2% (v/v) B27 supplement minus Vit. A (ThermoFisher Scientific cat. no. 12587010)

1% (v/v) N2 supplement (ThermoFisher Scientific cat. no. 17502048)

Neural progenitor cell (NPC) media formulation:

DMEM/F12 (+HEPES, +Glutamine) basal media (ThermoFisher Scientific cat. no. 11330032)

2% (v/v) B27 supplement minus Vit. A (ThermoFisher Scientific cat. no. 12587010)

1% (v/v) N2 supplement (ThermoFisher Scientific cat. no. 17502048)

20 ng/mL bFGF (PeproTech cat. no. 100-18B)

### Purification of NPCs

To remove contaminating neural crest cells from the NPC cultures, we used fluorescence activated cell sorting (FACS) to select for CD133+/CD271-cells following the protocol described in Cheng *et al.*^107^ with modifications. Briefly, NPCs were dissociated to a single-cell suspension with Accutase, resuspended in FACS buffer (see recipe below), and then incubated with PE-conjugated CD133 antibody (1:50 dilution; Miltenyi Biotec cat. no. 130-113-108) and PerCP-Cy5.5-conjugated CD271 antibody (1:50 dilution; BD Biosciences cat. no. 560834), including single antibody-stained and unstained controls. After antibody incubation, the cell suspension was diluted 10x with FACS buffer, spun down, and resuspended in FACS buffer for sorting. CD133+/CD271-cells were sorted using a BD FACSAria Fusion cell sorter at 5,000-10,000 events per second, and then plated at 100,000 cells per cm^2^ onto Matrigel-coated plates. Media was then changed every other day until the NPCs reached confluency, at which point the NPCs were passaged for expansion. After 2-3 additional expansion passages after sorting, NPCs were characterized by immunostaining and qPCR for NPC markers such as PAX6 and Nestin, and then cryopreserved in NPC media + 10% DMSO.

FACS buffer formulation:

DPBS (Milipore Sigma cat. no. D8537)

1% (w/v) BSA (Milipore Sigma cat. no. A9647)

2 mM EDTA (Milipore Sigma cat. no. 324506)

### Generation of iAstrocytes from NPCs

For iAstrocyte differentiation, CD133+/CD271-sorted NPCs generated from WTC11, TCW-1E44, and 162D iPCS with stably integrated CRISPRi machinery and dox-inducible NFIA-SOX9 were dissociated to a single-cell suspension with Accutase and then replated at 7,500-15,000 cells per cm^2^ in NPC media onto a 10-cm or 15-cm dish coated with Matrigel diluted at 1:200 in DMEM/F12. The next day, media was changed to ScienCell Astrocyte Media (ScienCell Research Laboratories cat. no. 1801) + 2 μg/mL doxycycline (Millipore Sigma cat. no. D9891) to initiate iAstrocyte differentiation. A full media was then changed every other day, with doxycycline maintained at 2 μg/mL throughout the differentiation process. When the differentiating NPCs reached confluency within 3-4 days, the culture was dissociated with Accutase and split 1:4-1:10 onto new Matrigel coated dishes for expansion, with some cells saved for cryopreservation. After confluency was reached within 5-6 days, the cultures were dissociated and split 1:4-1:8, saving some cells for cryopreservation. Expansion of the cultures at 1:4-1:8 split with cryopreservation of cells after each split was continued until day 20 of differentiation, yielding iAstrocytes. To maintain the post-split cell density between 10,000-20,000 cells per cm^2^, the split ratio was adjusted based on the proliferative tendency of the differentiating astrocyte precursors, which varied those generated from WTC11, TCW-1E44, and 162D hiPSCs. In terms of yield, starting from ∼400,000 NPCs plated into a 10 cm dish, in theory ∼100 million iAstrocytes could be obtained by around day 20 assuming that the cells are split 1:4 for a total of four times. However, the yield may vary depending on the proliferative tendency of the astrocyte precursors.

For cryopreservation, iAstrocytes were resuspended in ScienCell Astrocyte media + 10% DMSO, transferred into Corning CoolCell alcohol-free freezing containers (Corning cat. no. 432001) in cryogenic vials, and then transferred to liquid nitrogen. For experiments, cryopreserved iAstrocytes were thawed by warming cryovials in a 37 °C water bath until no ice was left, transferring the contents to a 15 mL conical tube, adding 4 volumes of DPBS, equilibrating at RT for 3 min, spinning down cells at 300 g for 3 min, and then resuspending the pellet in ScienCell Astrocyte Media.

Experiments were performed using iAstrocytes derived from WTC11 hiPSCs unless otherwise indicated.

### Generation of hiPSC-derived astrocytes using alternative protocols

In parallel to the generation of iAstrocytes, WTC11 hiPSC-derived astrocytes were also generated from CD133+/CD271-sorted NPCs according to the protocol published in TCW *et al.*^18^, which mirrored the process described above for iAstrocyte generation with the exception of adding doxycycline. In addition to the protocol published in TCW *et al.*, hiPSC-derived astrocytes were also generated from WTC11 iPSCs with stably integrated CRISPRi machinery and dox-inducible NFIA-SOX9 according to Li *et al.*^19^, or from WTC11 iPSCs with stably integrated CRISPRi machinery according to Krencik *et al.*^15^.

### Primary mouse astrocyte culture

P0 C57BL/6J pups were decapitated and the brains were removed and placed into ice cold HBSS. Cortices were removed from the rest of the brain in HBSS and dissociated using the Pierce Primary Neuron Isolation Kit (ThermoFisher Scientific cat. no. 88280). Instead of using the Neuronal Culture Medium from the kit, cells from the dissociated cortices were plated onto PDL-coated 96-well plates in primary astrocyte medium (see recipe below) at the recommended density. Media was changed every other day for 6 days, at which point the culture consisted mainly of astrocytes.

Primary astrocyte medium:

DMEM (ThermoFisher Scientific cat. no. 11965084)

10% FBS (VWR cat. no. 89510-186; lot no. 184B19)

1% Penicillin-streptomycin (ThermoFisher Scientific cat no. 15140122)

### Induction of inflammatory reactivity in hiPSC-derived astrocytes or primary mouse astrocytes

iAstrocytes or hiPSC-derived astrocytes generated according to TCW *et al.*^18^ were plated onto Matrigel-coated (1:200 diluted Matrigel) 96-well plates, 24-well plates or 6-well plates at 20,000 cells per cm^2^ in ScienCell Astrocyte Media without addition of doxycycline; hiPSC-derived astrocytes generated according to Li *et al.*^19^ or Krencik *et al.*^15^ were plated onto Matrigel-coated (1:200 diluted Matrigel) 96-well plates at ∼62,500 cells per cm^2^ in Astrocyte Maturation Media (AMM; see recipe below) without addition of doxycycline. A full media change was performed the next day, and then every other day afterwards. Five days after plating, hiPSC-derived astrocytes were treated with vehicle control or IL-1α (3 ng/mL; Peprotech cat. no. AF-200-01A), TNF (30 ng/mL; Peprotech cat. no. AF-300-01A), and C1q (400 ng/mL; Complement Technology cat. no. A099) with a full media change in the appropriate media to induce inflammatory reactivity according to Liddelow *et al.*^8^. For experiments involving all possible combinations of IL-1α, TNF, and C1q, the same concentrations as above were used. For experiments involving treatment with additional IL-6 or IFN-β, a IL-6/IL-6R chimeric protein (25 ng/mL; R&D systems cat. no. 8954-SR-025) or IFN-β (5 ng/mL; Peprotech cat. no. AF-300-02B) was added concurrently with all possible combinations of IL-1α, TNF, and C1q, which were added at the same concentration as above. We found that using a IL-6/IL-6R chimeric protein was necessary given our observation that addition of IL-6 itself had no additional effect on iAstrocytes, which is likely due to the fact that IL-6R is not expressed at high levels constitutively in most cell types^108^.

For primary mouse astrocytes, IL-1α+TNF+C1q treatment (same concentration as above) occurred after 6 days of culture. All assays were performed at 24 hours after cytokine treatment (see Fig. 1g) unless otherwise stated.

Astrocyte Maturation Media (AMM) formulation:

DMEM/F12 (+HEPES, +Glutamine) basal media (ThermoFisher Scientific cat. no. 11330032)

1% (v/v) B27 supplement minus Vit. A (ThermoFisher Scientific cat. no. 12587010)

0.5% (v/v) N2 supplement (ThermoFisher Scientific cat. no. 17502048)

1% (v/v) Antibiotic-Antimycotic (ThermoFisher Scientific cat. no. 15240062)

10 ng/mL CNTF (R&D Systems cat. no. 257-NT)

10 ng/mL BMP-4 (R&D Systems cat. no. 314-BP)

### CRISPRi-mediated gene knockdown in hiPSC-derived astrocytes using lentiviral sgRNA delivery

For experiments involving CRISPRi-mediated gene knockdown, iAstrocytes were transduced with lentivirus containing single-guide RNAs (sgRNAs) at the time of plating (see Fig. 2b). CRISPRi sgRNAs were cloned into pMK1334 ^105^ (Addgene cat. no. 127965) digested with BstXI and BlpI as previously described in Gilbert *et al.*^36^. Lentivirus containing sgRNAs was produced by transfecting HEK293T cells with pMK1334 and 3^rd^ generation lentiviral packaging plasmids with TransIT-Lenti Transfection Reagent (Mirus cat. no. MIR6606) according to the manufacturer’s instructions. The lentivirus was then precipitated using Lentivirus Precipitation Solution (ALSTEM cat. no. VC150) according to the manufacturer’s instructions, resuspended in DPBS at 1/10 of the original volume, and then aliquoted and stored at −80 °C. The functional titer of the lentivirus was then tested on iAstrocytes by serial dilution followed by measurement of BFP+ cells 48 hours after transduction. For all experiments involving CRISPRi-mediated gene knockdown in iAstrocytes, sufficient sgRNA lentivirus was added to transduced >70% of iAstrocytes.

### Immunostaining of hiPSC-derived astrocytes

iAstrocytes and hiPSC-derived astrocytes generated according to TCW *et al.*^18^ from WTC11, TCW-1E44, and 162D hiPSCs were plated at 20,000 cells per cm^2^ onto a Greiner μClear 96-well plate (Greiner Bio-One cat. no. 655087) coated with 1:200 diluted Matrigel, and then treated with vehicle control or IL-1α+TNF+C1q as described above. 24 hours after cytokine treatment, the astrocytes were washed with DPBS and then fixed with 4% paraformaldehyde (diluted from a 16% solution; Electron Microscopy Sciences cat. no. 15710) for 15 min at room temperature (RT). After washing three times with DPBS, blocking and permeabilization was performed with DPBS + 3% BSA + 0.1% Triton X-100 (Millipore Sigma cat. no. X100) for 30 min at RT. Primary antibodies against GFAP (1:500, rabbit polyclonal; ThermoFisher Scientific cat. no. PA1-10019), S100β (1:500, mouse monoclonal; Millipore Sigma cat. no. S2532), GLAST (1:500, mouse monoclonal; Miltenyi Biotec cat. no. 130-095-822), NFIA (1:200 rabbit polyclonal; Atlas Antibodies cat. no. HPA008884), Cx43 (1:250, rabbit polyclonal; ThermoFisher Scientific cat. no. 71-0700), glutamine synthetase (1:500, mouse monoclonal; ThermoFisher Scientific cat. no. MA5-27749), or vimentin (1:200, rabbit monoclonal; Cell Signaling Technologies cat. no. 5741) were then added in blocking buffer and incubated overnight at 4 °C. Afterwards, the samples were washed with DPBS + 0.1% Triton X-100 three times, incubated with pre-adsorbed secondary antibodies (1:500 Goat anti-mouse IgG AF647, 1:500 Goat anti-rabbit IgG AF555; Abcam cat. no. ab150119 and ab150086) for 1 hour at RT, washed two times with DPBS + 0.1% Triton X-100, incubated with 1 μg/mL Hoechst (ThermoFisher Scientific cat. no. H3570) and 1:10 ActinGreen 488 (ThermoFisher Scientific cat. no. R37110) for 20 min at RT, and then washed two additional times before imaging.

For immunostaining of C3 and IFIT3, iAstrocytes derived from WTC11, TCW-1E44, and 162D hiPSCs washed, fixed, and blocked as above and then incubated with primary antibodies against C3 (1:250, mouse monoclonal; BioLegend cat. no. 846302), IFIT3 (1:500, rabbit polyclonal; ThermoFisher Scientific cat. no. PA5-22230), or Na^+^/K^+^ ATPase (1:500, chicken polyclonal; VWR cat. no. 89162-626) overnight at 4 °C in blocking buffer. Afterwards, washing and secondary antibody incubation was performed as above (1:500 Goat anti-mouse IgG AF647, 1:500 Goat anti-rabbit IgG AF555, 1:500 Goat anti-chicken IgY AF647; Abcam cat. no. ab150119, ab150086, and ab150175) with 1 μg/mL Hoechst as a counterstain.

Samples were imaged with an IN Cell Analyzer 6000, using a 20X 0.45 NA objective, 2×2 binning, 100-400 ms exposure, an aperture width of ∼1 Airy unit, and 16 fields per well.

### Generation of hiPSC-derived brain endothelial-like cells and measurement of barrier integrity

hiPSCs were differentiated to brain endothelial-like cells as previously described^109^. Briefly, hiPSCs were dissociated with Accutase and seeded on Matrigel-coated plates in E8 medium containing 10 μM Y27632 at a density of 15,000 cells/cm^2^. Differentiation was initiated 24 hours after seeding by changing to E6 medium, with daily medium changes for 4 days. Next, cells were expanded with serum-free basal endothelial cell medium (EC medium) supplemented with 50x diluted B27 (Thermo Fisher Scientific), 1x GlutaMAX (Thermo Fisher Scientific), 10 μM retinoic acid (Sigma Aldrich), and 20 μg/ml FGF2 for 2 days without a media change. Following this treatment, cells were collected by a 20-minute incubation in Accutase and seeded onto Transwell filters (1.1 cm^2^ polyethylene terephthalate membranes with 0.4 μm pores; Fisher Scientific) coated with a mixture of 400 μg/ml collagen IV (Sigma Aldrich) and 100 μg/ml fibronectin (Sigma Aldrich). The following day, cells were switched to EC medium lacking FGF2 and RA. For co-culture with iAstrocytes, filters were transferred to 12-well plates containing iAstrocytes and the same medium was utilized. Starting at this time, transendothelial electrical resistance (TEER) was measured using STX2 chopstick electrodes and an EVOM2 voltameter (World precision Instruments) approximately every 24 hours. TEER readings on empty Transwell filters were subtracted from all measurements to reflect the resistance of only the cultured cells.

### Generation of hiPSC-derived neurons (iNeurons) and GCaMP iNeurons

hiPSC-derived neurons (iNeurons) were generated from WTC11 hiPSCs with stably integrated dox-inducible NGN2 (NGN2 iPSCs) according to Fernandopulle *et al.*^110^. To generate GCaMP iNeurons, NGN2 iPSCs were transduced with a lentivirus delivering GCaMP6m (gift from Dr. Michael Ward). To facilitate segmentation of neuron soma, an additional lentivirus transduction was performed to deliver pMK1334 containing a non-targeting sgRNA, which confers BFP expression localized to the nucleus. Clonal lines were then isolated by colony picking, differentiated to neurons according to Fernandopulle *et al.*^110^, and evaluated for homogeneity of GCaMP6m expression and the presence of spontaneous calcium oscillations. A clonal line satisfying the above criteria was selected for GCaMP imaging experiments.

### Measurement of calcium activity in GCaMP iNeurons

Briefly, GCaMP iNeurons at day 3 of differentiation (see Fernandopulle *et al.*^110^) were replated onto poly-D-lysine-coated 96-well plates (Corning cat. no. 354640) at 62,5000 cells per cm^2^ in neuron media + 2 μg/mL doxycycline. For co-culture experiments, iAstrocytes were added on day 3 at 10,000 cells per cm^2^ in an equivalent volume of ScienCell Astrocyte Media + 2 μg/mL doxycycline. For mono-culture experiments, an equivalent volume of neuron media + 2 μg/mL doxycycline was added. On day 6, half of the media was replaced with fresh neuron media + 2 μg/mL doxycycline; on day 10, half of the media was replaced with fresh neuron media without doxycycline. On day 17, the cultures were treated with vehicle control or IL-1α+TNF+C1q (using the same final concentration as described above for astrocyte experiments) by performing a half media change with fresh neuron media. Calcium activity in GCaMP iNeurons was recorded on day 18 and day 20 with an IN Cell Analyzer 6000, using a 20X 0.45 NA objective, 2×2 binning, environmental control set to 37 °C and 5% CO_2_, an aperture width of ∼1 Airy unit, and 1 frame (800 ms exposure) collected per second for 40 seconds per field (1 field per well).

### Isolation of synaptosomes and labeling with pHrodo

Synaptosomes were isolated from fresh Innovative Grade US Origin Rat Sprague Dawley Brains (Innovative Research, Inc.; Cat. No. IGRTSDBR) with the Syn-PER™ Synaptic Protein Extraction Reagent (ThermoFisher Scientific cat. no. 87793) according to the manufacture’s protocol with minor changes. Briefly, 10 mL of Syn-PER Reagent supplemented with 1x protease inhibitor cOmplete Mini, EDTA free (Roche cat. no. 11836170001) and 1x phosphatase inhibitor PhosSTOP (Roche cat. no. 4906845001) were added per gram of brain tissue. Dounce homogenization was performed on ice and homogenate was transferred to a conical tube and centrifuged at 1200 × g for 10 minutes at 4°C. The pellet was discarded, the supernatant was transferred to a new tube, and the centrifugation step was repeated. The supernatant was then centrifuged at 15,000 × g for 20 minutes at 4°C. The supernatant was removed and the wet pellet was weighed. The synaptosome fractions were resuspended at a concentration of 50 mg/ml. 3 μM of pHrodo Red, succinimidyl ester (ThermoFisher Scientific cat. no. P36600) was added to the synaptosome fraction and incubated for 45 min at room temperature in the dark. After diluting the solution 1:10 in DPBS, the synaptosomes were spun down at 2500 × g for 5 min. The supernatant was removed and then the synaptosomes were washed two times with DPBS. The pHrodo-labelled synaptosomes were resuspended in DMEM/F12 + 5% DMSO at a stock concentration of 50 mg/ml, aliquoted, and then frozen in liquid nitrogen for later use.

### Measurement of synaptosome phagocytosis

For synaptosome phagocytosis experiments, pHrodo-labeled rat synaptosomes were used for iAstrocytes, and pHrodo-labeled iNeuron synaptosomes were used for Li *et al.*^19^ and Krencik *et al.*^15^ hiPSC-derived astrocytes. Briefly, astrocytes were incubated with pHrodo-labeled synaptosomes resuspended in the appropriate astrocyte media (ScienCell Astrocyte media for iAstrocytes, AMM for Li *et al.* and Krencik *et al.* astrocytes) at 1 mg/mL for 3 hours at 37 °C; for negative controls, some samples were pre-treated with 10 uM cytochalasin D (Millipore Sigma cat. no. C8273) for 15 min and also incubated with synaptosomes in the presence of 10 uM cytochalasin D to inhibit phagocytosis. After incubation with pHrodo-labeled synaptosomes, astrocytes were washed with DPBS, dissociated with Accutase, and pHrodo fluorescence was measured by flow cytometry. The gating strategy to determine the percent of phagocytic cells was based on the separation between the fluorescence histograms of samples treated or not treated with cytochalasin D.

### Measurement of iNeuron viability in the presence of astrocyte conditioned media

Conditioned media was collected from iAstrocytes treated with vehicle control or IL-1α+TNF+C1q for 24 hours, spun down at 300 g for 10 min to remove dead cells, and transferred to day 17 iNeurons after removing the original iNeuron media. Unconditioned ScienCell Astrocyte Media was used as a negative control. After 72 hours, iNeuron viability was assessed by adding 10 ug/mL Hoechst and 1 μM TO-PRO-3 (ThermoFisher Scientific cat. no. T3605) in DPBS, incubating for 10 min at 37 °C, and then imaging on an IN Cell Analyzer 6000, using a 10X 0.45 NA objective, 2×2 binning, environmental control set to 37 °C and 5% CO_2_, an aperture width of ∼1 Airy unit, 200 ms exposure, and 4-9 fields per well. The percent of dead neurons (stained by TO-PRO-3) was calculated after image processing and segmentation with CellProfiler (see Data Analysis section).

### Bulk RNA-seq library prep

hiPSC-derived astrocytes (iAstrocytes, TCW *et al.*, Li *et al.*, and Krencik *et al.* astrocytes), NPCs, and iNeurons were cultured in their respective media and treated with vehicle control or IL-1α+TNF+C1q for 24 hours. RNA extraction was then performed with Zymo Quick-RNA Microprep kit (Zymo Research cat. no. R1051). 50-100 ng of RNA was then used to construct bulk RNA-seq libraries using the QuantSeq 3’ mRNA-Seq Library Prep Kit FWD for Illumina (Lexogen cat. no. 015.96) following the manufacturer’s instructions. The concentration of QuantSeq libraries mRNA-seq library was quantified using the Qubit dsDNA HS Assay Kit (ThermoFisher Scientific cat. no. Q32851) on a Qubit 2.0 Fluorometer. Library fragment-length distributions were quantified with High Sensitivity D5000 Reagents (Agilent Technologies cat. no. 5067-5593) on the 4200 TapeStation System. The libraries were sequenced on an Illumina NextSeq 2000 instrument with single-end reads.

### Antibody staining for flow cytometry

For antibody staining of cell-surface proteins (VCAM1, TFRC), hiPSC-derived or primary mouse astrocytes were dissociated with Accutase, washed with DPBS, incubated with conjugated primary antibodies for 20 min on ice in DPBS + 1% BSA, washed with DPBS, and then resuspended in DPBS + 1% BSA for flow cytometry. For antibody staining of intracellular proteins (C3), hiPSC-derived astrocytes were dissociated with Accutase, washed with DPBS, fixed with 2% paraformaldehyde for 10 min at RT, washed twice with DPBS + 0.5% Tween 20 (Millipore Sigma cat. no. P9461), incubated with unconjugated primary antibody for 20 min at RT, washed with DPBS + 0.5% Tween 20, incubated with conjugated secondary antibody for 20 min at RT, washed with DPBS + 0.5% Tween 20, and then resuspended in DPBS + 0.5% Tween 20 for flow cytometry. For dual staining of VCAM1 and C3, the protocol for intracellular protein staining was used.

Conjugated primary antibodies:

PE-Cy7 mouse anti-VCAM1 (1:80 dilution; BioLegend cat. no. 305818)

PE-Cy7 mouse anti-TFRC (1:80 dilution; BioLegend cat. no. 334112)

Unconjugated primary antibodies:

Mouse anti-C3 (1:250 dilution; BioLegend cat. no. 846302)

Rabbit anti-VCAM1 (1:250 dilution; Abcam cat. no. ab134047)

Conjugated secondary antibodies:

AF488 goat anti-mouse IgG (1:1000 dilution; ThermoFisher Scientific cat. no. A-11029)

AF568 goat anti-rabbit IgG (1:1000 dilution; ThermoFisher Scientific cat. no. A-11036)

### Pooled CRISPRi screening

To identify transcriptional regulators of inflammatory reactivity, we created a custom sgRNA library targeting the human transcription factors^42^, using 5 sgRNAs with the highest predicted activity scores from Horlbeck *et al.*^45^ per gene. The library was created by cloning a pool of sgRNA-containing oligonucleotides custom-synthesized by Agilent Technologies into our optimized sgRNA expression vector as previously described^36^. To screen against the druggable genome, we used the H1 sub-library from Horlbeck *et al.*^45^. The transcription factor and druggable genome libraries were packaged into lentivirus as previously described^105^. For each experimental replicate, ∼10 million iAstrocytes were plated onto 4 Matrigel-coated 15-cm dishes, transduced with the lentiviral transcription factor or H1 sgRNA library so that 70-80% of cells were transduced, treated with vehicle control or IL-1α+TNF+C1q for 24 hours, and then incubated with pHrodo-labeled rat synaptosomes or stained for cell-surface VCAM1 (using the PE-Cy7 mouse anti-VCAM1 antibody) as shown in Fig. 3a. iAstrocytes were sorted into pHrodo high vs. low or VCAM1 high vs. low (top and bottom 30% of cells) using a BD FACSAria Fusion cell sorter at 5,000-10,000 events per second, and then pelleted for genomic DNA extraction. sgRNA abundances were then measured using next-generation sequencing as previously described^105^. The screens were performed with two experimental replicates per condition.

We chose to transduce iAstrocytes with sgRNA libraries at 70-80% transduction because we found that we could not enrich for sgRNA-transduced iAstrocytes with puromycin selection, which changed the morphology and behavior of iAstrocytes. Although a transduction rate of 70-80% is higher than the usual target of 30%, it is not problematic for the following reasons: (i) in this regime, most cells will only be transduced with 1 or 2 sgRNAs, (ii) the majority of sgRNAs do not have a phenotype because most genes don’t have a phenotype in the screen; thus, the rate of cells with 2 sgRNAs that both cause phenotypes is very low, (iii) the screens are carried out at very high representation, i.e. each sgRNA is represented in hundreds of independent cells. Since pairings of sgRNAs during double infections can be assumed to be random, each sgRNA will mostly occur by itself or in combination with sgRNAs that don’t cause a phenotype, and the few instances where it is paired with other sgRNAs that do have a phenotype constitute a negligibly small fraction that will not substantially affect the average phenotype measured in a pooled screen.

### CROP-seq

sgRNAs targeting the top hits from the CRISPRi screens and also candidate regulators selected based on literature were cloned into pMK1334. The concentration of each sgRNA plasmid was measured using the Qubit dsDNA HS Assay Kit on a Qubit 2.0 Fluorometer, and then the plasmids were pooled. iAstrocytes were transduced with lentivirus generated from the sgRNA pool at low multiplicity of infection so that <30% of cells were transduced, treated with vehicle control or IL-1α+TNF+C1q for 24 hours, and then sorted for sgRNA-transduced cells via FACS. Sorted iAstrocytes were then used as input for single-cell RNA-seq using Chromium Next GEM Single Cell 3’ v3.1 reagents (10X Genomics cat. no. PN-1000121, PN-1000127, and PN-1000213), loading ∼45,000 cells per reaction into four reactions, with two reactions for vehicle control-treated iAstrocytes and two reactions for IL-1α+TNF+C1q-treated iAstrocytes. To facilitate association of single-cell transcriptomes with sgRNAs, sgRNA-containing transcripts were amplified as described in Tian *et al.*^105^. The sgRNA-enrichment libraries were separately indexed and sequenced as spike-ins alongside the whole-transcriptome single-cell RNA-seq libraries on a NovaSeq 6000, recovering on average ∼29,000 transcriptome reads and ∼5,000 sgRNA-containing transcript reads per cell.

### Measurement of cytokine concentrations

Conditioned media was collected from iAstrocytes, spun down at 300 g for 10 min to remove debris, and then frozen at −80 °C until analysis. The concentration of selected cytokines (IFN-α2a, IFN-β, IFN-γ, IL-6, CXCL10, GM-CSF) was measured by multi-array electrochemiluminescence using custom U-PLEX plates from Meso Scale Discovery following the manufacturer’s instructions.

### Immunostaining of human neuropathological samples

#### Alzheimer’s disease

Brain tissue collection procedures were approved by the Institutional Review Board at Vanderbilt University Medical Center, and written consent for brain donation was obtained from patients or their surrogate decision makers. A diagnostic post-mortem evaluation was performed to confirm the presence of Alzheimer’s disease following the National Alzheimer’s Coordinating Center (NACC) Neuropathology Data Form. Tissue for this study was flash-frozen in liquid nitrogen at the time of donation. Cryosectioned tissues were fixed with 4% paraformaldehyde, photobleached with a broad-spectrum LED array for two days, and then processed for immunofluorescence staining. Samples were permeabilized with 0.3% Triton X-100 in PBS, blocked with 10% goat serum in PBS, and incubated at 4L overnight with the following primary antibodies: mouse anti-C3 (1:200 dilution; Biolegend cat. no. 846302), chicken anti-GFAP (1:300 dilution; Aves Labs cat. no. GFAP), rabbit anti-VCAM1 (1:200 dilution; Abcam cat. no. ab134047). The following day, after washing with PBS, immunostaining was completed by a 1 hour room temperature incubation with secondary antibodies (goat anti-rabbit Alexa Fluor 488, goat anti-mouse Alexa Fluor 546, and goat anti-chicken Alexa Fluor 647; 1:1,000 dilution; ThermoFisher Scientific). After additional washes, tissue sections were mounted with the anti-fade Fluoromount-G medium containing DAPI (Southern Biotechnology). Images were acquired with a Leica DMi8 epifluorescence microscope. 6-8 fields were collected per sample.

#### Hypoxic ischemic encephalopathy

All human HIE tissue was collected with informed consent and in accordance with guidelines established by UCSF Committee on Human Research (H11170-19113-07) as previously described^111^. Immediately after procurement, all brains were immersed in PBS with 4% paraformaldehyde for 3 d. On day 3, the brain was cut in the coronal plane at the level of the mammillary body and immersed in fresh 4% paraformaldehyde and PBS for an additional 3 d. After fixation, all tissue samples were equilibrated in PBS with 30% sucrose for at least 2 d. After sucrose equilibration, tissue was placed into molds and embedded with optimal cutting temperature medium for 30 min at room temperature followed by freezing in dry ice–chilled ethanol. UCSF neuropathology staff performed brain dissection and its evaluation. The diagnosis of HIE requires clinical and pathological correlation; no widely accepted diagnostic criteria are present for the pathological diagnosis of HIE. HIE cases showed consistent evidence of diffuse white matter gliosis, as evaluated by the qualitative increase in the number of GFAP-positive cells in addition to the increased intensity of GFAP staining. For immunostaining of reactive astrocyte markers, tissue slides were bleached with UV overnight, rinsed with PBS for 10 minutes, incubated with blocking solution (10% normal goat serum + 0.2% Triton X-100 in PBS) for 1 hour at room temperature, and then incubated with the following primary antibodies overnight at 4 L: mouse anti-C3 (1:200 dilution; Biolegend cat. no. 846302) + rabbit anti-GFAP (1:500 dilution; Agilent Dako cat. no. Z0334), or mouse anti-GFAP (1:500 dilution; Milipore Sigma cat. no. G3893) + rabbit anti-VCAM1 (1:200 dilution; Abcam cat. no. ab134047). Afterwards, the samples were rinsed three times with PBS + 0.2% Triton X-100 for 10 minutes each time, and then incubated with secondary antibodies for 1 hour at room temperature in the dark.

### Immunostaining of mouse tissue

#### Animals

All *in vivo* experiments were conducted according to protocols approved by the Chancellor’s Animal Research Committee of the Office for Protection of Research Subjects at the University of California, Los Angeles. Mice were housed in a 12 h light/dark cycle with controlled temperature and humidity and allowed *ad libitum* access to food and water. Wild-type C57BL/6J mice (RRID: IMSR_JAX:000664) were compared with transgenic mice with the *Stat3* gene conditionally deleted from astrocytes using a well-characterized 73.12 *mGfap*-Cre-*Stat3*-loxP mice on a C57BL/6J background^13^.

#### LPS-mediated neuroinflammation, euthanasia, and histology preparation

*E. coli*-derived lipopolysaccharide (LPS) (Sigma L5024, O127:B8; Lot 0000084086 with >500,000 endotoxin units per mg) was dissolved and diluted to 1 mg/ml in sterile PBS and stored in aliquots at −80°C. LPS (5 mg/kg) was administered to mice by intraperitoneal injection and mice were euthanized 24 h later. Young adult male and female mice were used between two and four months of age. After terminal anesthesia by barbiturate overdose, mice were perfused transcardially with 4% paraformaldehyde (Electron Microscopy Sciences 15714). Brains were removed, post-fixed 4 hours, and cryoprotected in buffered 30% sucrose for 48 h.

#### Immunohistochemistry

Embedding of fixed brains was accomplished by blotting brains dry, mounting into a cryomold (Epredia 2219) which was then filled with OCT compound (Sakura Finetek 4583). To snap freeze, cryomolds were partially submerged into a pool of 2-propanol cooled by a bed of dry ice. Brains were sectioned coronally at 40 µm on a cryostat (Leica CM1950) with a 34° MX35 Premier+ blade (Epredia 3052835) and stored in 0.05% NaN_3_ in PBS at 4°C. Sections were moved to a 48 well plate, washed with PBS, and blocked for 1 h with blocking buffer: 5% goat serum (Gibco 16210-064), 1% bovine serum albumin (Sigma A7906), 0.03% Triton X-100 (Sigma T8787) in PBS. The brains were incubated overnight at 4°C in blocking buffer with primary antibodies against ISG15 (rabbit, clone: 1H9L21, Invitrogen 703132, 1:100), C3 (rat, clone: 11H9, Abcam ab11862, 1:50), and GFAP (chicken, polyclonal, Aves Labs GFAP, 1:500). Sections were washed three times with PBS and incubated for 2 h at room temperature with secondary antibodies in blocking buffer: goat anti-rabbit Alexa Fluor 488 (Invitrogen A-11034, 1:1,000), goat anti-rat Alexa Fluor 555 (Invitrogen A-21434, 1:1,000), and goat anti-chicken Alexa Fluor 647 (Invitrogen A-21449, 1:1,000). Sections were washed three times with PBS and moved to glass slides (Fisher 12-550-15). After PBS was removed, Fluoromount-G with DAPI (Invitrogen 00-4959-52) was added and a coverslip (Globe Scientific 1415-15) applied. Slides were dried at room temperature in dark overnight. All sections imaged were located between bregma −1.22 and −2.70.

#### Microscopy and Quantification

Mouse brains were imaged on a Leica DMi8 microscope coupled with a Yokagawa CSU-W1 SoRa spinning disk confocal scanner unit using a 20x/0.80NA air or 100x/1.47NA oil immersion objective lens, 200-350 ms exposure, 12-bit sensitivity, 1×1 binning, 1.5-2.5 laser intensity with 401 nm, 487 nm, 561 nm, and 639 nm excitation lasers, running Micro-Manager version 2.0.1 software (Weill Innovation Core, UCSF Center for Advanced Light Microscopy). Mouse brains were also imaged with a Zeiss Axio Scan.Z1 with a Zeiss Colibri 7 unit, 20x/0.8NA objective lens, 5-30 ms exposure, 1×1 binning, 25%-100% intensity using 425 nm, 495 nm, 570 nm, and 655 nm lasers, running ZEN version 2.6 software. Quantification was performed on the slide scanner images (see “Data analysis” section below).

### Data analysis

#### Pooled CRISPRi screens

We analyzed the data from the pooled CRISPRi screens using the MAGeCK-iNC bioinformatic pipeline previously described in Tian *et al.*^105^. Briefly, reads from next-generation sequencing were cropped and aligned to the reference using Bowtie to determine sgRNA counts in each sample. Next, counts files of samples subject to comparison were input into MAGeCK^112^ and log_2_ fold changes (LFCs) and *P* values were calculated for each sgRNA using the ‘mageck test – k’ command, which takes into account variance among replicates. Following that, gene level knockdown phenotype scores were determined by averaging LFCs of the top 3 sgRNAs targeting a given gene with the most significant *P* values. The *P* value associated with each gene was determined by comparing the set of *P* values for sgRNAs targeting that gene with the set of p values for non-targeting control sgRNAs using the Mann-Whitney U test. To call hit genes, we first generated a null distribution by performing random sampling of 5 (the number of sgRNAs against each gene in the library) with replacement from non-targeting control sgRNAs to generate ‘negative-control-quasi-genes’ and calculated knockdown phenotype scores and *P* values for each of them. Then, we calculated the hit strength (gene score), defined as the product of knockdown phenotype score and –log (*P* value), for all genes in the library and for ‘negative-control-quasi-genes’ generated above. Based on the distribution of the gene scores, a cutoff value was chosen so that less than 10% of hit genes were negative control quasi-genes.

### Master regulator analysis (MRA)

To collect human astrocyte RNA-seq data for co-expression network reconstruction, we downloaded gene-level counts from samples annotated as “Astrocyte” in ARCHS4 ^113^. For published human astrocyte RNA-seq datasets not found in ARCHS4, we manually downloaded FASTQ files from GEO, and then processed the FASTQ data using Elysium^114^, which implements the alignment pipeline used in ARCHS4. In addition to published human astrocyte RNA-seq datasets, we also used Elysium to process additional human astrocyte RNA-seq datasets from our labs (Ye Zhang lab, UCLA^41^, and Kampmann lab; all datasets used for MRA available at https://kampmannlab.ucsf.edu/inflammatory-reactive-astrocyte-analysis). We merged the gene-level count matrices from the above sources into a single matrix, which was then used for MRA (see Supplementary Table 11 for the metadata corresponding to all samples used for co-expression network reconstruction). Batch correction was first performed across datasets using ComBat-seq^115^, which is a part of the R (version 4.0.3) package “sva” (version 3.38.0). Gene-level counts were then transformed to log-scale using the variance-stabilizing transformation in DESeq2 (version 1.30.1). A list of human transcription factors (TFs) was obtained from Lambert *et al.*^42^, and a list of human kinases and phosphatases was obtained from Manning *et al.*^43^ and Liberti *et al.*^44^. Genes and regulators (TFs, kinases and phosphatases) with low expression in hiPSC-derived astrocytes used in our study (mean transcripts per million < 1) were removed from the matrix. MRA was then performed using the R package “RTN” (version 2.16.0)^38–40^, following the RTN vignette available on Bioconductor.

### CROP-seq

Alignment and quantification were performed on 10X single-cell RNA-seq libraries and sgRNA-enriched libraries using Cell Ranger (version 5.0.1) with default parameters and reference genome GRCh38-3.0.0. Cellranger aggr was used to aggregate counts belonging to the same sample across different GEM wells. sgRNA unique molecular identifier (UMI) counts for each cell barcode were obtained using the pipeline described in Hill *et al.*^116^. sgRNAs were assigned to cells using demuxEM^117^. The gene vs. cell barcode matrix outputted by Cell Ranger was converted into a SingleCellExperiment (SCE) object using the read10xCounts function from the DropletUtils R package (version 1.10.388). sgRNA assignments were appended to the SCE metadata and filtered to only include cells with a single sgRNA. The SCE object was then converted into a Seurat object for subsequent analysis. Data normalization, log-transformation, and identification of highly variable genes were performed using Seurat::SCTransform^118^.

For exploratory analysis of the effect of knocking down each regulator targeted in the CROP-seq experiment, a separate Seurat object was created for each regulator consisting of iAstrocytes transduced with a non-targeting (NTC) sgRNA and iAstrocytes transduced with the sgRNA targeting that regulator. For each of these Seurat objects, the data was renormalized with SCTransform, dimensionality reduction was performed with Seurat::RunPCA (30 PC’s retained) and Seurat::RunUMAP, and clustering was performed with Seurat::FindNeighbors and Seurat::FindClusters (resolution = 0.5). Regulators whose knockdown resulted in clear spatial separation between knockdown iAstrocytes and NTC iAstrocytes in UMAP were selected for further analysis. The PCA embeddings from the Seurat objects corresponding to the selected regulators were used for aligned UMAP^119^, which allows the effect of knocking down different regulators to be visualized in the same UMAP embedding. This aligned UMAP embedding was used for Fig. 4a,b,d and Fig. 5c, which displayed only NTC iAstrocytes. Markers of IRAS1 and IRAS2 in NTC iAstrocytes were identified by performing Student’s t-tests on the Pearson residuals from SCTransform using Seurat::FindMarkers.

To find genes whose differential expression induced by IL-1α+TNF+C1q treatment was changed by regulator knockdown, we used the R package limma (version 3.46.0) to perform linear regression on the Pearson residuals from running SCTransform on the Seurat object containing all iAstrocytes assigned with a single sgRNA. We used the design formula y ∼ regulatorKnockdown + cytokineTreatment + regulatorKnockdown:cytokineTreatment, where the interaction term regulatorKnockdown:cytokineTreatment reflects the effect of knocking down a given regulator on the differential expression induced by IL-1α+TNF+C1q treatment. For each regulator knockdown, genes with a statistically significant interaction term were extracted for the analysis presented in Extended Data Fig. 4d-e.

To construct the heatmap in Extended Data Fig. 4e, the log_2_-fold-change (LFC) values of the above DEGs were weighted by their associated multiple testing-corrected *P*-values (*P*_adj_) to generate a gene score, where gene score = LFC * (1 – *P*_adj_). Hierarchical clustering (Ward’s method) was then performed on both the rows (genes) and columns (regulators) of the gene score matrix, using Pearson correlation as the distance metric. The heatmap was drawn using R package ComplexHeatmap (version 2.6.2)^120^.

### Integration of iAstrocyte single-cell RNA-seq data with external single-cell RNA-seq datasets

UMI count matrices of single-cell RNA-seq data from Barbar *et al.*^29^ were downloaded from Synapse (syn21861229). Processed single-cell RNA-seq data objects corresponding to data from Wheeler *et al.*^83^ were requested from Zhaorong Li and Michael Wheeler. UMI count matrices of single-cell RNA-seq data from Hasel *et al.*^58^ were downloaded from GEO (GSE148611). Integration of NTC iAstrocytes from the CROP-seq experiment with the above previously published single-cell RNA-seq datasets was performed by first renormalizing each dataset with Seurat::SCTransform and re-running Seurat::RunPCA, then selecting features for integration using Seurat::SelectIntegrationFeatures (nfeatures = 3000) and Seurat::PrepSCTIntegration, and then performing data integration using reciprocal PCA with Seurat::FindIntegrationAnchors (normalization.method = ‘SCT’, reduction = ‘rpca’) and Seurat::IntegrateData (normalization.method = ‘SCT’), using NTC iAstrocytes as the reference dataset. The integrated PCA embeddings were then used for clustering (resolution = 0.4) and UMAP. For integration with Barbar *et al.*^29^, the k.anchor parameter of Seurat::FindAnchors was set to 20; for Wheeler *et al.*^83^, k.anchor was set to 50; for Hasel *et al.*^58^, k.anchor was set to 100. Seurat::AddModuleScore was used to generate module expression scores for IRAS1 vs. IRAS2 markers in astrocytes from external single-cell datasets.

### Bulk RNA-seq of hiPSC-derived astrocytes generated in this study

Alignment and quantification were performed using Salmon (version 1.4.082), with the -- noLengthCorrection flag and an index generated from the human transcriptome (GRCh38, Gencode release 37). The R package Tximport (version 1.18.083) was used to obtain gene-level count estimates. Genes with zero counts across all samples were removed from the analysis. Differential gene expression analysis was performed with DESeq2 (version 1.30.1).

### Reanalysis of external bulk RNA-seq datasets

For Guttenplan *et al.*^9^, we downloaded the table of DEGs induced by IL-1α+TNF+C1q in immunopanned astrocytes from wild-type mice provided in GSE143598. We used BioJupies^121^ to reanalyze bulk RNA-seq data and obtain differentially expressed genes from Perriot *et al.*^122^ (GSE120411; hiPSC-derived astrocytes treated with IL1β+TNF), Barbar *et al.*^29^ (syn21861229; CD49f+ astrocytes sorted from cerebral organoids treated with vehicle control or IL-1α+TNF+C1q), and Anderson *et al.*^51^ (GSE76097; astrocyte-specific RNA from *Stat3* cKO vs. wild-type mice subject to spinal cord injury). For Anderson *et al.*, sample GSM1974209 was removed because it was an outlier in PCA visualization of the samples.

### Pathway enrichment analysis

We used Enrichr^123–125^ to perform enrichment analysis of gene lists. For display of enrichment results against the TRRUST gene set library, terms corresponding to mouse gene sets were removed for genes lists derived from human astrocytes; for genes lists derived from mouse astrocytes, terms corresponding to human gene sets were removed.

### Flow cytometry

Data from flow cytometry experiments were analyzed using FlowJo (version 10.7.1). Live cells were gated by plotting SSC-A vs. FSC-A and then single cells were gated by plotting FSC-H vs. FSC-A. For experiments involving CRISPRi knockdown, analysis was restricted to sgRNA-transduced cells (gating on the histogram of BFP fluorescence values). For antibody staining experiments where median fluorescence intensity (MFI) values were reported, the average MFI of unstained control samples were subtracted from the MFI of stained samples.

### Image segmentation and analysis

We used CellProfiler (v3.15)^126^ to segment and quantify neuron viability, neuron GCaMP recordings, and iAstrocyte immunostaining images. For neuron viability, monochrome images of Hoechst and TO-PRO3 were segmented to nuclei and TO-PRO3+ objects, respectively; TO-PRO3+ neurons (i.e. dead neurons) were determined by overlap of nuclei with TO-PRO3+ objects using the RelateObjects module in CellProfiler. For neuron GCaMP recordings, neuronal soma were segmented on the maximum intensity projection of all GCaMP images in the recording, nuclei were segmented on a single BFP image collected at the end of recording, and only nuclei that were contained within a GCaMP+ soma were retained using the RelatedObjects and FilterObjects modules in CellProfiler. The median GCaMP intensity per soma at each time point was then extracted for analysis. For each neuronal soma, ΔF was calculated as F_t_ − F_t-1_ and F_0_ was calculated as the average of a 10-second rolling window around F_t_.

For immunostaining of astrocyte markers, the total image intensity of GFAP, S100β, GLAST, NFIA, Cx43, glutamine synthetase, or vimentin was calculated and divided by the total image intensity of Hoechst to normalize for cell number. For immunostaning of C3 and IFIT3, nuclei were segmented from monochrome images of Hoechst followed by segmentation of cell boundaries from monochrome images of Na^+^/K^+^ ATPase. The integrated intensity of C3 and IFIT3 was then calculated on a per-cell basis. Thresholds to define C3+ and IFIT3+ cells based on integrated C3 and IFIT3 intensity were determined from samples treated with vehicle control.

For analysis of images from immunostaining of human neuropathological tissue, monochrome images of DAPI, VCAM1, C3, and GFAP staining were converted to probability maps of nuclei, VCAM1+ objects, C3+ objects, and GFAP+ objects respectively using ilastik^127^; annotations for representative nuclei, VCAM1+ objects, C3+ objects, and GFAP+ objects were inputted manually to train the classifier. The probability maps were then inputted into CellProfiler to segment nuclei, VCAM1+ objects, C3+ objects, and astrocytes (defined as nuclei-containing GFAP+ objects), and the VCAM1 or C3 staining status of astrocytes was determined by overlap of astrocytes with VCAM1+ or C3+ objects using the RelateObjects module in CellProfiler.

For analysis of images from immunostaining of mouse tissue, images from the Axio Scan.Z1 were imported into QuPath (v0.3.2)^128^. The corpus callosum or hippocampus were outlined, and then nuclei were segmented on the DAPI channel using the “Cell detection” module, with virtual cell boundaries created by expansion of the nuclei segmentation. Classifiers were then created to distinguish Gfap+, C3+, and Isg15+ cells by thresholding on the maximum intensity of the respective markers contained within the virtual cell boundary. The classifiers were then applied sequentially to obtain classify astrocytes (GFAP+ cells) as Isg15−/C3+, Isg15+/C3−, or Isg15+/C3+.

### Beta regression, linear regression, and multiple testing adjustment of *P* values

When multiple comparisons needed to be made in a single experiment, we used beta regression for data consisting of percentages and linear regression for fluorescence values, cytokine concentrations, etc. For beta regression, which is appropriate for data ranged from 0 to 1, we first converted percentages to proportions. In cases where data occupied the extremal values of 0 or 1, we applied the transformation (y · (*n* − 1) + 0.5)/*n*, where *n* is the sample size, as previously proposed by Smithson & Verkuilen^129^. We then calculated *P* values using the betareg package in R (v3.1-4)^130^, using a logit link function (link = ‘logit’) and the bias-corrected maximum likelihood estimator (type = ‘BR’). For linear regression, we used the function lm in R with parameters set to default values. Where appropriate, *P* values obtained via beta or linear regression were adjusted for multiple hypothesis testing with Holm’s method using the function p.adjust in R (method = ‘Holm’).

### Statistics and reproducibility

Sample sizes were determined by referencing existing studies in the field. Major findings were validated using independent samples and orthogonal approaches. Numbers of replicates are listed in each figure. For the mouse experiments, mice of each genotype were randomized to LPS treatment.

---

**Extended Data Fig. 1.**
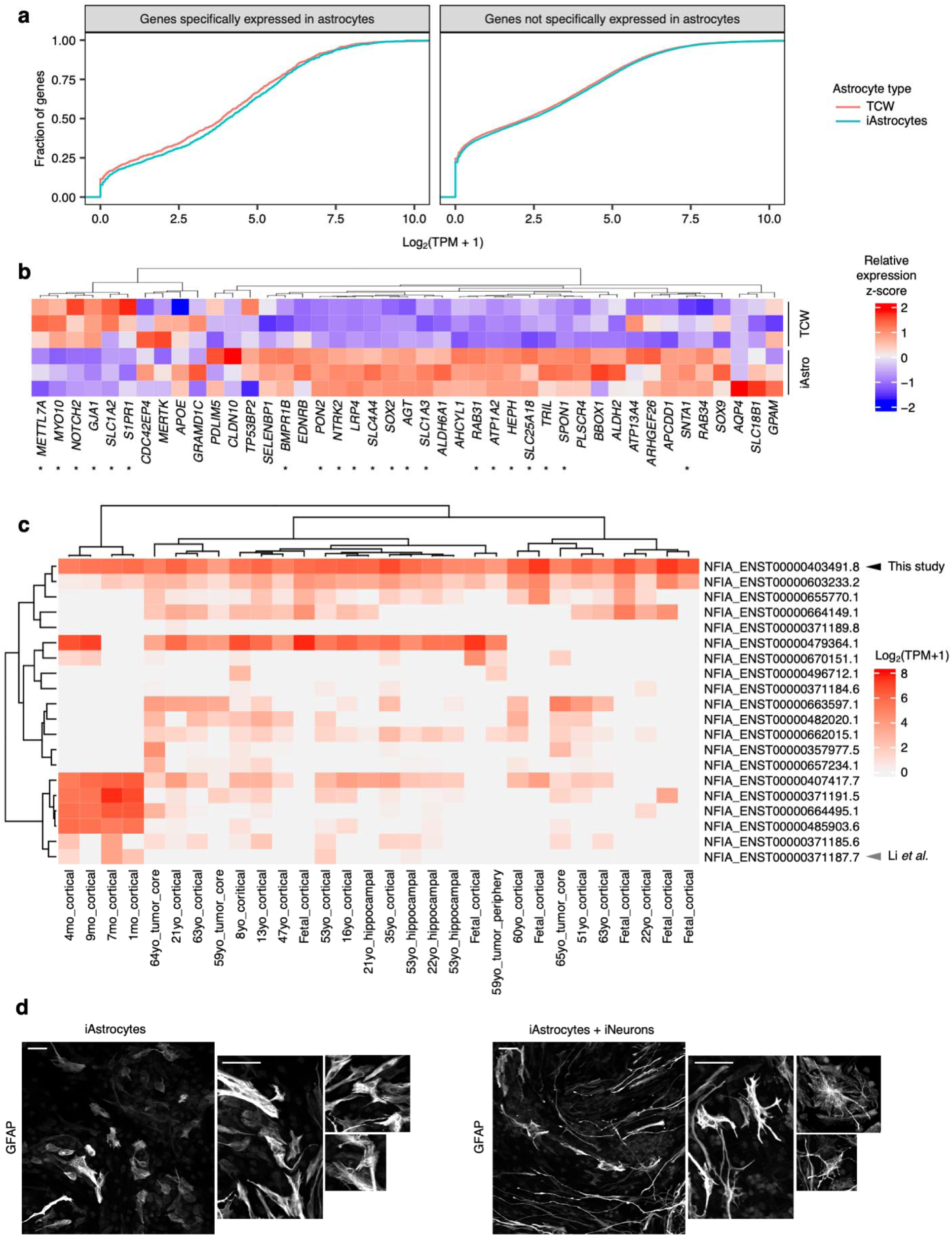
iAstrocytes express higher levels of astrocyte-specific genes compared to iPSC-derived astrocytes generated using the protocol from TCW *et al.*^18^. **a**, Empirical cumulative distribution functions of the mean expression of genes (averaged across experimental replicates, *n* = 3 wells) with astrocyte-specific expression (astrocyte fidelity > 40) or without astrocyte specific expression (astrocyte fidelity < 40) in iAstrocytes vs astrocytes generated using the TCW *et al*. protocol (TCW astrocytes). Genome-wide astrocyte fidelity scores were obtained from Kelley *et al.* TPM: transcripts per million. **b**, Relative expression (z-scored) of the top 50 genes with the highest astrocyte fidelity scores from Kelley *et al.*^133^ (organized by hierarchical clustering, see Methods) in iAstrocytes vs. TCW astrocytes (*n* = 3 experimental replicates corresponding to heatmap rows). Genes with statistically significant differential expression (adjusted *P* value < 0.1) between iAstrocytes and TCW astrocytes are marked with asterisks. **c**, Heatmap of log-scaled transcripts per million (TPM) values of NFIA transcripts in human primary astrocytes from Zhang *et al.*^92^ **d**, Representative images from immunostaining of GFAP in iAstrocytes cultured alone or with iNeurons. In each case, an entire field of view is displayed (left) next to magnified sections (all at same scale) containing representative astrocyte morphologies (right). Scale bars correspond to 60 μm.

**Extended Data Fig. 2.**
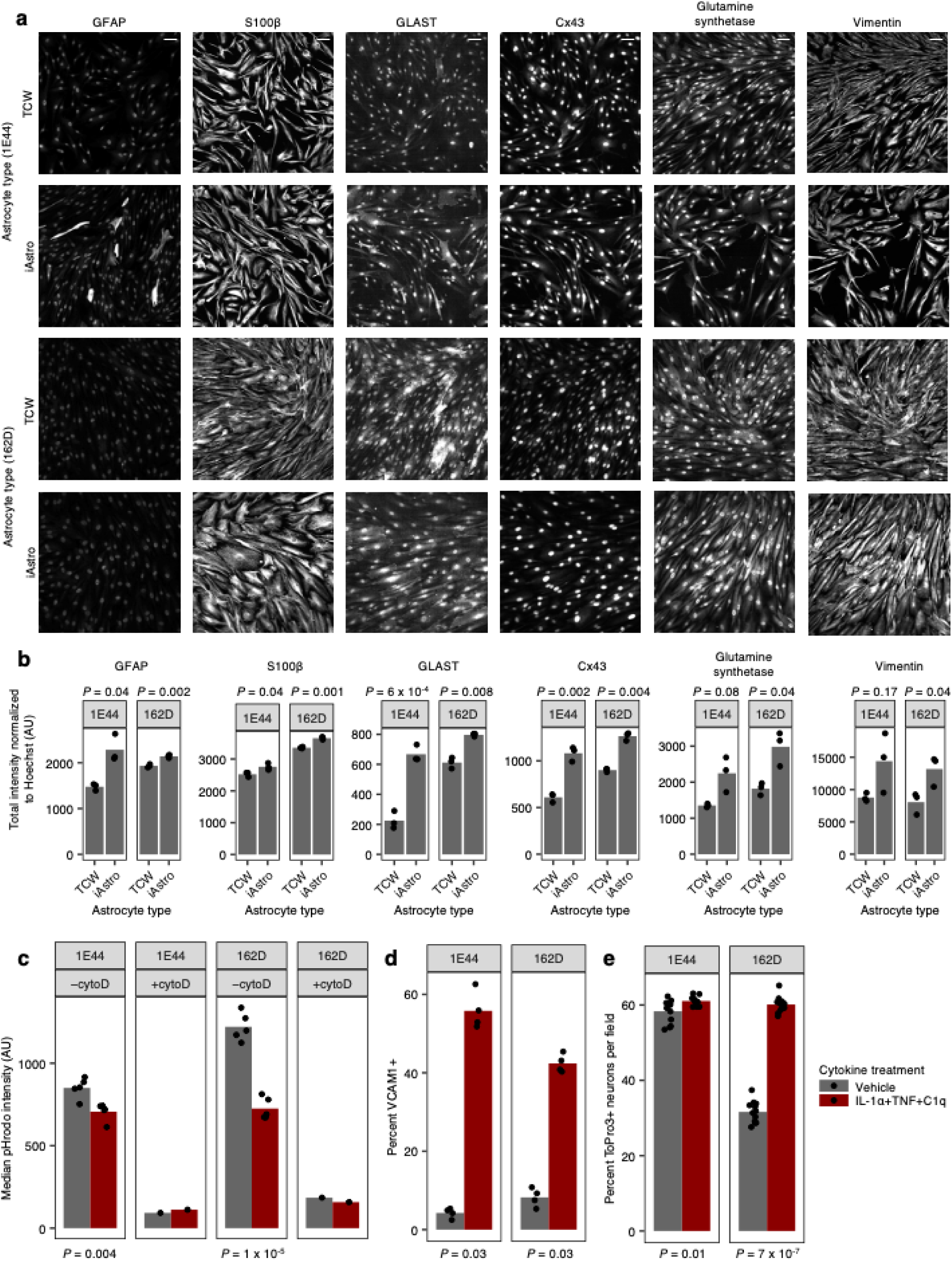
Validation of iAstrocyte differentiation from two additional hiPSC lines. **A**, Representative images of immunofluorescence against GFAP, S100β, GLAST, Cx43, glutamine synthetase, or vimentin in iAstrocytes vs. TCW astrocytes derived from TCW-1E44 or 162D hiPSCs (scale bar: 60 μm). **b**,Quantification of GFAP, S100β, GLAST, Cx43, glutamine synthetase, or vimentin immunofluorescence intensity (*n* = 3 wells). **c**, Phagocytosis of pHrodo-labeled rat synaptosomes (median fluorescence intensity measured by flow cytometry) by iAstrocytes derived from TCW-1E44 or 162D hiPSCs in the absence (*n* = 5 wells) or presence (*n* = 1 well) of cytochalasin D (cytoD). **d**, Percent VCAM1+ cells in TCW-1E44 or 162D iAstrocytes treated with vehicle control vs. IL-1α+TNF+C1q (*n* = 4 wells). **e**, Percentage of dead cells (measured by TO-PRO-3 permeability) in iNeurons incubated with conditioned media from TCW-1E44 or 162D iAstrocytes treated with vehicle control or IL-1α+TNF+C1q (*n* = 12 wells). In panels **b** and **c**, *P* values were calculated using the two-sided Student’s t-test. In panels **d** and **e**, *P* values were calculated using the two-sided Mann-Whitney U-test.

**Extended Data Fig. 3.**
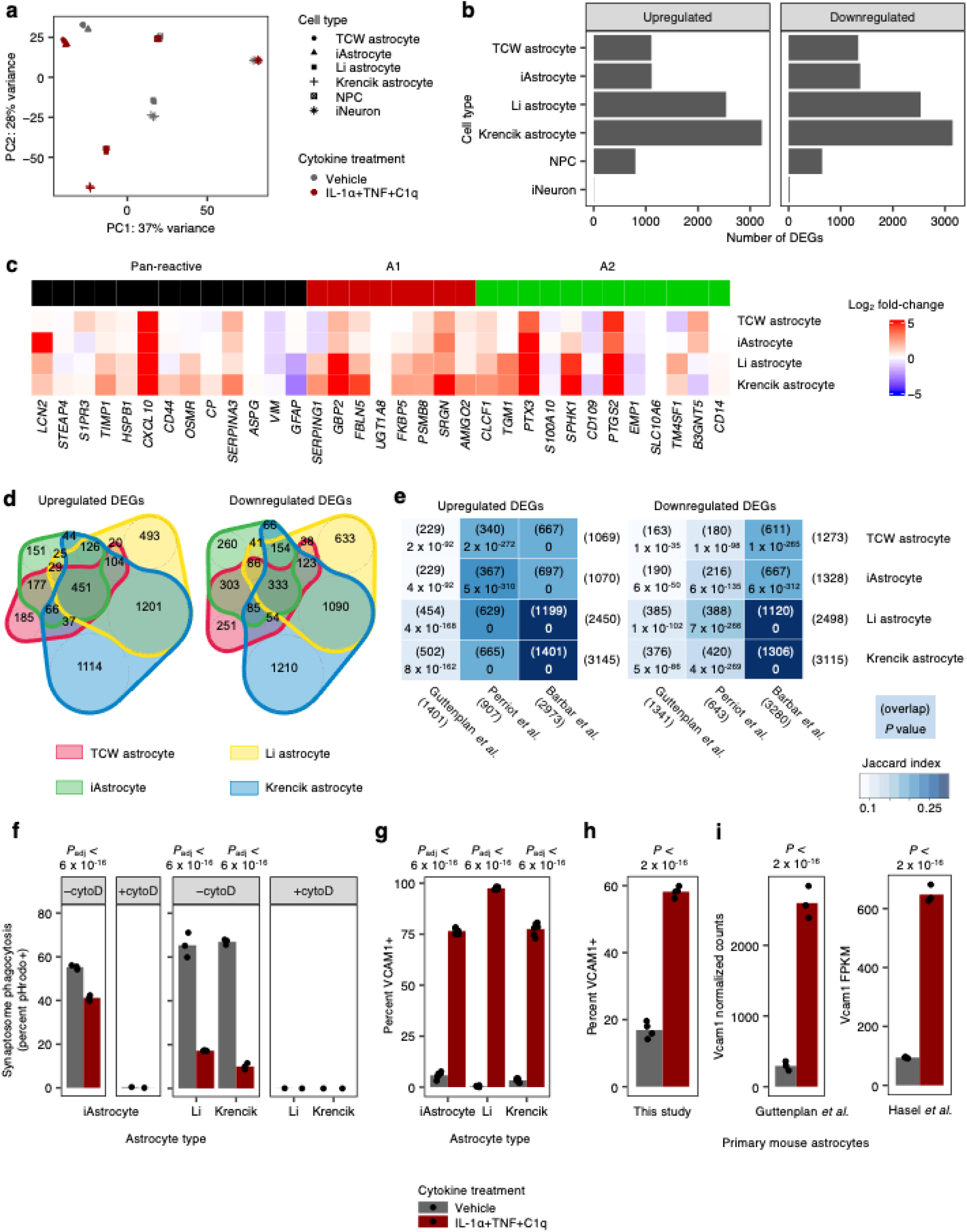
iAstrocytes respond to IL-1α+TNF+C1q in a highly similar manner as hiPSC-derived astrocytes generated using different protocols and primary mouse astrocytes. **a**, Principal component (PC) analysis plot of the gene expression profiles (top 5000 variable genes) of iAstrocytes vs. astrocytes derived using the protocols from TCW *et al.*^18^ (TCW astrocytes), Li *et al.*^19^ (Li astrocytes), or Krencik *et al.*^15^ (Krencik astrocytes), as well as iPSC-derived neurons (iNeurons) and neural progenitor cells (NPCs), treated with vehicle control or IL-1α+TNF+C1q (*n =* 3 wells for astrocyte samples, *n* = 2 wells for iNeuron and NPC samples). **b**, Number of differential expressed genes (DEGs) induced by IL-1α+TNF+C1q. **c**, Log_2_-fold-changes of pan-reactive, A1 reactive, and A2 reactive genes defined in Liddlelow *et al*.^8^ in hiPSC-derived astrocytes from this study. **d**, Overlap of upregulated and downregulated DEGs induced by IL-1α+TNF+C1q among hiPSC-derived astrocytes from this study. **e**, Overlap of upregulated and downregulated DEGs induced by IL-1α+TNF+C1q from hiPSC-derived astrocytes from this study compared to DEGs from inflammatory reactive astrocytes in other studies. **f**-**g**, Phagocytosis of pHrodo-labeled synaptosomes (**f**; *n* = 3 wells for −cytoD, *n* = 1 well for +cytoD) or induction of cell-surface VCAM1 (**g**; *n* = 6 wells) by iAstrocytes compared to Li and Krencik astrocytes. cytoD: cytochalasin D. **h**-**i**, Induction of VCAM1 expression by IL-1α+TNF+C1q in primary mouse astrocytes measured by flow cytometry in this study (**h**; *n* = 4 wells) or primary mouse astrocytes measured by RNA-seq in Guttenplan *et al.*^9^ and Hasel *et al.*^58^ (**i**; *n* = 3 mice). In panels **f** and **i**, *P* values were calculated using the two-sided Student’s t-test. In panels **g** and **h**, *P* values were calculated using beta regression (see Methods).

**Extended Data Fig. 4.**
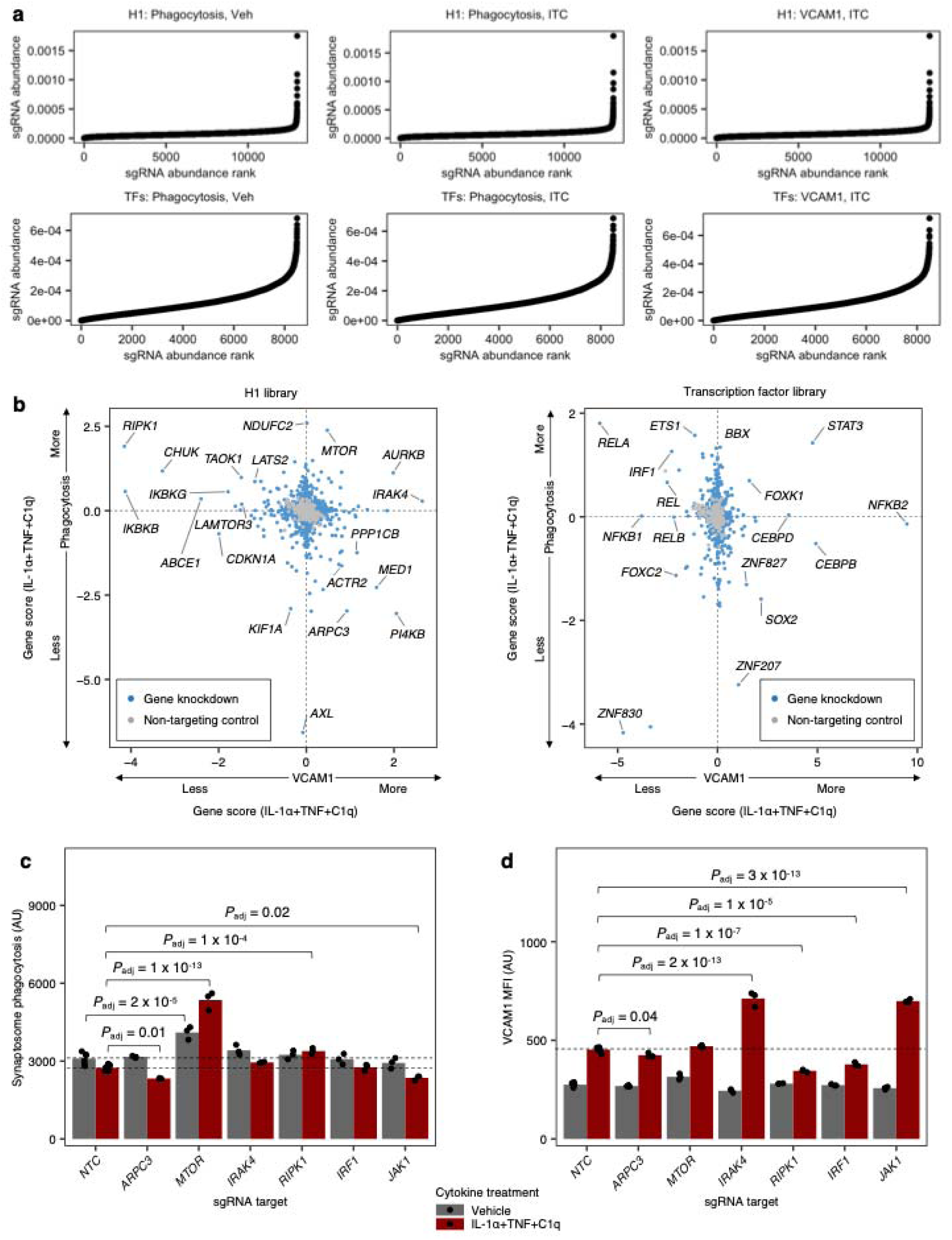
sgRNA abundance distribution in CRISPRi screens, comparison of phenotypes from VCAM1 and phagocytosis CRISPRi screens, and validation of selected hits from CRISPRi screens. **a**, sgRNA abundance distribution for the CRISPRi screens shown in Fig. 3. **b**, Gene scores (see Methods) from the phagocytosis vs. VCAM1 CRISPRi screen against transcription factors (left) or the druggable genome (right) in iAstrocytes treated with IL-1α+TNF+C1q. **c**-**d**, Phagocytosis of pHrodo-labeled synaptosomes **(c**) or induction of cell-surface VCAM1 **(d**) by iAstrocytes transduced with non-targeting sgRNA (NTC) vs. sgRNAs targeting selected hits from the screens shown in Fig. 3, treated with vehicle control or IL-1α+TNF+C1q (*n* = 6 wells for NTC, *n* = 3 wells for knockdowns). MFI: median fluorescence intensity measured by flow cytometry. In panels **c** and **d**, *P* values were calculated by linear regression (see Methods) and adjusted for multiple testing (*P*_adj_; Holm’s method) per family of tests (all comparisons within a plot).

**Extended Data Fig. 5.**
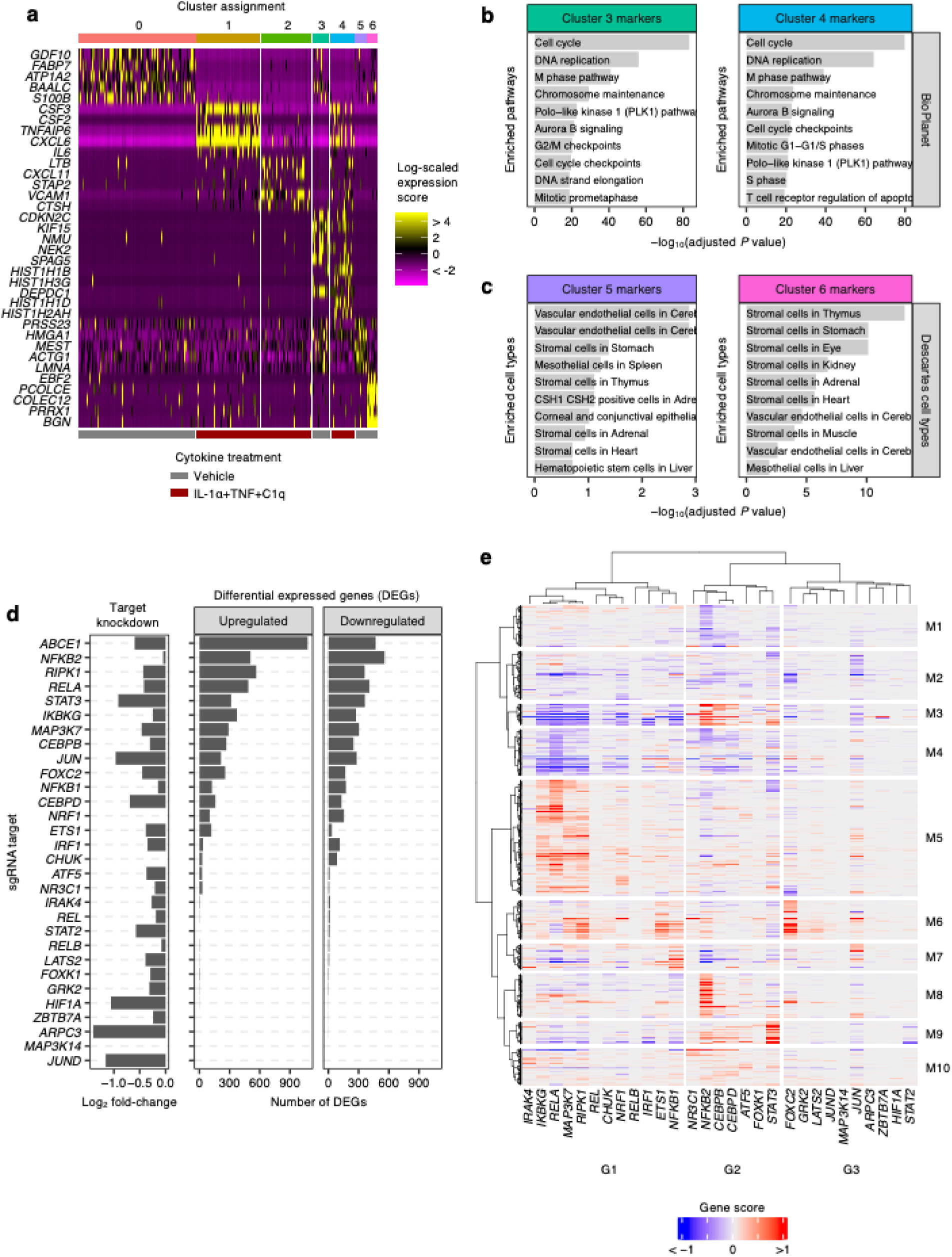
Additional analyses of CROP-seq data. **a**, Expression levels of the top cluster markers of non-targeting control (NTC) sgRNA-transduced iAstrocytes shown in Fig. 4a. **b**-**c**, Cellular pathway (BioPlanet^134^) enrichment analysis of Cluster 3 and 4 markers (**b**) and cell type marker (Descartes^135^) enrichment analysis of Cluster 5 and 6 markers (**c**) of NTC sgRNA-transduced iAstrocytes shown in Fig. 4a. **d**, The degree of regulator knockdown (left) or the number of differentially expressed genes (DEGs) whose differential expression induced IL-1α+TNF+C1q is significantly altered by regulator knockdown. **e**, Hierarchical clustering of the *P*-value-weighted log-fold-changes (gene score) of the union of knockdown-associated DEGs from panel **d**; DEGs associated with *ABCE1* knockdown were excluded due to a significant number of DEGs also being caused by *ABCE1* knockdown in vehicle control-treated iAstrocytes.

**Extended Data Fig. 6.**
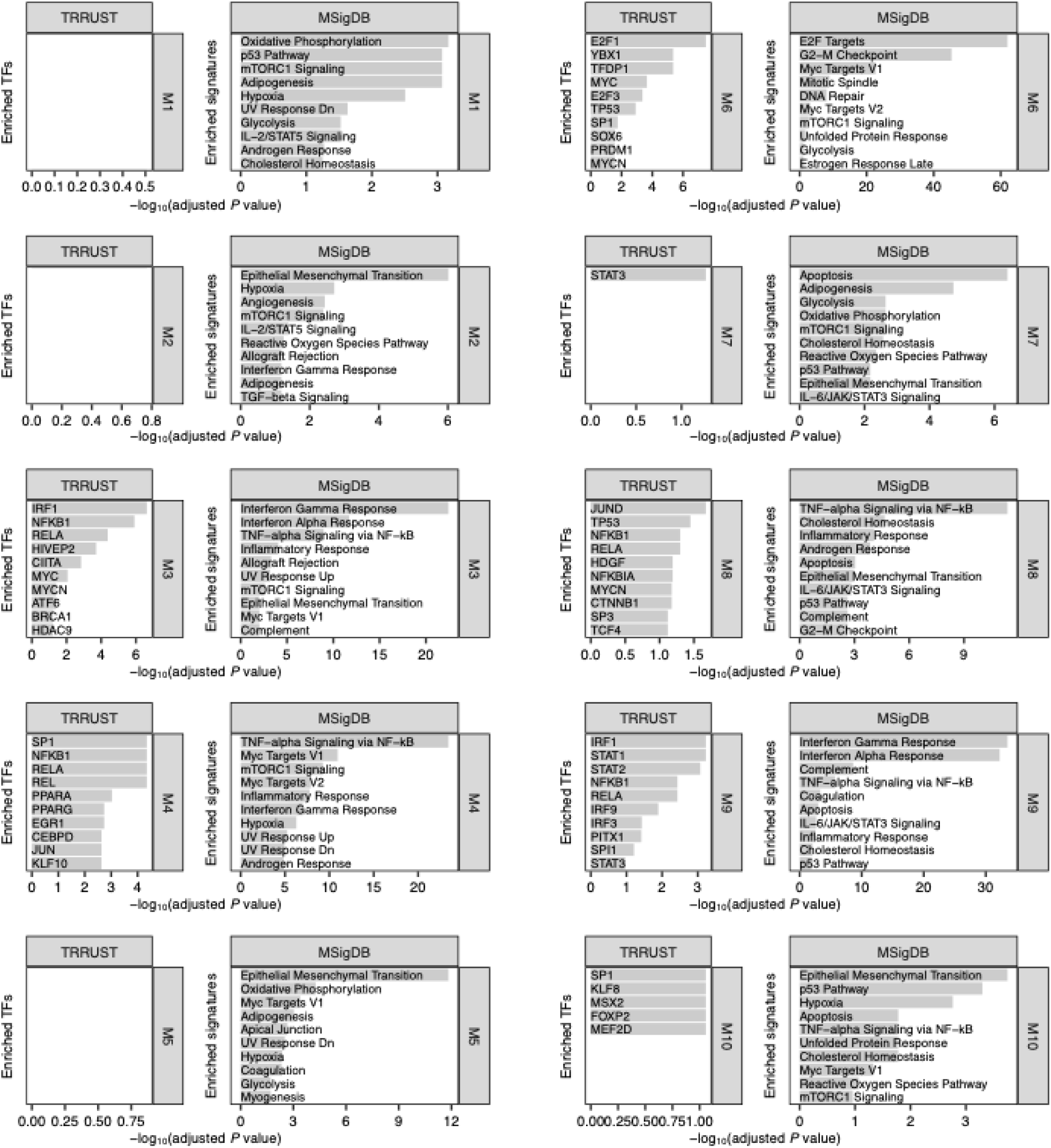
Enrichment analysis of CROP-seq knockdown-associated gene modules. Cellular pathway (MSigDB^131^) and upstream transcription factor (TRRUST^132^) enrichment analysis of gene modules from Extended Data Fig. 4e; TF – transcription factor.

**Extended Data Fig. 7.**
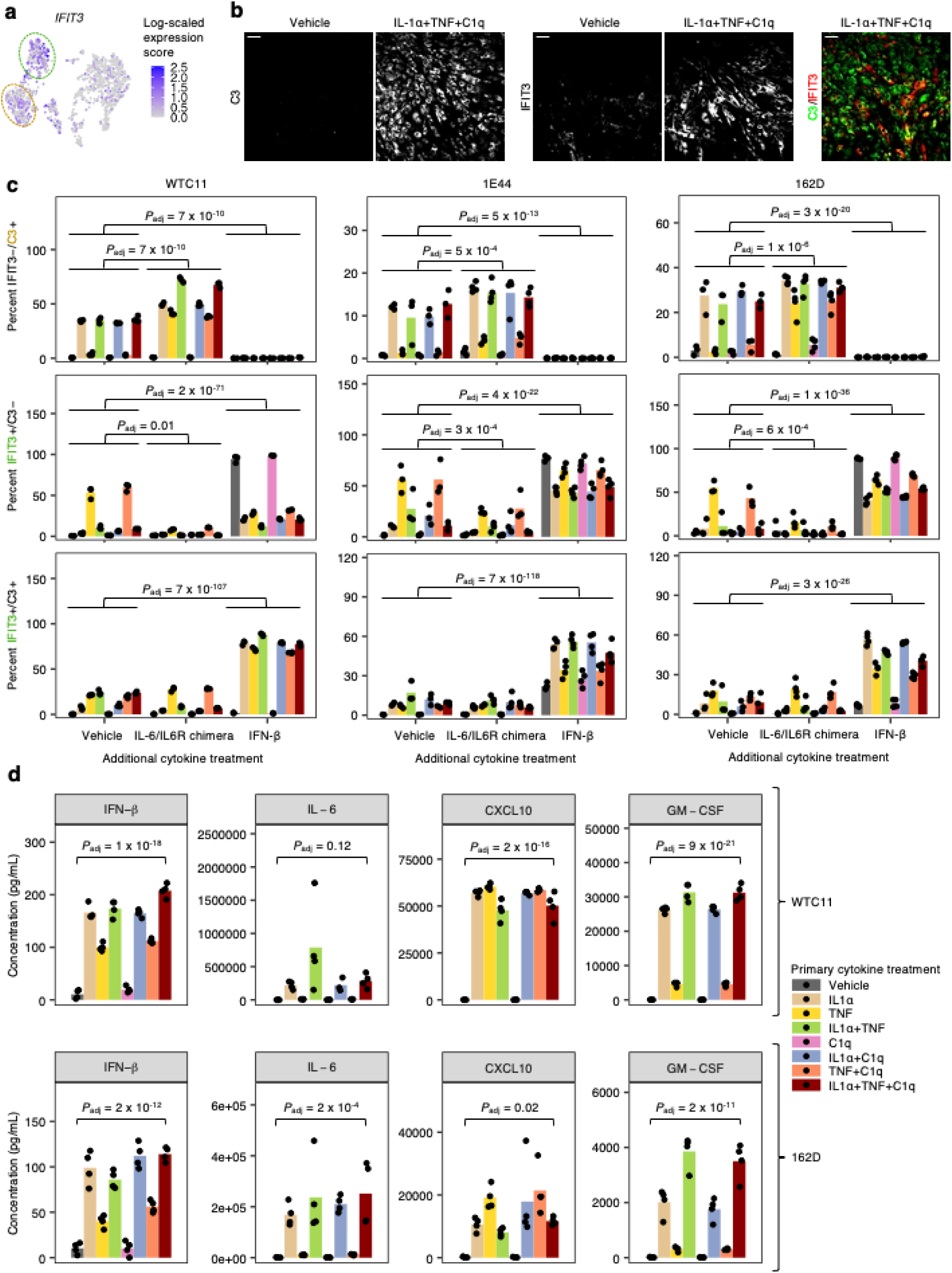
C3 and IFIT3 expression and cytokine production in iAstrocytes derived from multiple hiPSC lines. **a**, Transcript levels of *IFIT3* overlaid onto the UMAP embedding from Fig. 3a. **b**, Representative immunofluorescence images of C3 and IFIT3 staining (scale bar: 60 μm). **c**, Percent IFIT3−/C3+, IFIT3+/C3−, or IFIT3+/C3+ cells measured by immunofluorescence in iAstrocytes derived from multiple hiPSC lines (WTC11, TCW-1E44, 162D) treated with vehicle control vs. all possible combinations of IL-1α, TNF, and C1q, in the absence or presence of additional IL-6/IL6R chimera (25 ng/mL) or IFN-β (5 ng/mL) added concurrently (*n* = 3-4 wells). **D**, Concentration of IFN-β, IL-6, CXCL10, or GM-CSF in conditioned media from iAstrocytes derived from multiple hiPSC lines (WTC11, 162D) treated with vehicle control vs. all possible combinations of IL-1α, TNF, and C1q (*n* = 4 wells). For panels **c a**, *P* values were calculated using beta regression. For panel **d**, *P* values were calculated using linear regression (see Methods). *P* values were adjusted for multiple testing (*P*_adj_; Holm’s method) per family of tests (all comparisons within a plot).

**Extended Data Fig. 8.**
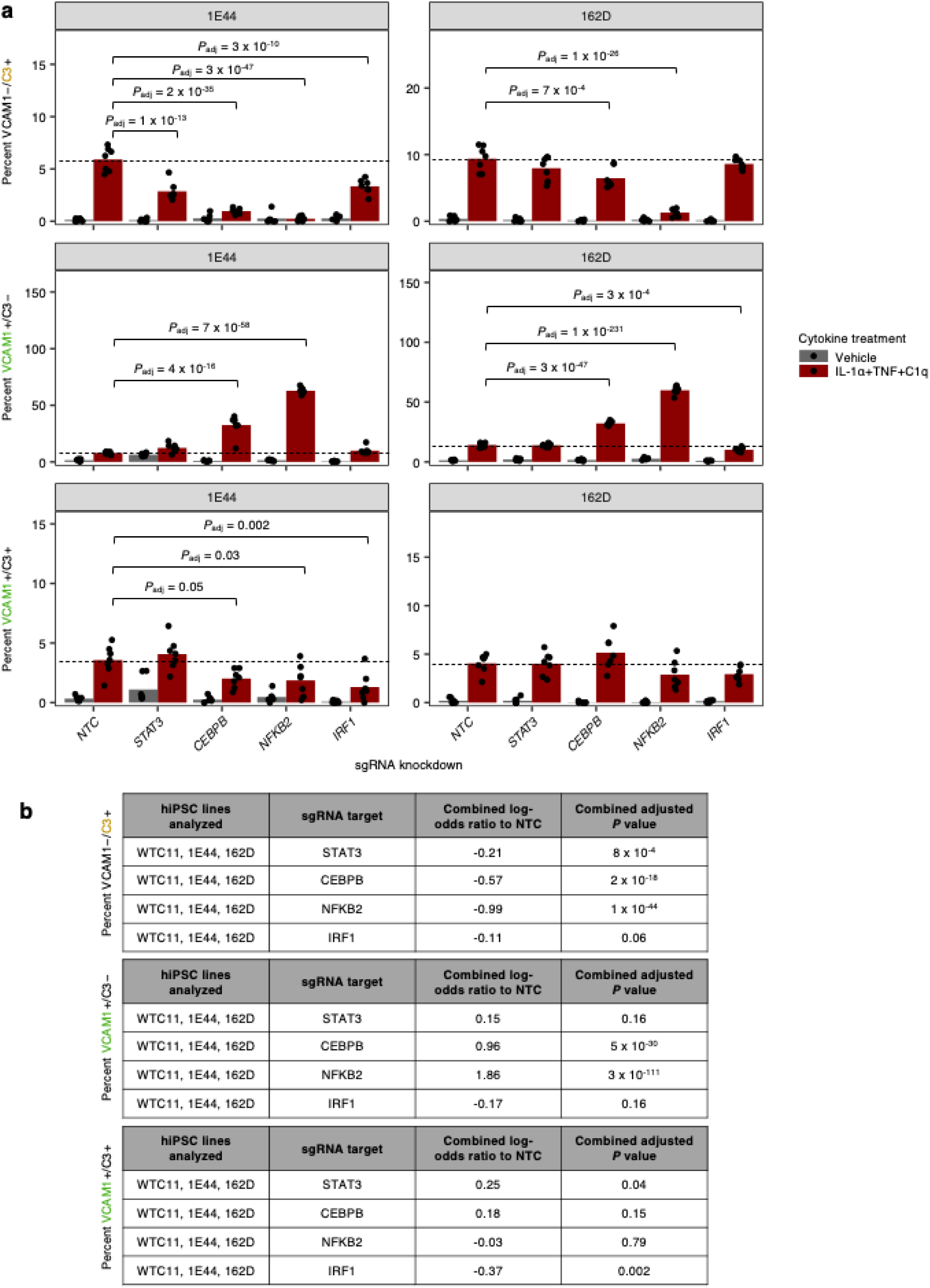
Validation of STAT3, CEBPB, NFKB3, and IRF1 knockdown in iAstrocytes derived from multiple hiPSC lines. **a**, Percent VCAM1−/C3+, VCAM1+/C3−, or VCAM1+/C3+ cells measured by flow cytometry)in iAstrocytes derived from multiple hiPSC lines (TCW-1E44 and 162D) transduced with non-targeting sgRNA (NTC) or sgRNAs targeting *STAT3*, *CEBPB*, *NFKB2*, or *IRF1* (*n* = 6 wells). **b**, Combined statistical analysis of the effect of *STAT3*, *CEBPB*, *NFKB2*, or *IRF1* knockdown compared to NTC in IL-1α+TNF+C1q-treated iAstrocytes derived from multiple hiPSC lines (WTC11, TCW-1E44, 162D). For panels **a** and **b**, *P* values were calculated using beta regression and adjusted for multiple testing (*P*_adj_; Holm’s method) per family of tests (all comparisons within a plot or table).

**Extended Data Fig. 9.**
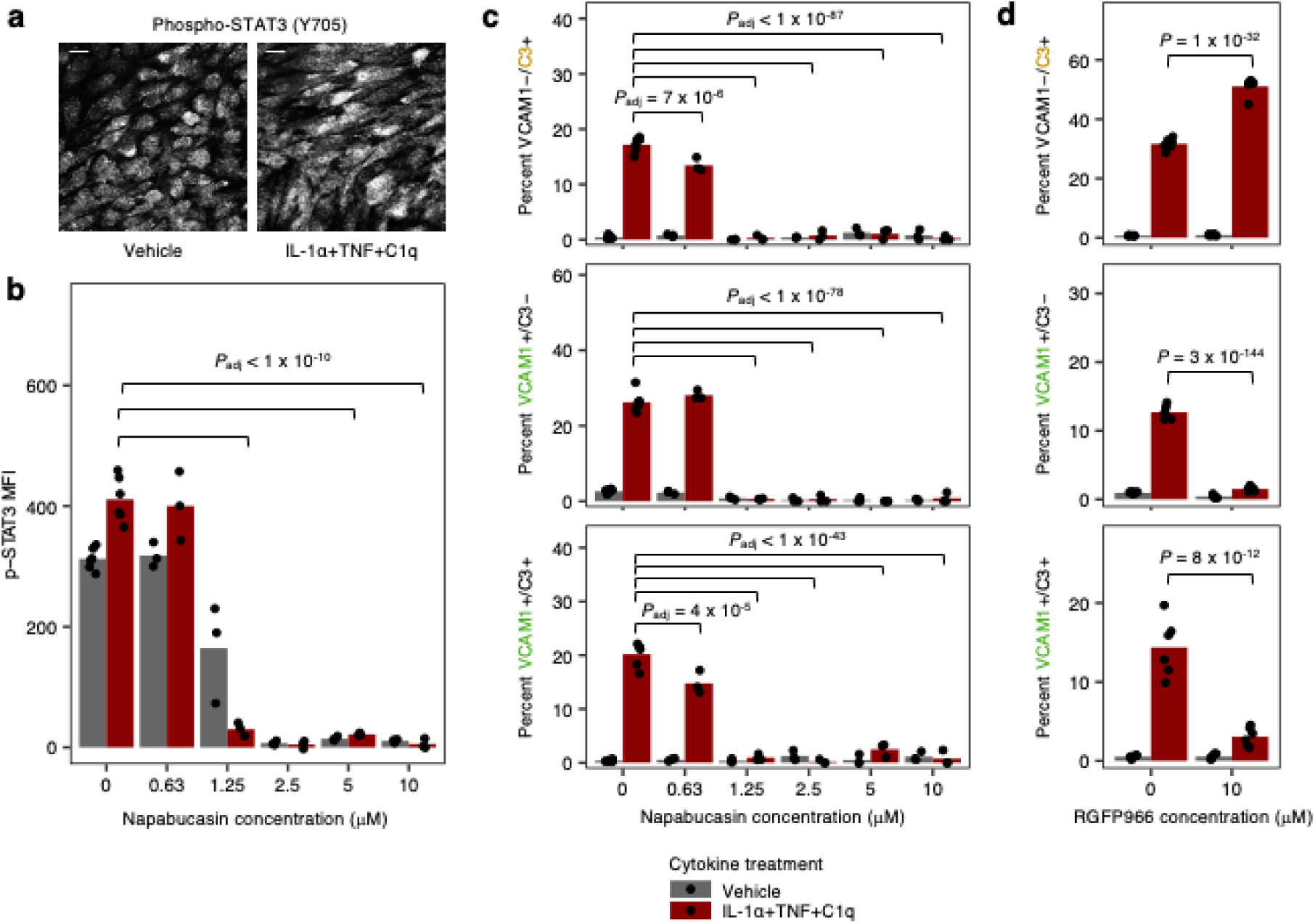
Effect of small molecule modulators of STAT3 or STAT1/2 activity. **a**, Representative immunofluorescence images of phospho-STAT3 (Y705) staining in vehicle control vs. IL-1α+TNF+C1q-treated iAstrocytes. **b**, Phospo-STAT3 (Y705) levels measured by flow cytometry in iAstrocytes treated with vehicle control vs. IL-1α+TNF+C1q in the presence of increasing doses of napabucasin (*n* = 6 wells for 0 μM napabucasin, *n* = 3 for napabucasin > 0 μM). MFI: median fluorescence intensity. **c**, Percent VCAM1−/C3+, VCAM1+/C3−, or VCAM1+/C3+ cells measured by flow cytometry)in iAstrocytes treated with vehicle control vs. IL-1α+TNF+C1q in the presence of increasing doses of napabucasin (*n* = 6 wells for 0 μM napabucasin, *n* = 3 for napabucasin > 0 μM). **d**, Percent VCAM1−/C3+, VCAM1+/C3−, or VCAM1+/C3+ cells measured by flow cytometry in iAstrocytes treated with vehicle control vs. IL-1α+TNF+C1q, with or without concurrent RGFP966 treatment (*n* = 6 wells). In panels **b**-**d**, *P* values were calculated using linear regression for MFI values or beta regression for percentages and adjusted for multiple testing (*P*_adj_; Holm’s method) per family of tests (all comparisons within a plot).

**Extended Data Fig. 10.**
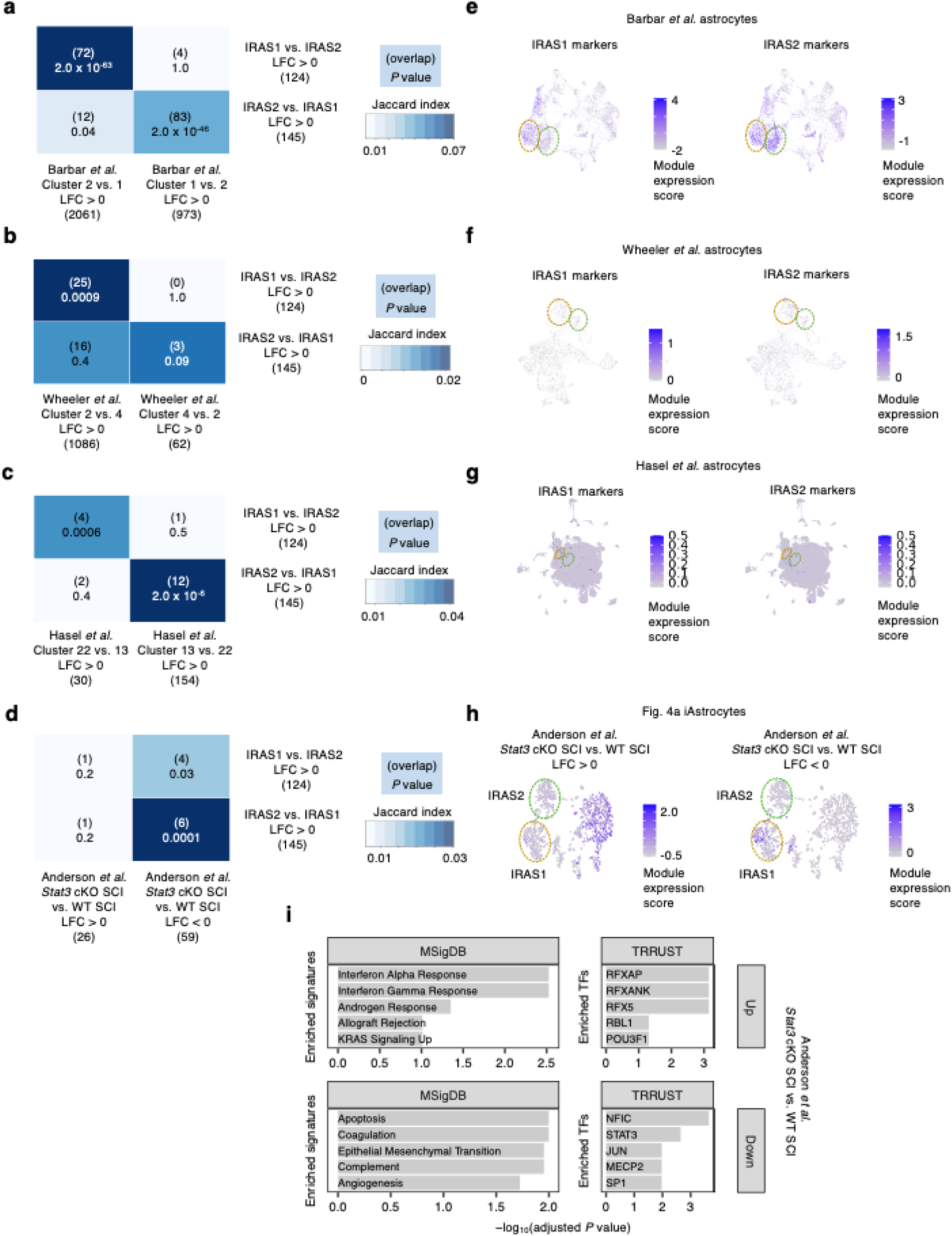
Overlap analysis of IRAS1 and IRAS2 markers with external datasets. **a**-**c**, Overlap analysis (Fisher’s exact test; see Methods) of differentially expressed genes (DEGs) between IRAS1 vs. IRAS2 with DEGs between IRAS1- and IRAS2-co-clustering astrocytes from Barbar *et al.*^29^ (**a**), Wheeler *et al.*^83^ (**b**), or Hasel *et al.*^58^ (**c**). **d**, Overlap analysis of DEGs between IRAS1 vs. IRAS2 with DEGs between astrocytes from *Stat3* astrocyte-specific conditional knockout (cKO) mice vs. wild-type (WT) mice subject to spinal cord injury (SCI) from Anderson *et al.*^51^. **e**-**g**, Module expression score (see Methods) of IRAS1 or IRAS2 markers overlaid onto the UMAP embedding of Barbar *et al.*^29^ (**e**), Wheeler *et al.*^83^ (**f**), or Hasel *et al.*^58^ (**g**) astrocytes from Fig. 7d, h, and l, respectively. **h**, Module expression score of upregulated vs. downregulated DEGs between astrocytes from *Stat3* cKO SCI vs. WT SCI mice from Anderson *et al.*^51^ overlaid onto the UMAP embedding of iAstrocytes from Fig. 4a. **i**, Cellular pathway (MSigDB^131^) and upstream transcription factor (TRRUST^132^) enrichment analysis of upregulated vs. downregulated DEGs between astrocytes from *Stat3* cKO SCI vs. WT SCI mice from Anderson *et al.*^51^.

## SUPPLEMENTARY TABLE LEGENDS

**Supplementary Table 1 | Bulk RNA-seq differentially gene expression analysis results.** From this study: iAstrocytes and hiPSC-derived astrocytes generated using the protocols from TCW *et al.*^18^, Li *et al.*^19^, and Krencik *et al.*^15^ treated with vehicle vs. IL-1α+TNF+C1q. From external datasets: immunopanned primary mouse astrocytes treated with vehicle vs. IL-1α+TNF+C1q from Guttenplan *et al.*^9^, hiPSC-derived astrocytes treated with vehicle vs. IL-1β from Perriot *et al.*^122^, and human cerebral organoid-derived astrocytes treated with vehicle vs. IL-1α+TNF+C1q from Barbar *et al.*^29^. For data from hiPSC-derived astrocytes in this study, expression was quantified using Salmon and differentially expressed genes (DEGs) called using DESeq2. For external datasets, the DEGs from Gutteplan *et al.* were downloaded from the supplementary table included in GSE143598, and raw data from Perriot *et al.* and Barbar *et al.* were analyzed with BioJupies^121^ to call DEGs.

**Supplementary Table 2 | Phenotype scores and p-values from CRISPRi screens.** See the first tab of the Excel file for a description of the contents of the remaining tabs and information regarding the columns in each tab.

**Supplementary Table 3 | Hits from MRA and associated scores and statistics.** See Castro *et al.*^38^ for interpretation of the activity score. The mean LFC score of a regulon is calculated by averaging the log_2_-fold-change of all differentially expressed genes induced by IL-1α+TNF+C1q in that regulon.

**Supplementary Table 4 | Cluster markers of iAstrocytes in** Fig. 4a and dif**ferential expression analysis between IRAS1 and IRAS2 iAstrocytes.** The column “avg_diff” contains the magnitude of the difference in the mean Pearson residual values of a given gene between cells in the cluster of interest vs. all other cells, or between IRAS2 (cluster 2) vs. IRAS1 (cluster 1) iAstrocytes. The columns “pct.1” and “pct.2” contain the percent of cells expressing the gene of interest in the cluster of interest (“pct.1”) or all other cells (“pct.2”), or in IRAS2 (“pct.1”) iAstrocytes or IRAS1 (“pct.2”) iAstrocytes.

**Supplementary Table 5 | Enrichment analysis of IRAS1 and IRAS2 markers.** Enrichment results for the MSigDB^131^ and TRRUST^132^ libraries were downloaded from Enrichr^123^. See Chen *et al.*^123^ for how the “combined score” is calculated.

**Supplementary Table 6 | Differential gene expression analysis results from knockdown of regulators in CROP-seq experiment.** The results shown correspond to the interaction term between cytokine treatment and regulator knockdown, i.e. they reflect how regulator knockdown affects the change in gene expression induced by IL-1α+TNF+C1q. See the first tab of the Excel file for the number of cells recovered for each knockdown.

**Supplementary Table 7 | Gene modules and enrichment analysis of gene modules from Extended Data Fig. 4d.** Enrichment results for the MSigDB^131^ and TRRUST^132^ libraries were downloaded from Enrichr^123^. See Chen *et al.*^123^ for how the “combined score” is calculated.

**Supplementary Table 8 | Cluster markers and DEGs from integrated analysis of iAstrocytes with Barbar *et al.*, Wheeler *et al.*, or Hasel *et al.* astrocytes.** Cluster markers and DEGs are derived using only astrocytes from Barbar *et al.*^29^, Wheeler *et al.*^83^, or Hasel *et al.*^58^. The column “avg_diff” contains the magnitude of the difference in the mean Pearson residual values of a given gene between cells in the cluster of interest vs. all other cells, or between two clusters of interest (the first cluster minus the second cluster). The columns “pct.1” and “pct.2” contain the percent of cells expressing the gene of interest in the cluster of interest (“pct.1”) or all other cells (“pct.2”), or the two clusters of interest.

**Supplementary Table 9 | Differential gene expression analysis between Stat3 cKO SCI vs. WT SCI astrocyte transcriptomes from Anderson *et al.* and enrichment analysis of DEGs.** The raw data from Anderson *et al.*^51^ was analyzed with BioJupies^121^ to call DEGs between Stat3 cKO SCI vs. WT SCI astrocyte transcriptomes. Enrichment results for the MSigDB^131^ and TRRUST^132^ libraries were downloaded from Enrichr^123^. See Chen *et al.*^123^ for how the “combined score” is calculated.

**Supplementary Table 10 | Clinical metadata of human neuropathology samples.** For the AD cohort, Braak stage is provided for each sample. Additional neuropathology scoring (e.g. Thal and CERAD scores) are provided if available.

**Supplementary Table 11 | Metadata of astrocyte bulk RNA-seq samples used for co-expression network reconstruction in master regulator analysis.** Samples from publicly available datasets can be cross-referenced with GEO using their GSM accession numbers.

**Supplementary Table 12 | sgRNA information for CRISPRi libraries.** sgRNA_ID: A unique ID for each sgRNA that includes the gene target, orientation, genomic coordinate, and the TSS targeted. Gene_target: the gene targeted by the sgRNA. TSS: the transcriptional start site targeted by the sgRNA. sgRNA_sequence: the protospacer sequence of the sgRNA.

**Supplementary Table 13 | Information on mice used for immunostaining.**

