## Supplementary Figure 1 for "CRISPRi screens in human astrocytes elucidate regulators of distinct inflammatory reactive states"

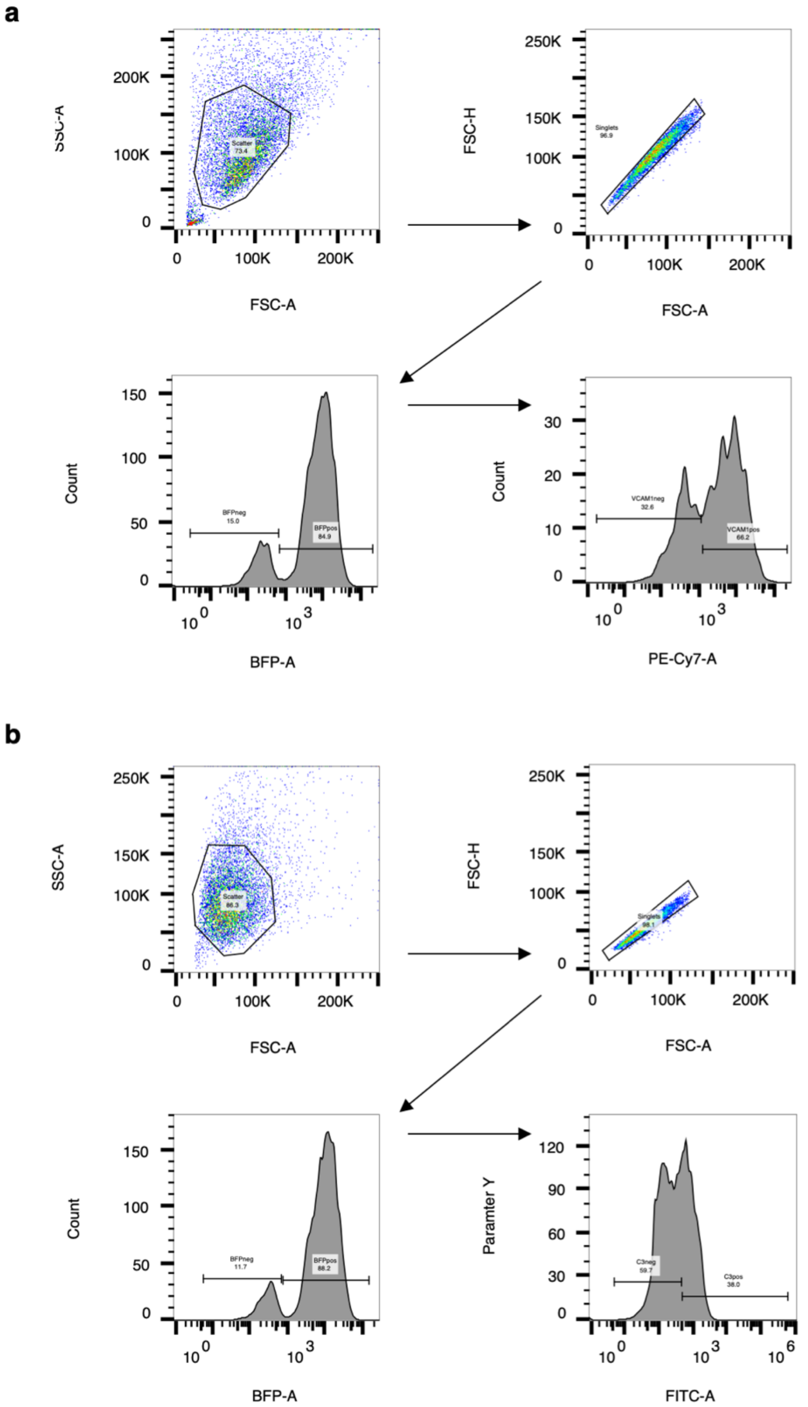


**Supplementary Fig. 1 | Flow cytometry gating strategy. a-b**, Gating strategy for VCAM1 staining (**a**, live cells) or C3 (**b**, fixed and permeabilized cells) staining. Cells were separated from debris using the FSC-A vs. SSC-A gate, single cells were selected using the FSC-A vs. FSC-H gate, BFP+ cells (sgRNA-transduced cells) were then selected using the BFP-A gate (applicable only to experiments where astrocytes were transduced with sgRNA lentivirus), and then cells were gated as VCAM1+ vs. VCAM1- using the PE-Cy7-A gate or C3+ vs. C3- using the FITC-A gate. The same gating strategy also applies to phagocytosis experiments and TFRC staining. For the FACS-based VCAM1 and phagocytosis screens, the same gating strategy applies, except that cells are sorted in to a low bin (bottom 30% of events from parent gate) or high bin (top 30% of events from parent bin) based on pHrodo or VCAM1 fluorescence intensity.
